# Separation of halide oxidation and substrate halogenation chemistries rationalizes site-selective vanadium dependent haloperoxidase catalysis

**DOI:** 10.1101/2025.10.24.684477

**Authors:** Jackson T. Baumgartner, Lukas A. Varga, Jackson T. Calhoun, Vitor H. Balasco Serrão, Sarah Loerch, Shaun M. K. McKinnie

## Abstract

Vanadium-dependent haloperoxidases (VHPOs) catalyze the halogenation of organic molecules under mild aqueous conditions. Selective bacterial VHPOs exhibit exquisite regio- and enantiocontrol, however the precise mechanisms dictating selectivity have remained elusive. We have solved the single-particle cryo-electron microscopy (cryo-EM) structure of a selective bromoperoxidase from *Enhygromyxa salina* (esVHPO). Mutagenesis demonstrates that halide oxidation and substrate halogenation occur in two distinct pockets, with halide transfer mediated by critical lysine residue K329. Isolation of a stable intermediate following bromide oxidation (BrOx) enables single turnover catalysis in the presence of organic substrate; subsequent application of a chemoselective fluorescent probe provides support for an intermediate bromamine involved in selectivity. Cryo-EM of the BrOx state reveals a ‘camera shutter’ mechanism that compacts the halide entry tunnel and vanadate pocket, minimizing the premature dissociation of hypohalous acid. These findings collectively unveil a multilayered halogen trapping and transfer mechanism and provide a rationale for selective VHPO catalysis.

**Graphical abstract:** 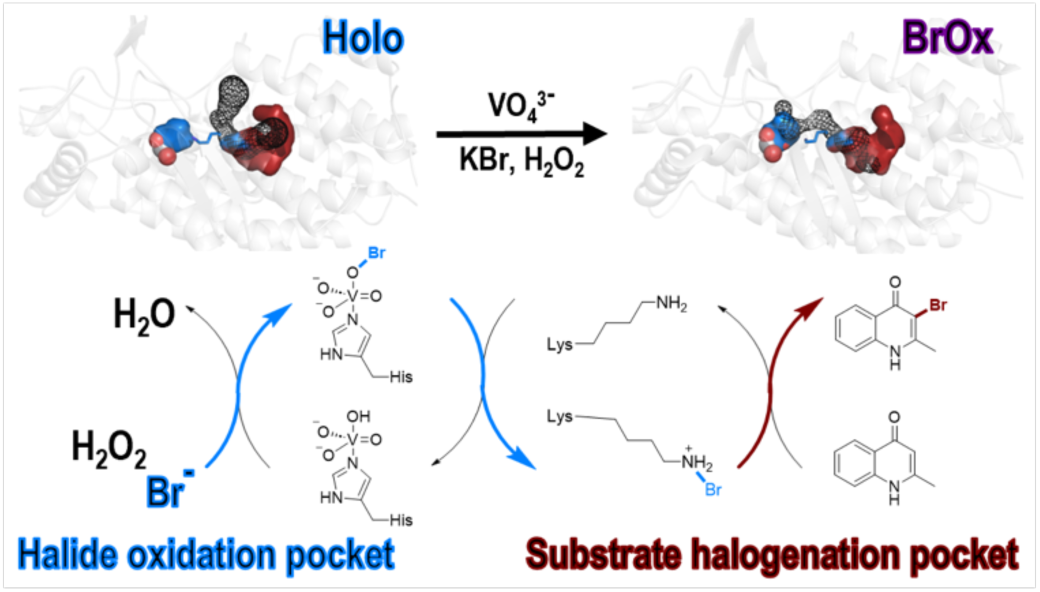

## Introduction

Halogens are some of the most synthetically useful and therapeutically relevant functional groups in organic chemistry.^1^ Their inclusion enhances potency, cell permeability, and pharmacokinetics; this impact on activity is reflected in the majority of our most important drugs containing halogens.^2^ Traditional halogenation requires harsh conditions, toxic reagents, or expensive catalysts, while selectivity remains a persistent challenge. In contrast, Nature has evolved halogenases that catalyze similar reactions under mild aqueous conditions with high regio- and stereoselectivity.^3^ Recent advances in enzyme discovery and engineering have positioned biocatalysis as a powerful tool for synthesis and the distinct catalytic strategies of halogenases provide multiple avenues to improve synthetic route design.^4–6^ While nucleophilic and radical halogenases have been heavily explored for biocatalysis,^7–9^ electrophilic halogenases display high biocatalytic potential.^10^ Flavin-dependent halogenases (FDH) are the best studied electrophilic family; mechanistically, FDHs physically separate halide oxidation from substrate binding and connect the sites *via* hydrophobic tunnel and catalytic lysine. While FDHs have been widely studied and engineered, vanadium-dependent haloperoxidases (VHPO) remain underutilized.

VHPOs uniquely use a histidine-coordinated orthovanadate and hydrogen peroxide to oxidize halide ions (Cl^-^, Br^-^, I^-^) into hypohalous acid (HOX).^11,12^ Depending on the most electronegative halide oxidized, they are designated as chloroperoxidases (VCPO), bromoperoxidases (VBPO), or iodoperoxidases (VIPO). Fungal and algal VHPOs feature solvent-exposed vanadate sites that generate freely diffusible HOX, leading to spontaneous halogenation in solvent (**Fig. 1A**). Though specific substrate preferences have been exhibited, uncontrolled HOX diffusion occurs in the absence of acceptor substrates.^13–15^ Despite this, VHPOs offer appealing biocatalytic traits, like temperature and solvent tolerance, redox-neutral catalysis, and independence from auxiliary co-factors or proteins.^16,17^ These properties have enabled efforts to harness *in situ* HOX production under mild conditions (**Extended Data Fig. 1A**).^18^

**Figure 1.**
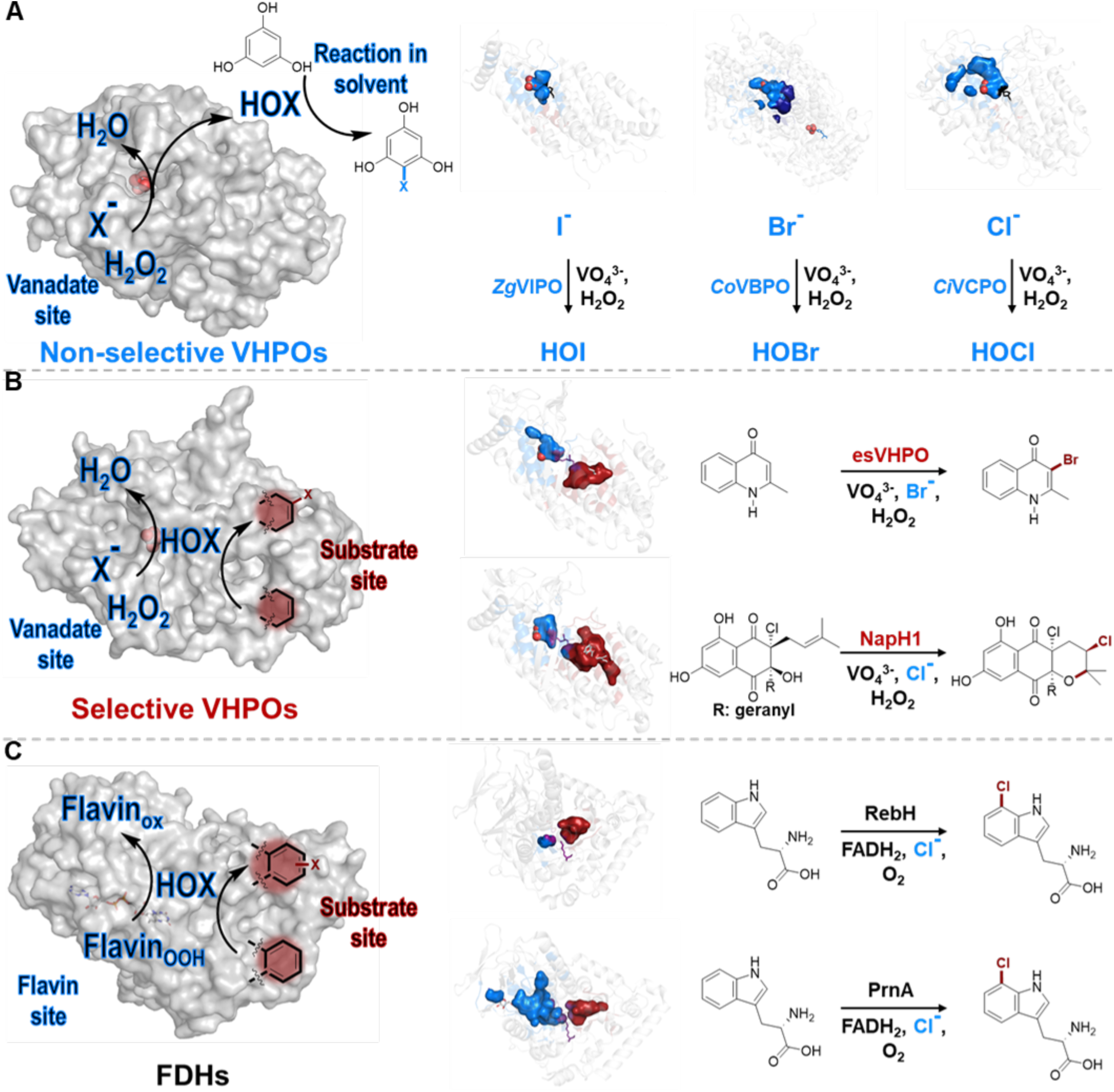
Chemistry and structure of electrophilic halogenases. The halide oxidative co-factor sites for VHPOs (vanadate site) and FDHs (flavin site) are highlighted in blue while the substrate binding sites and selective halogenation chemistries are highlighted in red. **A)** Reactions catalyzed by non-selective VHPOs. *Ci*VCPO (PDBID 1VN1) is shown in grey surface representation with vanadate as spheres. Subunits with internal cavity structures are shown in grey ribbons with vanadate as spheres and proline substitutions of catalytic lysine highlighted in black. Structures and chemistries for *Zg*VIPO (PDBID 4USZ), *Co*VBPO (PDBID 7QWI), and *Ci*VCPO (PDBID 1VN1) are shown. **B)** Reactions catalyzed by selective VHPOs. NapH1 (PDB ID 3W36) with internal cavities are represented as in A, with the catalytic lysine in purple. Structures and chemistries for esVHPO (AlphaFold 2.0 model) and NapH1 (PDBID 3W36) are shown. **C)** Reactions catalyzed by FDHs. PrnA (PDB ID: 2AQJ) with flavin shown as white sticks and internal cavities are represented as in B. Structures and chemistry for RebH (PDBID 2E4G) and PrnA (PDBID 2AQJ) are shown.

Bacterial biosynthetic VHPOs offer regio- and enantioselective halogenation with high substrate selectivity.^19^ While certain algal VHPOs have demonstrated low yielding enantioselective terpene cyclization,^13^ the Streptomyces-derived NapH1 marked the first discovery of a dedicated biosynthetic VHPO.^20^ Other bacterial VHPOs selectively halogenate naphthoquinone, phenazine, and alkyl quinolone (AQ) scaffolds (**Extended Data Fig. 1B**).^21–23^ Selective VHPOs mediate complex rearrangements and exhibit broader activity scopes than initially thought. Notably, selective VHPOs do not release HOX, suggesting a distinct and controlled mechanism of halide oxidation and halogenation.^19^ This mechanism remains unresolved, limiting rational design and engineering efforts. Structural comparison reveals that while non-selective VHPOs possess solvent-exposed vanadate sites (**Fig. 1A**), selective VHPOs exhibit buried vanadate sites which access a defined putative substrate pocket (**Fig. 1B**); an architecture reminiscent of the FDHs (**Fig. 1C**). Additionally, the conserved lysine and hydrophobic tunnel found in FDHs are also present in selective VHPOs.^24,25^ We hypothesize that selective VHPOs analogously separate halide oxidation and substrate halogenation, mediated through a conserved lysine. This hypothesis is supported by structural and mutagenesis studies of NapH1.^26^

Here we investigate a selective VHPO from *Enhygromyxa salina* (esVHPO), which catalyzes regioselective AQ bromination.^23^ Using cryogenic electron microscopy (cryo-EM), mutagenesis, chemical probes, and kinetics, we demonstrate that selective VHPOs achieve substrate selectivity by physically separating vanadate-catalyzed halide oxidation from substrate halogenation. HOX control is exerted by a lysine-bromamine intermediate and conformational gating in the vanadate site. This mechanism enables precise halogenation and supports VHPO engineering for broader biocatalytic applications.

## Results

### Cryo-EM structure demonstrates conservation of core features in esVHPO

Our hypothesis that selective VHPOs use a structural paradigm distinct from non-selective VHPOs was predicated on limited available structural data, primarily the crystal structure of NapH1 (PDBID 3W35 and 3W36). To extend this model, we determined the structure of esVHPO using single-particle cryo-EM to 2.6 Å overall resolution (PDBID 9PTR; EMD-71844) (**Extended Data Figs. 2, 3**). The holo-esVHPO reconstruction revealed a dimer with C2 symmetry (**Fig. 2A, B**), consistent with size exclusion chromatography (SEC). Residues 70-528 (**Fig. 2A, B**) are resolved, while 69 N-terminal residues, partially corresponding to a secretion signal, are disordered. The structure comprises an N-terminal helical bundle (NHB, residues 76-205), an interconnecting region (ICR, residues 206-288), and a C-terminal helical bundle (CHB, residues 289-521). Each subunit shows clear density for one orthovanadate coordinated to H496 and stabilized by the sidechains of K373, R380, R490, S427, and S429 (**Fig. 2C**). Backbone interactions with A428 and S429, on the N-terminal cap of helix 12, likely stabilize the helix and modulate the electronic properties of vanadate.^27^ The vanadate geometry adopts previously observed trigonal pyramidal geometry, contrasting with recent reports of a planar metavandate.^15,26,28–30^ esVHPO shares a highly similar fold and quaternary structure with NapH1 (1.7 Å^2^ rmsd, 448 C_alphas_) and contains the proposed haloamine forming Lys-Ser dyad (**Fig. 2D**). Distinct conserved waters in the vanadate site are also observed. The ICR contributes to dimerization through a conserved handshake-like interaction where F238 inserts into a hydrophobic pocket formed by a beta hairpin (residues 250-260) lined by the aliphatic face of T236, P233, P248, and H262 of the other subunit (**Fig. 2E**). A similarly conserved interface forms a β-sheet between the subunits (**Fig. 2F**). Analogous to FDHs, esVHPO and NapH1 position hydrophobic residues between the vanadate site and putative substrate site (**Supplementary Fig. S4**). Despite their similarities, esVHPO and NapH1 differ in a few key areas. The beta-hairpin of the ICR (residues 241-254 in NapH1 and residues 253-256 in esVHPO) that hangs over the entry to vanadate site is conspicuously larger in NapH1. This distinction is consistent across VBPO and VCPO sequence alignments, however the functional role of this difference is unclear (**Fig. 2D**). Another notable difference is a four residue insertion in NapH1 (A74-S77) that disrupts the N-terminal helix lining the putative substrate pocket (**Fig. 2D**); this insertion is shared with the non-halogenating NapH3 that modifies a similar substrate to NapH1.^26^

**Figure 2.**
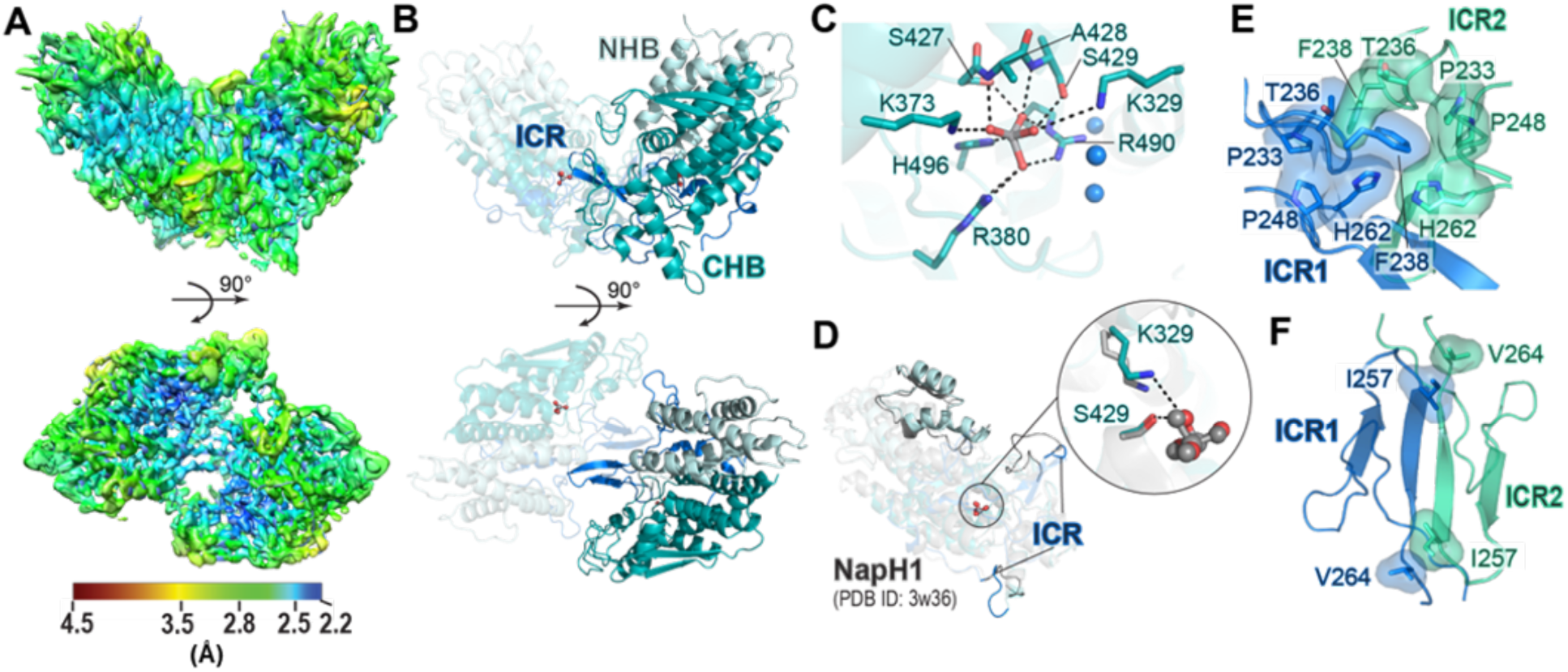
Architecture of holo-esVHPO. **A)** Local resolution map of the esVHPO dimer (PDBID 9PTR). **B)** The overall structure of dimeric holo-esVHPO. The N-terminal helical bundle (NHB) is shown in pale cyan, the interconnecting region (ICR) in marine, and the C-terminal helical bundle (CHB) in teal. **C)** Closeup of the vanadate coordinating residues and conserved waters. **D)** A comparison of holo-esVHPO with closest structural homolog NapH1 (PDBID 3W36) from *Streptomyces*. Divergent regions are highlighted in opaque. Only one subunit is shown for clarity. The structural conservation of the K329-S429 dyad in holo-esVHPO are shown as teal sticks in the insert. The equivalent K324-S427 dyad in NapH1 is shown as grey sticks. **E)** Reciprocal tongue-in-groove interactions between the interconnecting domains at the dimer interface. **F)** Second hydrophobic interacting regions between the two monomers.

### Steady-state kinetics of esVHPO WT

For biophysical characterization, dimeric esVHPO wildtype (WT) was purified by SEC and its activity was confirmed consistent with prior reports.^23^ Non-selective macroalgal VHPOs follow a bi-bi ping-pong mechanism, where hydrogen peroxide first reacts with vanadate followed by halide oxidation as the rate limiting step.^12^ We expected esVHPO to follow a similar mechanism and performed steady-state kinetics by monitoring Br-MHQ formation while varying individual substrates (**Extended Data Fig. 4**). Bromide oxidation was rate-limiting and displayed standard Michaelis-Menten kinetics with a K_0.5_ of ∼1.0 mM, comparable to seawater bromide concentrations. In contrast, the Hill equation provided the best fit for AQ and hydrogen peroxide, indicating a positive cooperativity not observed in non-selective VHPOs or FDHs.^31,32^ Bromide showed no cooperativity, suggesting the source of cooperativity is independent from bromide binding. A Hill coefficient of 2.0 for MHQ implies either a conformational requirement for binding or communication between multiple substrate binding sites. To probe the bromination step, we synthesized a C3-deuterated MHQ substrate which yielded a secondary KIE, supporting a minor rate contribution from C3-deprotonation after bromination. This aligns with the rate contribution of Wheland-intermediate proton abstraction observed in the FDHs.^33^

### Identification of the putative substrate binding site

To identify the AQ binding site, we aligned sequences of characterized selective VHPOs and mapped conservation onto holo-esVHPO using ConSurf.^34^ Selective VHPOs form a distinct phylogenetic clade, separated from other bacterial VHPOs and further sub-divided along substrate specificity. (**Fig. 3A**). In AQ-VHPOs, the vanadate site and adjacent tunnel residues are highly conserved, consistent with their isofunctionality (**Fig. 3B**). Notably, a conserved pocket adjacent to the tunnel residues suggested a putative substrate site, with subtle variations rationalizing differences in substrate specificity across homologs.^23^ Expanding the same workflow to include actinobacterial meroterpenoid biosynthetic VHPOs showed vanadate and tunnel conservation, but dramatically reduced conservation in the substrate site (**Fig. 3C**). To validate the pocket as the substrate site, we docked HHQ into holo-esVHPO. Initial docking was restricted to the vanadate site and adjacent pocket of one subunit. The top 20 lowest-energy poses localized within the putative substrate pocket, with the quinolone ring oriented toward the hydrophobic tunnel (**Extended Data Figure 5A, B**). Expanded docking was later performed on the entire dimer with recently available server-based software which expanded our search space to the entire dimeric unit (**Extended Data Fig. 5C**).^35^ This resulted in an even distribution of poses between subunits, recapitulated AutoDock orientations, and the absence of surface or dimer-interface binding, indicating a unique mode of substrate binding compared to non-selective *Am*VHPO and *Ci*VCPO.^14,15^

**Figure 3.**
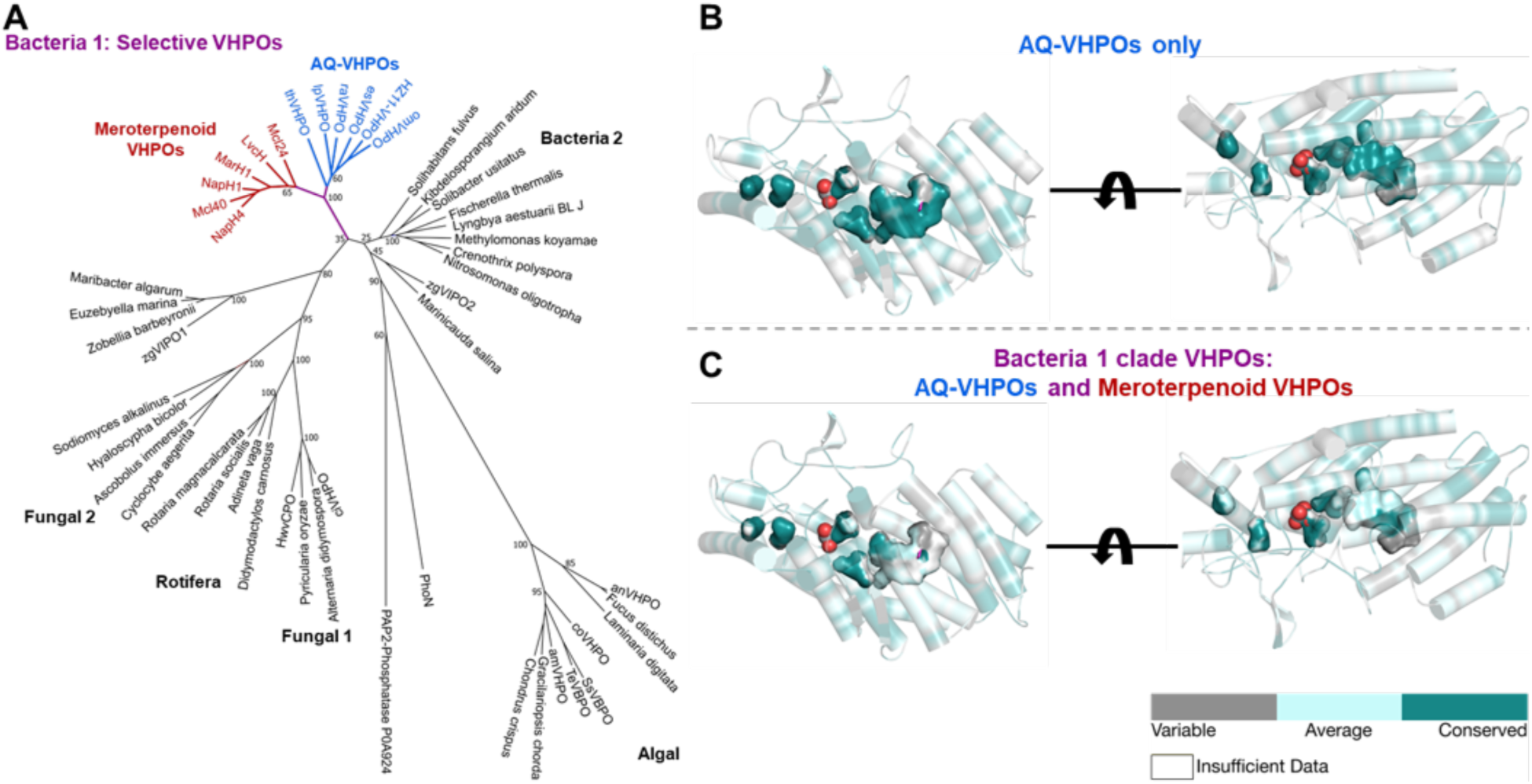
Selective VHPO phylogenetics and conservation. A) All kingdom phylogenetic tree of VHPOs. Non-selective VHPO clades are in black, the selective VHPO branch is in purple, meroterpenoid VHPOs are in red, and AQ-VHPOs are in blue. Two Type-II phosphatases are included as roots. Bootstrap values are shown for major splits. Clade labels are based on previous designations.^67^ B) Sequence alignment of characterized AQ-VHPOs were mapped onto the structure of holo-esVHPO using the ConSurf server.^34^ Vanadate is shown as spheres, the secondary structure as ribbons and cylindrical helices, and internal cavities as surface. Conservation scores were determined by ConSurf and coloration represents consolidated conservation scores. C) As panel B, however a sequence alignment including both AQ-VHPOs and meroterpenoid VHPOs was used instead.

### Mutagenesis reveals distinct substrate binding-site

We selected fourteen residues for site-directed mutagenesis to probe their roles in catalysis (**Fig. 4A-D**). These included three vanadate pocket residues (H496, E327, S429), three lining the putative tunnel (K329, V102, L333), and eight forming the predicted substrate pocket (H436, I330, G334, T106, T358, N359, D355, S110). Alanine mutations were prioritized with additional substitutions introduced as needed. Each variant was evaluated by analytical SEC to assess solubility and aggregation. Minor differences in oligomerization were observed, however higher-order oligomers were shown to have little impact on bromination (**Supplementary Figs. S7, S8**). Functional activity was assessed using two orthogonal assays: Ultra-high performance liquid chromatography mass spectrometry (UPLC-MS) to quantify bromination of five AQ substrates (**Fig. 4B**) and a colorimetric thymol blue assay to track diffusible HOI production (**Fig. 4C).** Together these assays distinguish residues involved in substrate bromination or halide oxidation, respectively (**Fig. 4D**).

**Figure 4.**
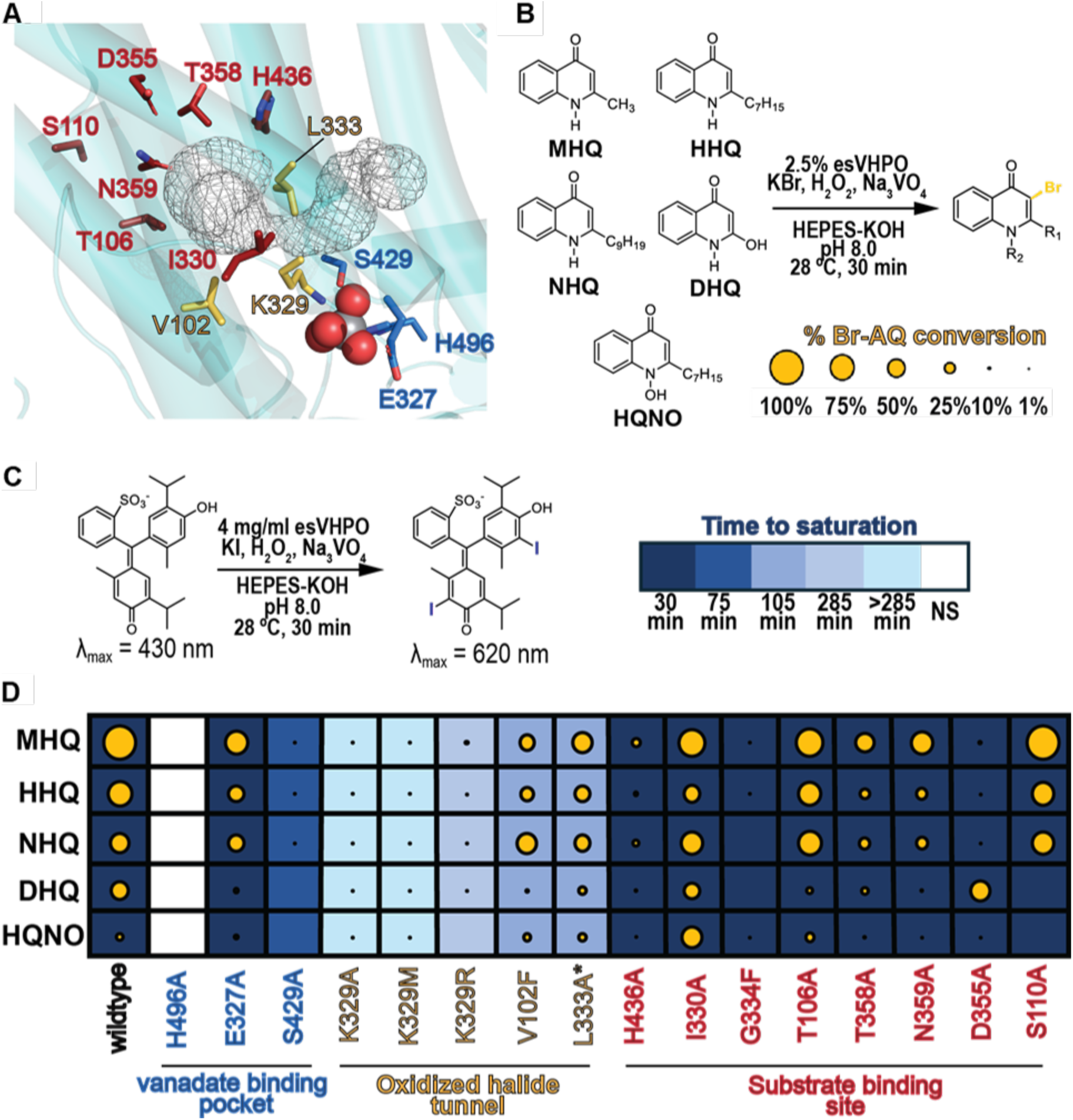
esVHPO mutagenesis supports separation of halide oxidation and substrate bromination. **A)** Structural localization of mutated residues in this study. Holo-esVHPO is in teal, vanadate is shown as spheres, and a CAVER tunnel is shown in black mesh. Vanadate site residues are in blue, tunnel residues are in yellow, and substrate site residues are in red. **B)** General reaction scheme for esVHPO variant substrate profiling. The legend for extent of substrate bromination in panel D is also shown. **C)** Thymol blue assay for esVHPO variant iodide oxidation and diffusion. A scale for the amount of time to achieve complete iodination of the thymol blue reporter used in panel D is also shown. **D)** Activity panel for all prepared esVHPO variants. Each column represents the activity of a specific variant. Rows represent the bromination activity of a variant on the substrate profile. Blue shading represents the diffusible HOI production of each variant. Residues are listed in order of distance from vanadate rather than chain position. L333A exhibited high soluble aggregation so activity measurements may not directly reflect its effect on turnover. Conversion percentages, significance scores, and thymol blue time course assays for each variant are shown in **Supplementary Figs. S9, S10**.

Controls with native and boiled esVHPO WT established a baseline for non-enzymatic iodide oxidation under assay conditions.^36^ Mutation of the vanadate-binding H496 abolished HOI production and AQ bromination. E327A and S429A retained iodide oxidation, but only E327A retained partial substrate bromination activity. S429A failed to produce brominated AQs, implying a role specific to bromination. Removal of the K329 positive charge (K329A and K329M) drastically reduced iodide oxidation and substrate bromination; while restoration of the charge (K329R) partially rescued HOI production but maintained a comparable loss in AQ bromination. Consistent with its proposed role in NapH1 and FDHs, the positive charge and free amine of K329 are required for efficient catalysis. Other tunnel mutants (V102F and L333A) modestly reduced overall activity likely due to catalytic or structural disruptions.

The most striking results arose from mutating the substrate pocket (**Fig. 4D**). All eight variants exhibited wild-type levels of HOI production, suggesting they do not disrupt halide oxidation. However, AQ bromination profiles varied widely. T106A and S110A mirrored wild-type activity, consistent with their distal positions from vanadate (15-20 Å) and supporting the docking result that substrate binds adjacent to the tunnel. T358A and N359A caused moderate reductions, suggesting roles in substrate organization or hydrogen bonding. I330A maintained bromination on most substrates but showed enhanced activity on HQNO potentially due to expanded pocket size (**Extended Data Fig. 6A**). In contrast, H436A and G334F, nearly abolished bromination across all substrates. Despite its distance from docked HHQ, H436 may contribute to pocket shaping by second shell interactions (**Extended Data Fig. 6B**). G334, located at the entrance of the pocket, is highly conserved among selective VHPO homologs. G334F ablates AQ bromination activity, likely through restricting access to or shrinking the pocket (**Extended Data Fig. 6C**). Lastly, D355A abolished bromination of all C2-alkylated AQs but retained activity on the C2-hydroxylated DHQ (**Extended Data Fig. 6D**); this activity suggests that D355 does not catalyze deprotonation of the AQ cationic intermediate but rather mediates substrate binding. D355 is situated 4.4 Å from N1 of docked HHQ and its strong conservation in AQ-VHPOs suggests a role in adjusting substrate nucleophilicity. Electrostatic modeling revealed that D355A inverts the negatively charged substrate pocket to slightly positive, improving compatibility with mildly acidic DHQ (**Extended Data Fig. 6E**). These cumulative results reveal the functional separation of halide oxidation and bromination in esVHPO and generate strong support for the existence of a separate substrate binding pocket.

### A long-lived oxidized bromide intermediate is facilitated by K329

To probe mechanistic parallels between esVHPO and FDHs, we tested if esVHPO forms a stable oxidized halide intermediate, analogous to the long-lived intermediate reported for FDH RebH.^24^ esVHPO was incubated with vanadate, H_2_O_2_, and KBr to generate a bromide-oxidized intermediate (BrOx-esVHPO), which was subsequently purified into halide-free conditions using SEC (**Fig. 5A**). Incubation of BrOx-esVHPO with MHQ led to near stoichiometric Br-MHQ formation as determined through UPLC-MS (**Fig. 5B**). This intermediate exhibited temperature-dependent stability with a 5-hour half-life when incubated on ice (**Fig. 5C**). These data support that the esVHPO bromination reaction proceeds through a sequestered, stable, and long-lived intermediate.

**Figure 5.**
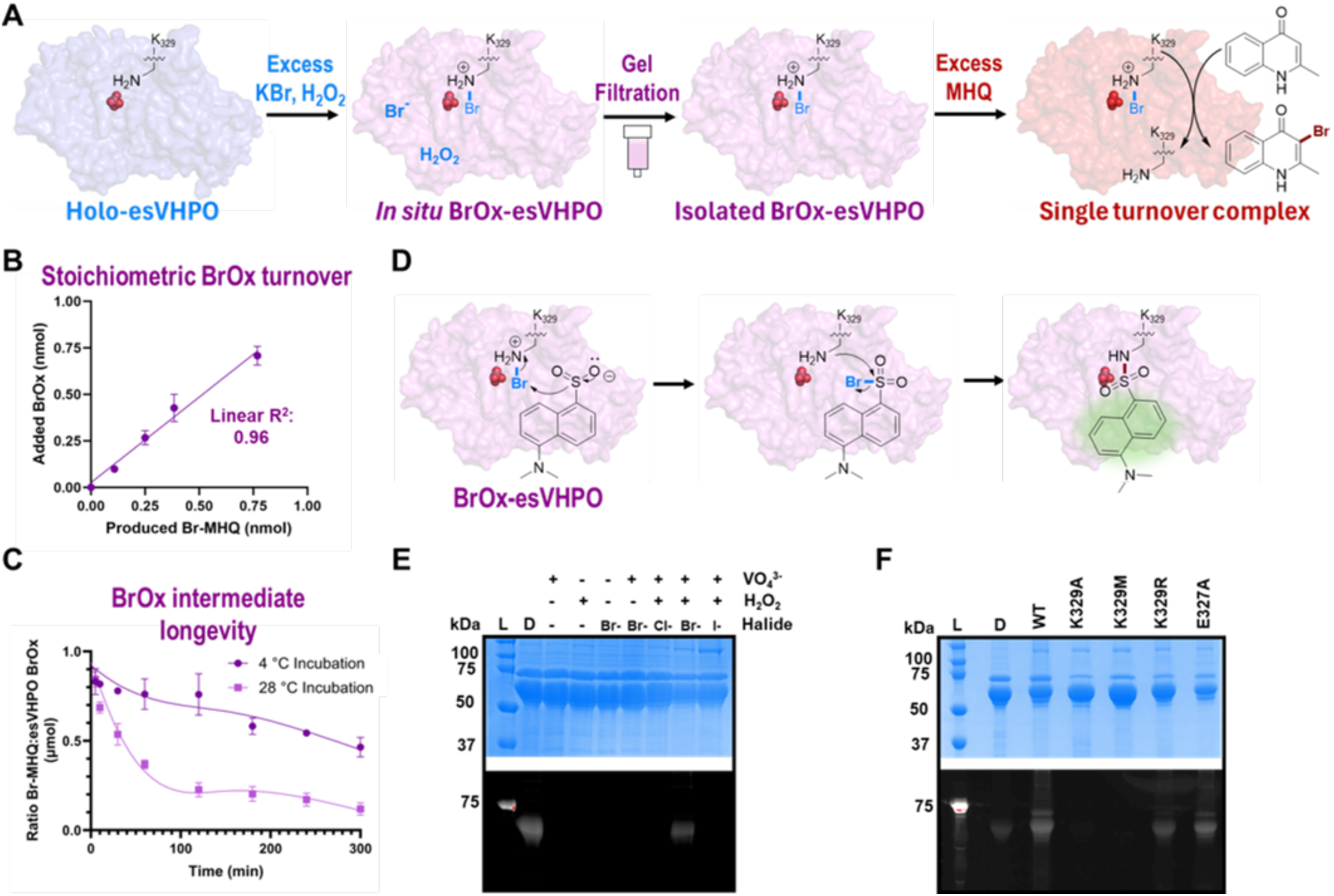
BrOx-esVHPO single turnover catalysis and haloamine chemistry. A) Isolation and preparation of the BrOx-esVHPO single turnover reaction. The cryo-EM models of holo-esVHPO and BrOx-esVHPO are shown as surface. B) Stoichiometric turnover of isolated BrOx-esVHPO with MHQ. The total product (Br-MHQ) produced in the single turnover assay in nmol is plotted against the amount in nmol of isolated BrOx-esVHPO added to reaction. Error bars represent the mean ± s.d. (n = 3). C) BrOx-esVHPO stability at different temperatures. The y-axis is plotted as the ratio of Br-MHQ (µmol) produced to the moles of isolated BrOx-esVHPO (µmol) added to the reaction. Error bars represent the mean ± s.d. (n = 3). D) Proposed reaction mechanism of BrOx-esVHPO haloamine with dansyl sulfinic acid (DANSO_2_H). Labeling only occurs in the presence of haloamine functionality. E) Substrate dependency of esVHPO labeling by DANSO_2_H, visualized via SDS-PAGE. Dansyl chloride (D) was used as a positive control and protein ladder (L) guided visualization. Gels were fluorescently imaged prior to staining with Coomassie blue. F) Reactions of esVHPO variants were prepared and visualized as panel E. Each reaction was performed in BrOx conditions.

In FDHs, a conserved active-site lysine directs substrate halogenation, either through a haloamine intermediate^24,37^ or ε-amine-HOX coordination.^38,39^ Given the structural similarities and our mutagenesis data, we hypothesize that K329 in esVHPO plays a similar role in HOX stabilization and delivery. To distinguish between covalent and non-covalent mechanisms, we employed a dansyl sulfinic acid probe previously used to detect chloramines through proximity-induced sulfonamide formation and “turn-on” fluorescent visualization (**Fig. 5D**).^40^ Isolated BrOx-esVHPO yielded strong fluorescent labeling following SDS-PAGE; removal of vanadate, H_2_O_2_, or KBr resulted in no signal (**Fig. 5E**). No signals were detected when bromide was substituted with chloride or iodide consistent with a lack of oxidation or HOX release, respectively. To probe K329 further, labeling was repeated with K329 variants (**Fig. 5F**). Both K329A and K329M abolished fluorescence, while K329R partially restored signal. This reinforces the importance of the positive charge in this position and the precedent of *N*-halogenated guanidine as a synthetic halogenating reagent support the putative covalent adduct.^41^ Contrastingly, the E327A mutant retained labeling, indicating that the carboxylate does not contribute to the formation or direct stabilization of oxidized bromide in esVHPO. This data supports a mechanism in which K329 facilitates the formation of a long-lived ε-bromamine intermediate necessary for substrate halogenation. This implicates lysine-mediated halogenation as a central feature of esVHPO catalysis.

### Cryo-EM captures a reaction intermediate of esVHPO during bromide oxidation

To gain insight into the structural basis of halide oxidation and halogenation, we determined the cryo-EM structure of BrOx-esVHPO. Using the same templated 2D classification approach on tilted and untilted datasets as for holo-esVHPO, we obtained an overall 2.4 Å reconstruction (PDBID 9PTS; EMD-71845) (**Extended Data Figs. 2, 7)**. Comparison of BrOx-esVHPO with holo-esVHPO reveals minimal global changes (RMSD: 0.57 Å^2^ for matching Cα atoms of residues 70-526) (**Fig. 6A**), but distinct local changes consistent with catalytic activation and a larger fraction of dimers with a well-resolved ICR (**Extended Data Figs. 3, 7**). Overall, the dimer compacts slightly, including the active site (**Fig. 6B**). Strikingly, BrOx-esVHPO contains unique density adjacent to the apical oxygen of vanadate and the ε-amine of K329 (**Fig. 6C**). We conservatively modeled this density as a water molecule due to the difficulty of assigning positively charged atoms in cryo-EM. However, the bond distance to the ε-amine is ∼2.0 Å, which aligns with a covalent N-Br bond^42^ while clashing with typical hydrogen bond distances. While little changed directly in the substrate site, CAVER analysis reveals the presence of a 12-Å long hydrophobic tunnel connecting the vanadate and substrate sites in BrOx-esVHPO that does not form in holo-esVHPO (**Extended Data Fig. 8**). This tunnel is composed of conserved residues, including K329, and mirrors analogous features in the FDHs where dynamic tunnel geometries gate HOX flow between co-factor and substrate sites.^25,43^ Combined with kinetic data, this indicates that subtle conformational changes regulate substrate bromination in esVHPO.

**Figure 6.**
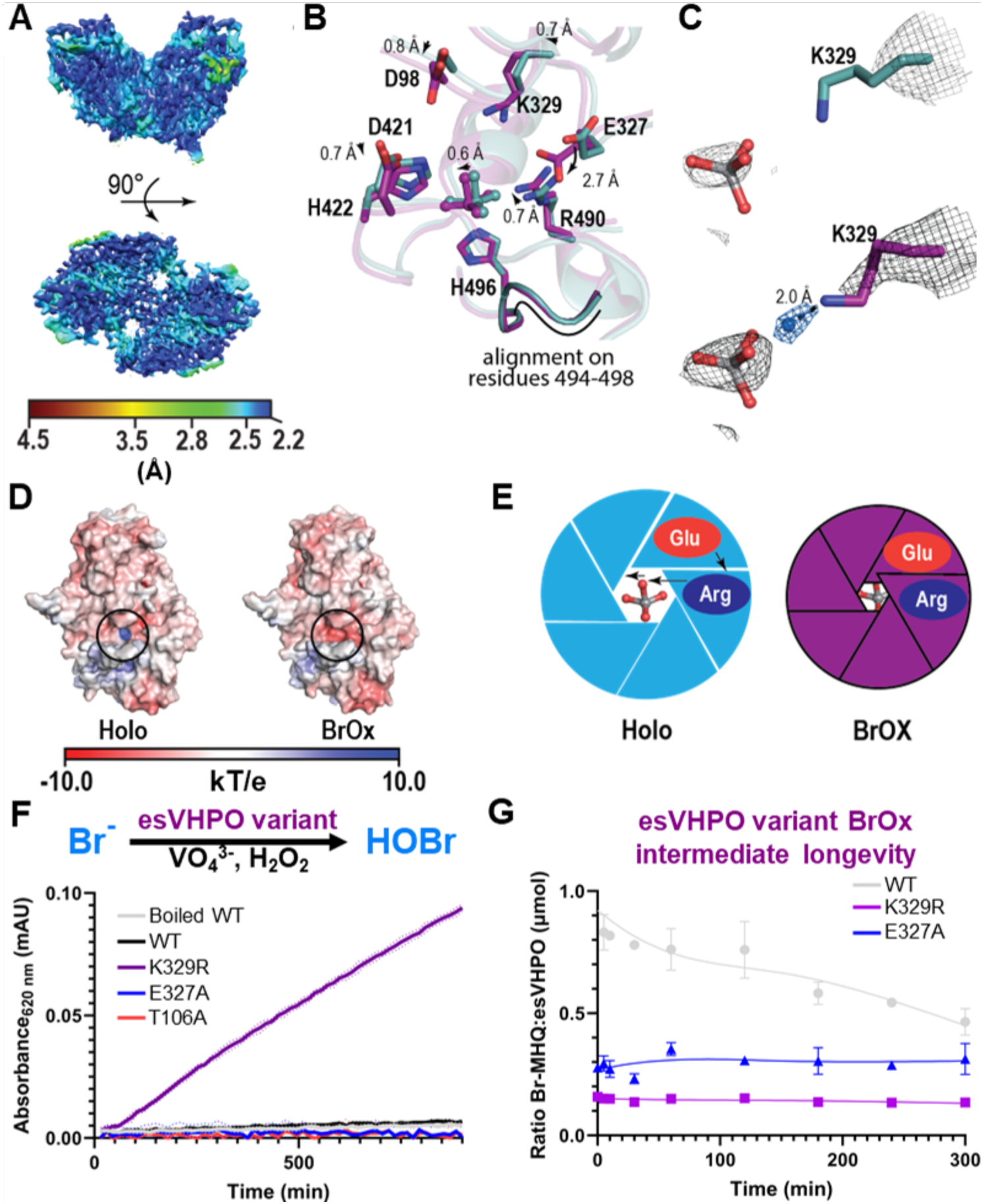
Cryo-EM structure of BrOx-esVHPO and the basis of HOBr retention. A) Local resolution map of the BrOx-esVHPO dimer (PDBID 9PTS). B) Overlay of the holo and BrOx-esVHPO active sites. The structures were aligned on residues 494-498, which contain the vanadate-coordinating histidine. Relative movements of atoms and groups are indicated by arrows and distances are indicated accordingly. C) Vanadate and K329 with respective potential maps shown at equivalent standard deviations. Both maps were normalized and scaled in Chimera.^53^ Holo-esVHPO is shown in teal, illustrating the disordered K329 sidechain. BrOx-esVHPO is shown in magenta. The K329 sidechain shows clear density and, at higher standard deviations, a covalentely bound bromide or coordinated water shown in blue appears 2.0 Å from the ε-amino group. D) Electrostatic potential surface of holo-esVHPO and BrOx-esVHPO. The black circle indicates the bromide entry tunnel towards the vanadate site. E) Schematic of the “open” shutter of holo-esVHPO and the “closed” shutter of BrOx-esVHPO. F) Thymol blue assay in the presence of bromide with selected esVHPO variants, with increasing absorbance representing the production of HOBr. G) Longevity of BrOx-esVHPO intermediates in selected mutant variants. Conditions are as in Fig. 5C with both variants 4 °C and previously shown WT longevity shown in gray for reference.

### An anionic gate prevents hypobromous acid dissociation

The most striking local change occurs in the vanadate site. The sidechains and loops surrounding vanadate-binding H496 show a concerted movement around the solvent tunnel to the vanadate site, which collectively acts as a “camera shutter” that dynamically encloses the core catalytic machinery (**Fig. 6B, D, E)**. This includes a 0.6 Å vanadate shift, general active site compaction, and a concerted 0.6-0.8 Å counterclockwise rotation of conserved residues D98, E327, K329, D421, and R490. Strikingly, BrOx E327 adopts an alternate rotamer, displacing its carboxylate by 2.7 Å, and forming a salt bridge with R490 that occludes solvent access to the vanadate site and changes the surface electrostatics of the putative halide-entry pore (**Fig. 6B, D, E)**. To test the functional implications of this gate on HOBr retention, we evaluated the iodide oxidation-capable E327A and K329R mutants. Performing the thymol blue assay in the presence of bromide with each esVHPO variant, we found esVHPO K329R was the only variant to freely diffuse HOBr, despite its impaired AQ bromination activity (**Fig. 6F**). Unlike wildtype, K329R and E327A were unable to form long-lived BrOx intermediates (**Fig. 6G**). Notably, K329R and E327A produced fluorescent signals when immediately derivatized by dansyl sulfinic acid (**Fig. 5F**). Together these data suggest that while a positively charged nitrogen is sufficient to form the bromamine intermediate, the primary amine of K329 and carboxylate of E327 are required for its longevity. The specific mechanism through which E327 stabilizes the haloamine is unclear. Most likely, the negative charge repels the hypohalite anion from dissociating through the halide tunnel or the E327-R490 salt bridge prevents the entry of water molecules and premature hydrolysis of the bromamine.

## Discussion

Despite distinct evolutionary origins, VHPOs and FDHs have convergently evolved remarkably similar strategies to achieve selective electrophilic halogenation. Both families feature spatially distinct co-factor and substrate binding sites connected by a hydrophobic tunnel and facilitated by a catalytic lysine. In FDHs, this lysine is paired with a conserved glutamate and is proposed to form either a covalent haloamine intermediate or a non-covalent coordination of HOX to direct aromatic halogenation; the glutamate catalyzes deprotonation of the Wheland-intermediate.^44,45^ Selective VHPOs lack an equivalent glutamate or conserved general base near the substrate site. E327 and D355 are the only basic residues in esVHPO positioned to fill this role, however mutagenesis indicates they are not directly involved in catalysis. Instead, structural and mutational evidence from esVHPO and NapH1 support a lysine-serine dyad (K329 and S429) that stabilizes a long-lived haloamine intermediate. The critical role of K329 is emphasized by the loss of substrate halogenation but retention of halide oxidation upon mutation in esVHPO and NapH1, as well as inducing HOBr diffusion by esVHPO K329R.^26^ Non-selective VHPOs, like *Ci*VCPO, appear to rely on electrostatic gradients to channel HOX from the solvent-exposed vanadate site toward the protein surface.^14^ This strategy excludes haloamine formation and explains the broader, substrate-directed reactivity of non-selective VHPOs. In contrast, selective VHPOs display minimal HOX leakage and high substrate specificity, reinforcing the presence of a protected oxidized halide intermediate. These mechanistic distinctions offer new opportunities for comparative enzymology to refine our overall understanding of enzymatic electrophilic halogenation.

While significant progress has been made in characterizing vanadate-dependent halide oxidation, the precise molecular determinants of halide specificity and substrate binding in VHPOs remain unclear. Early studies established that residues in the vanadate primary coordination sphere are essential for activity, while mutations to second sphere residues could alter halide specificity or enhance activity.^46^ However, these effects are not broadly applicable across homologs, suggesting subtle conformational changes play a more decisive role. Recent structures of *Cp*VBPO reveal that shifts of a leucine occludes water and enables halide access to the reactive vanadium(V)-peroxo species.^30,47^ Upon this shift, an adjacent aspartate is proposed to polarize a histidine critical to vanadium(V)-peroxo formation. Specific aromatic (W379 in esVHPO) or cationic residues have also been implicated in halide specificity, but their variability across VHPOs complicates generalization. Substrate binding architecture similarly varies across homologs. In *Ci*VCPO, preferred substrates are suggested to bind in a solvent-exposed groove where it reacts with channeled HOX.^14^ In contrast, a defined substrate binding pocket directly connected to the vanadate site is observed in the crystal structure of *Am*VHPO R425S, located at the obligate dimer interface.^15^ Both VHPOs feature solvent-accessible vanadate sites that release HOX in the absence of substrate. Selective VHPOs differ strikingly, conserving buried vanadate sites, the K329-S429 dyad, and distal substrate pockets that are absent in non-selective VHPOs. The absence of this dyad in the non-halogenating NapH3 homolog further supports its functional role.^26^ Anionic gating residues like E327 are conserved among selective VHPOs but not in non-selective VHPOs, suggesting a general mechanism for HOX retention. Our biochemical and structural data supports a mechanism where K373 polarizes the vanadium(V)-peroxo species to promote formation of the vanadate(V)-oxybromide intermediate (**Extended Data Fig. 9A**). This triggers conformational shifts involving E327 to close the vanadate pocket from solvent. The oxybromide is polarized through hydrogen bonding with S429 enabling nucleophilic attack by deprotonated K329 to form the bromamine (**Extended Data Fig. 9B**).

These mechanistic insights establish selective VHPOs as a distinct and tunable class of electrophilic halogenases. Their natural preference for non-aromatic sp^2^ carbon centers, like those in linear (mero)terpenoids or conjugated keto-enols, complements the aromatic selectivity of native FDHs, thus expanding the chemical space accessible to biocatalytic halogenation. Selective VHPOs do not always co-localize in biosynthetic gene clusters,^22,23,48^ therefore identifying sequence-based determinants of selectivity will aid in the discovery of VHPOs with novel substrates. In the absence of definitive structural observations of substrate binding in selective VHPOs, our docking and biochemical approach localizes the site of substrate binding for the first time in these homologs. This highlights the tunability potential of VHPO substrate selectivity, exemplified by the altered selectivity of D355A, and paves the way for rational engineering of VHPOs with non-native substrate scopes. By integrating structural and functional analyses, we have defined the molecular logic underlying halide oxidation and substrate halogenation in site-selective VHPOs. These principles lay the groundwork for the development of new halogenase biocatalysts capable of late-stage functionalization and selective transformations of relevant molecular scaffolds.

## Online Methods

### General materials and methods

All chemicals, media, and solvents used were purchased from commercial suppliers (Acros, Sigma Aldrich, Difco, Fisher, Oakwood, or Alfa Aesar) and used as received. PCR was run using a Mastercycler nexus – PCR Thermal Cycler (Eppendorf). Routine protein purification was performed on an ÄKTA go (GE Healthcare) FPLC and data was collected with UNICORN v 7.5 (GE Healthcare). Centrifugations were performed with a Centrifuge 5424 R (Eppendorf) (FA-45-24-11 rotor), a Sorvall Lynx 6000 (Thermo) (Fiberlite F9-6×1000 Lex rotor), and/or a Centrifuge 5804 R (Eppendorf) (S-4-72 rotor or FA-45-6-30 rotor). A BioSpectrometer kinetic (Eppendorf) and a NanoDrop One^C^ (Thermo) were used to collect UV-vis and optical density (OD_600_) readings. An EnVision HTS Multilabel Reader (Perkin Elmer) was used to record UV-Vis data in 96-well plate formats. Solid media bacterial cultures were grown in a Heratherm Compact Microbiological Incubator (Thermo) and liquid bacterial cultures were grown in a New Brunswick Scientific Excella E24 Shaker (Eppendorf) or a New Brunswick Inova44 (Eppendorf).

NMR Spectra were collected using an Avance III HD 500 MHz Spectrometer (Bruker) equipped with a BBFO SmartProbe CD_3_OD or DMSO-*d_6_* as solvent. Chemical shifts are reported in ppm and referenced to the CD_3_OD or DMSO-*d_6_* solvent signal (δ = 3.31 ppm or δ = 2.50 ppm for ^1^H, respectively). NMR splitting data are reported as follows: singlet (s), doublet (d), triplet (t), quartet (q), pentet (p), multiplet (m), coupling constant (J, Hz). UPLC-MS data was collected on an Elute UPLC instrument (Bruker) coupled to an Amazon ESI-ion trap mass spectrometer (Bruker) in positive mode. Compounds were separated in reversed-phase chromatography using 0.1% formic acid (Solvent A) and acetonitrile + 0.1% formic acid (solvent B) with a Bruker Intensity Solo C18(2), 2 mm x 100 mm column. The following LC separation method was used: 10% to 20% solvent B over 3 minutes, 20% to 100% solvent B over 3 minutes, 100% solvent B over 2 minutes, 100% to 90% solvent B over 1 minute, 90% to 10% solvent B over 2 minutes. All MS data was collected in positive mode. UPLC components were controlled with Compass Hystar 5.1 (Bruker), MS was controlled with ESI Compass trapControl v 8.0 (Bruker), and UPLC-MS data was analyzed with Compass DataAnalysis v 5.2 (Bruker). Synthetically prepared Br-MHQ (as prepared previously^23^) was used as a standard. All chromatograms were analyzed in BPC mode to confirm that side reactions or additional products were not produced. A calibration curve was prepared for Br-MHQ with the following concentrations appearing in the linear range and normalized against 250 nM nalidixic acid: 10 nM, 50 nM, 100 nM, 250 nM, 500 nM, 1.0 µM, 2.5 µM, 5 µM, and 10 µM. All mass spectrum chromatograms were exported to InkScape for graphical workup. All protein quantification was performed with the Bradford method using Quick Start Bradford 1x Dye Reagent (Bio-Rad Laboratories) and a standard curve generated with Bovine Serum Albumin (Thermo Scientific). Gel Filtration Standards (MW 1350-670,000 Da) (BioRad) were used to calibrate all gel filtration columns for size exclusion chromatography.

### General biological methods

Plasmid propagation and heterologous expression were performed in *Escherichia coli* DH10β competent cells (Invitrogen) and in *E. coli* BL21 (DE3) competent cells (New England Biolabs), respectively. *E. coli* was cultured in either Lysogeny Broth (LB) or Modified Terrific Broth (TB) (Fisher). Plasmids were transformed into competent *E. coli* strains with the following protocol. 100 µL aliquots of chemically competent *E. coli* were removed from -70 °C and thawed on ice. Plasmids were directly pipetted into the aliquot (1 µL of pure 40-60 ng/µL plasmid) and then gently tapped to mix. The mixture was then incubated on ice for 30 minutes before being placed at 42 °C for 45 seconds. After heat shock, cells were kept at room temperature for 2 minutes before transferring to 700 µL LB media and shaking for 1 hour at 37 °C. 200 µL of the cells were then plated on LB Agar supplemented with 50 µg/mL kanamycin. Plates were incubated for 16-20 hours at 37 °C. Single colonies were selected for all future applications.

### Preparation of gene constructs and protein purification

The gene for esVHPO was previously cloned into pET28a(+) vector and was expressed and purified as previously described.^23^ Briefly, esVHPO WT was transformed into competent *E. coli* BL21(DE3), inoculated into an overnight starter culture, inoculated again into 2 L of media, grown to an OD_600_ of 0.5-0.7 before induction with 100 µM IPTG, and expressed for 18 hours at 18 °C. Cells were lysed by sonication, clarified supernatant was prepared by centrifugation, purified by Ni-NTA affinity chromatography, buffer exchanged into cold storage buffer (25 mM HEPES-KOH pH 8.0, 300 mM KCl, 10% glycerol), and frozen at -70 °C. The purity acquired by Ni-NTA purification was sufficient for most applications; however if higher purity esVHPO WT was needed it was further purified *via* size exclusion chromatography (SEC) using a HiLoad 16/600 Superdex 200 pg (GE Healthcare Life Sciences) equilibrated in SEC Buffer (20 mM HEPES pH 8.0 and 300 mM KCl).

Mutated variants of esVHPO were cloned using the primers in **Table S2** and an *E. coli* codon optimized esVHPO WT gene cloned into pET28a(+) as template.^23^ PCR reactions were prepared on 25 µL scale with the following components: 12.5 µL 2X Q5 Polymerase master mix (New England Biolabs Inc.), 500 nM forward and reverse primers, 1 ng/25 µL template, and 3% DMSO. PCRs were run with the following cycling: Initial denaturation (98 °C, 30 seconds); 30 rounds of denaturation (98 °C, 30 seconds), annealing (50-65 °C, 5 seconds), and elongation (72 °C, 150 seconds); final elongation (72 °C, 500 seconds). Annealing temperature was adjusted as necessary for each primer pair. Linear PCR product was then ligated with KLD (kinase, ligase, DpnI) reaction mix (New England Biolabs Inc.) and transformed into *E. coli* DH10β competent cells. Successful mutations were confirmed *via* Sanger sequencing with high GC modifications (Azenta Life Sciences). esVHPO mutant variants were purified in the same manner as esVHPO WT. The following Ni-NTA purification yields were acquired: V102F (12 mg/L), T106A (3 mg/L), S110A (14 mg/L), E327A (1 mg/L), K329A (14 mg/L), K329M (22 mg/L), K329R (8.5 mg/L), I330A (0.3 mg/L), L333A (5 mg/L), G334F (14 mg/L), D355A (28 mg/L), T358A (5 mg/L), N359A (8 mg/L), S429A (2 mg/L), H436A (4.5 mg/L), H496A (47 mg/L).

To confirm general retention of fold, the wild type and the mutants were prepared for analytical gel filtration on a HiLoad 16/60 Superdex 200 pg (GE Healthcare Life Sciences) equilibrated in SEC Buffer. 1 mL of 1 mg/mL protein was loaded for each construct and ran at a maximum flow rate of 1 mL/min. Chromatograms were directly compared with esVHPO WT to judge if similar levels of aggregation and dimerization were present in each construct.

### Single-particle cryo-EM grid preparation

For all grids, esVHPO WT was purified by Ni-NTA affinity chromatography and SEC. Prior to preparation for grid application, esVHPO was buffer exchanged to a non-glycerol containing buffer (GF Buffer: 50 mM HEPES-KOH, 300 mM KCl, pH 8.0) and passed through a 0.22 µm filter. Three grids were prepared for each of the untilted data sets. esVHPO-holo was prepared as a mixture of 0.6 mg/mL esVHPO WT, 100 µM Na_3_VO_4_, and 20 mM KBr in GF buffer. esVHPO-BrOx was prepared as esVHPO-holo, however 1 mM H_2_O_2_ was added and the mixture was incubated at room temperature for 2 minutes prior to freezing on the sample grid. Each esVHPO state was prepared in the same manner as the untilted datasets, however 0.06 mg/mL esVHPO and 0.1 mM H_2_O_2_ were used instead. All samples were vitrified using a Vitrobot Mark IV (ThermoFisher Scientific) on UltrAufoill (Quantifoil) 300 mesh 1.2/1.3R grids at 4 °C and 1.5 s blotting time.

### Cryo-EM data collection

Initial screening sessions were performed at the UCSC Biomolecular cryo-EM Facility using a ThermoFisher Scientific Glacios cryo-Twin operating at 200 kV and coupled to a Gatan K2 Summit direct electron detector. The top-selected grids of holo-esVHPO sample were collected at 0° (untilted) and 30° tilted on a ThermoFisher Scientific Krios G2 operated at 300 kV at the UCSF Cryo-EM Facility. The microscope was equipped with a Gatan K3 summit detector and a Gatan Bioquantum energy filter operated at 20 eV. 8,832 and 4,819 micrographs were collected at a nominal magnification of 105kx and a physical pixel size of 0.845 A for the untilted and tilted datasets, respectively. The total electron exposure was 45.8 e-/ Å^2^ over 79 frames.

All BrOx-esVHPO datasets were collected at the Pacific Northwest Cryo-EM Center – PNCC (#160337) on a ThermoFisher Scientific Krios G3i equipped with a Gatan K3 summit detector. A Gatan Biocontinuum energy filter was operated with a slit width of 10 eV. The nominal magnification was 130kx and physical pixel size of 0.648 Å. Both datasets were collected in super-resolution mode with a total electron exposure of 50 e^−^/Å^2^ over 60 frames and 2,488 and 2,849 micrographs were collected for each 30°-tilted and untilted dataset. Data collection parameters are summarized in **Table S1**.

### Cryo-EM data processing

We performed all data processing using cisTEM.^49^ We initially picked particles using a maximum radius of 22 Å, a characteristic radius of 20 Å, a threshold of 0.1 and the highest resolution used for picking of 5 Å. Particles of the holo esVHPO dataset were extracted in 324 pix^2^ boxes, and particles of activated and charged datasets were extracted in 384 pix^2^ boxes. Tilted and untilted datasets were initially processed separately and merged after 2D classification. Data-processing parameters are summarized in **Table S1**.

We initially ran 2D classification using standard parameters without automasking over 20 rounds including progressively higher resolution data from 40 Å to 8 Å. We picked the best views for *ab initio* reconstruction for each dataset. We observed similar views in the tilted and untilted datasets, which led us to hypothesize that we either only pick top views or that alignments of rather small side-views fail. We optimized picking parameters and were able to slightly improve the angular distribution of particles. However, the improvement was not sufficient to obtain an isotropic reconstruction. To improve the alignment, we filtered the refined BrOX-esVHPO *ab initio* model to ∼10 Å and back-projected it in 10° steps. Accounting for the C2 symmetry, we obtained 210 views. We then used these views as templates for 2D classification for 20 rounds starting at 20 Å using 1° angular search steps. We initially ran 2D classification on each dataset individually and then merged particles from good classes from the respective tilted and untilted datasets. This significantly improved the angular distribution of the particles indicating that alignment of views on the narrow side of the particle remained challenging. Subsequently we ran 3D autorefinement using the *ab initio* model as the initial model for 3D alignment without automasking. Resolution-dependent filtering was enabled.

After obtaining a consensus reconstruction for both holo- and BrOx-esVHPO, we classified each dataset into 3 classes, repeating 10 rounds of classification to 5 Å without alignment followed by 3 rounds of alignment without classification, three times. For holo-esVHPO, only one class displayed a well-resolved ICR, whereas for BrOx-exVHPO, two classes were well resolved. The two classes were merged and subjected to 3 rounds of refinement with alignment to 5 Å. Final maps were sharpened in phenix.auto_sharpen^50^and DeepEMhancer^51^ to aide manual model building.

### Model building and validation

As a starting point for modeling, we generated a predicted model using ColabFold.^52^ Initially, we used rigid body fitting in Chimera^53^ and then manually adjusted the model in Coot.^54^ Each structure was iteratively refined using Phenix dev-5246,^55^ followed by further adjustments in Coot. Reconstructions were evaluated for anisotropy using the 3DFSC server (by calculating the 3D Fourier Shell correlations and sphericity of the reconstruction.^56^ Final models were evaluated using MolProbity,^57^ and model-to-map FSCs were obtained from Phenix.real_space_refine.^58^

### Structure and sequence visualization and analysis

Figures were prepared using PyMOL (The PyMOL Molecular Graphics System, Version 3.0 Schrödinger, LLC) and Chimera.^53^ Electrostatic potential molecular surfaces were calculated using the Adaptive Poisson-Boltzmann Solver plugin in PyMOL.^59^ Surfaces represent the Conolly surface (solvent excluded surface). Autodock Tools 4.2.6 were used to dock HHQ into holo-esVHPO.^60^ A model of HHQ was prepared in Avogadro and energy minimized with the MMFF94 force field.^61^ A search space for binding was specified in the putative substrate binding pocket and vanadate site. Twenty-five Genetic Algorithm runs were performed and data was output in Lamarckian 4.2; all other default settings were used. The lowest energy pose of HHQ was selected for further analysis. DynamicBind was used to dock the twenty lowest energy poses in dimeric holo-esVHPO, using the entire coordinate file as the search space and default settings.^35^ All sequences alignments for phylogenetic trees and phylogeny calculations were performed in Mega11.^62^ Sequences were aligned with the built in ClustalW algorithm. Phylogenetic analysis was then performed using Maximum Likelihood as the statistical method, 50 bootstraps, the Jones-Taylor-Thornton (JTT) model, using all sites for Gap/Missing data treatment, and the Nearest-Neighbor-Interchange heuristic method. Tree was exported and enhanced in InkScape to improve image quality and visual comprehension. The sequences from characterized non-selective and selective VHPOs, as well as uncharacterized VHPOs from unrepresented genera are listed in **Table S3**. Multiple sequence alignments were performed with ClustalW^63^ and visually prepared in ESPript.^64^ Conservation maps were generated on the ConSurf server using the ClustalW sequence alignments and holo-esVHPO coordinate file.^34^ Internal tunnels were calculated using CAVER 3.0.3 PyMol Plugin with default settings.^65^ Tunnel calculations were started from Glycine 334. The atoms from vanadate were included in all CAVER calculations.

### Steady state kinetics

All steady-state assays were performed in triplicate and analyzed *via* UPLC-MS. All reactions were performed in a 96-well clear flat-bottom polystyrene plate in a reaction volume of 100 µL. Each reaction consisted of 20 mM HEPES-KOH, 250 nM esVHPO WT, 100 µM Na_3_VO_4_, and 10% glycerol. When unvaried, the following substrate concentrations were used: 1 mM H_2_O_2_, 20 mM KBr, and 1 mM MHQ. For the K_0.5_ of MHQ, concentrations were varied from 5 µM to 250 µM. For the K_0.5_ of H_2_O_2_, concentrations were varied from 5 µM to 250 µM. For the K_0.5_ of KBr, concentrations were varied from 0.1 mM to 15 mM. Reactions were initiated simultaneously by the addition of 1 mM H_2_O_2_ and incubated for 5 minutes at 28 °C. For H_2_O_2_ variance, 20 mM KBr was used instead to initiate the reaction. Reactions were simultaneously quenched by the addition of 100 µL methanol. Reactions were transferred to microcentrifuge tubes and 250 nM nalidixic acid was added. If necessary, reactions were further diluted in methanol such that the final concentration of Br-MHQ product was expected to be 10 µM or less. Reactions were centrifuged for 3 minutes at 11000 x g and then transferred to UPLC-MS sample vials. Extracted ion peak areas of Br-MHQ product were normalized to the nalidixic acid internal standard peak area and converted to concentration with a calibration curve. Data was fit to the Hill equation in Prism (GraphPad):

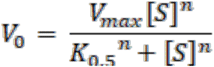

where V_0_ is the initial rate, V_max_ is the maximum rate of the reaction, [S] is the concentration of varied substrate, K_0.5_ is the concentration of substrate at one-half of V_max_, and n is the hill coefficient. The steady state kinetics for C3-Deuterated-MHQ (D-MHQ) was performed precisely as for undeuterated MHQ, but MHQ was replaced with equal concentrations of D-MHQ. Kinetic isotope effect (KIE) was calculated by dividing the catalytic efficiency vs MHQ by the catalytic efficiency vs D-MHQ.

### Iodide oxidation assay

Thymol blue assay was prepared according to previous work.^66^ Briefly, a 250 µL reaction was prepared for each esVHPO variant, in triplicate, with the following components: 10 µM thymol blue, 40 µg/mL esVHPO construct, 100 µM Na_3_VO_4_, 2 mM KI, and 20 mM HEPES-KOH (pH 8.0). Reactions were initiated by the addition of 40 µM H_2_O_2_ and monitored for five hours at 15 minute intervals at 635 nm on an Envision plate reader. For each reaction, a blank solution was prepared by abstaining H_2_O_2_ addition. For each time point, the blank solution was subtracted from the reaction solution to account for spontaneous iodide oxidation. Control reactions without the presence of enzyme or in the presence of boiled esVHPO WT were run to account for spontaneous oxidation of iodide by hydrogen peroxide. The same protocol was used to monitor HOBr production by esVHPO variants but used 10 mM KBr in place of KI. esVHPO K329R bromide concentration gradients were prepared as above but with 5 mM, 50 mM, or 500 mM KBr.

### Alkyl quinolone bromination assay

The bromination of AQ substrates (HHQ, MHQ, NHQ, HQNO, and DHQ) by esVHPO WT and the mutant constructs were assayed as previously performed.^23^ Briefly, 100 µL reactions of 100 µM AQ substrate, 2.5 µM esVHPO construct, 100 µM Na_3_VO_4_, 20 mM KBr, 20 mM HEPES-KOH, and 10% glycerol were prepared in triplicate. Reactions were initiated with addition of 1 mM H_2_O_2_ and reacted for 30 minutes at 28 °C and 200 RPM. Each reaction was quenched with an equal volume of methanol, then further diluted 5x in methanol (final dilution of 1:10) and spiked with 250 nM nalidixic acid. The extent of bromination for each construct and substrate was analyzed by UPLC-MS.

### esVHPO-BrOX isolation and single-turnover

A 2.5 mL reaction consisting of 1.4 mg/mL esVHPO WT, 20 mM KBr, 100 µM Na_3_VO_4_, 1 mM H_2_O_2_, and 20 mM HEPES-KOH, pH 8.0 was prepared and incubated for 15 minutes at 28 °C. Reaction mixture was then buffer exchanged into SEC buffer using a pre-equilibrated PD-10 column. For calculation of the concentration of esVHPO-BrOx, it was assumed that 100% of the population was in the state. Reactions were immediately prepared after buffer exchange. Five 100 µL reaction conditions with varied amounts of esVHPO-BrOx (0 µM, 1µM, 2.5 µM, 5 µM, and 10 µM esVHPO-BrOx), 100 µM MHQ, and 20 mM HEPES-KOH, pH 8.0 were then prepared in triplicate. Reactions were initiated by the addition of enzyme and incubated for 30 minutes at 28 °C. Reactions were then quenched with an equal volume of methanol and analyzed *via* UPLC-MS.

To determine the time-based stability of esVHPO-BrOx, the intermediate was prepared and isolated as above, however esVHPO-BrOx was stored on ice or at 28 °C for the duration of the time course. Reactions were prepared after esVHPO-BrOx storage on ice or at 28 °C for the following lengths of time: 0 minutes, 5 minutes, 15 minutes, 30 minutes, 60 minutes, 2 hours, 3 hours, 4 hours, and 5 hours. The reactions conditions for each time point were prepared as above. The same reaction was performed with esVHPO E327A and esVHPO K329R.

### Dansyl sulfinic acid derivatization

Each esVHPO WT reaction condition was prepared as 100 µL reactions with 2 mg/mL enzyme. Each reaction condition was ran in the presence of 300 µM Na_3_VO_4_, 20 mM KX (where X is the appropriate halide), and 1 mM H_2_O_2_, as appropriate. Each reaction condition was incubated at 30 °C for 15 minutes. 5 mM L-methionine was added to each reaction and allowed to react for 5 minutes at room temperature to quench excess unreacted H_2_O_2_. 7.5 mM dansyl sulfinic acid (50-fold molar excess esVHPO) was added to each reaction and incubated at 37 °C and 200 RPM for 1 hour. In the case of the positive control, a 10-fold molar excess of dansyl chloride was added instead. Reactions were shielded from light during these incubation steps. 20 µg of protein was then prepared for SDS-PAGE in Laemmli buffer, boiled for 10 minutes and loaded on a 12% SDS-PAGE gel. Electrophoresis was performed at 200 V until the dye front was run off the gel. Gels were fluorescently imaged with GelDoc Imager (BioRad) using the Alexa 488 blotting program. After fluorescent imaging, gels were stained with Coomassie blue and then imaged photographically. The same reaction conditions were used for esVHPO K329A, K329M, K329R, and E327A.

## Supporting information

esVHPO-cryoEM-SI-PDF

## Acknowledgements

We gratefully acknowledge financial support from the National Institutes of Health (R35-GM147235 to S.M.K.M), and the University of California—Santa Cruz for startup funding (S.M.K.M and S.L.). We appreciate the contributions of H.-W. Lee, L. Sanchez, and B. Rabbits (University of California Santa Cruz) for assistance and maintenance of nuclear magnetic resonance, high resolution mass spectrometry, and plate-reader instrumentation, respectively. We thank Dr. Marzia Miletto and the Pacific Northwest Center for Cryo-EM (PNCC) at Oregon Health & Science University for data collection and support on data processing. A portion of this research was supported by NIH grants U24GM129547 and R24GM154185 performed at the PNCC at OHSU and accessed through EMSL (grid.436923.9), a DOE Office of Science User Facility sponsored by the Office of Biological and Environmental Research. We thank Dr. David Bulkley and the University of California, San Francisco Cryo-EM Facility for data collection. We also acknowledge the Biomolecular cryo-Electron Microscopy Facility at the Department of Chemistry and Biochemistry of the University of California—Santa Cruz (RRID: SCR_021755) for scientific and technical assistance (NIH High-End Instrumentation program, S10-OD02509). Molecular graphics and analyses performed with UCSF ChimeraX, developed by the Resource for Biocomputing, Visualization, and Informatics at the University of California, San Francisco, with support from NIH R01-GM129325 and the Office of Cyber Infrastructure and Computational Biology, National Institute of Allergy and Infectious Diseases.

## Author contributions

J.T.B., S.L., and S.M.K.M. designed research; J.T.B., L.A.V., J.T.C., and V.H.B.S. performed research; J.T.B., S.L., and S.M.K.M. analyzed data; J.T.B., S.L., and S.M.K.M. prepared the manuscript with contributions from all authors.

## Data Availability Statement

The structural models and cryo-EM maps have been deposited in the PDB and EMDB under accession codes 9PTR (holo-esVHPO) and 9PTS (BrOx-esVHPO), and EMD-71844 (holo-esVHPO) and EMD-71845 (BrOx-esVHPO), respectively. Uncropped SDS-PAGE gels, PDB validation reports, and NMR FIDs are provided with this paper. All other data is available upon request.

## Extended data Figures and legends

**Extended Data Figure 1.**
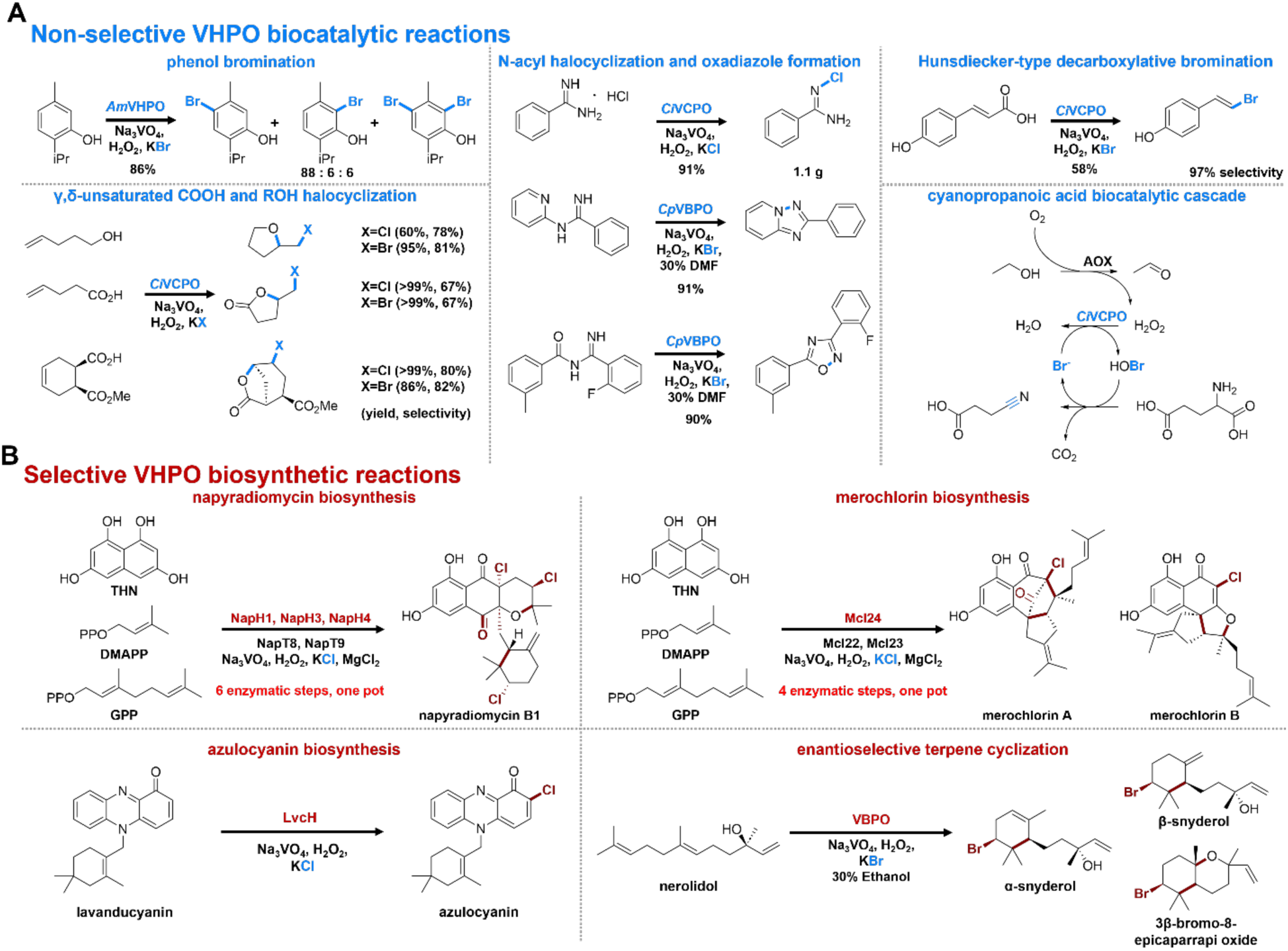
VHPO biocatalytic and biosynthetic reactions. A) Biocatalytic reactions developed using non-selective VHPOs. Installed halogens, bonds to halogens, or bonds formed through cryptic halogenation are highlighted in blue. Examples of phenol bromination,^17^ alcohol or carboxylic acid halocycilization,^68^ N-acyl halocyclization,^69^ Hunsdiecker decarboxylation,^70^ and a biocatalytic cascade are given.^71^ B) Enantioselective and/or regioselective biosynthetic reactions catalyzed by VHPOs. Bonds formed and halogens installed by VHPOs are highlighted in red. The reactions involving VHPOs in napyradiomycin B1 biosynthesis,^72^ the reactions involving Mcl24 to form merochlorins,^73^ the reaction catalyzed by LvcH to form azulocyanin,^22^ and the enantioselective terpene cyclization catalyzed by macroalgal VBPO are shown.^13^

**Extended Data Figure 2.**
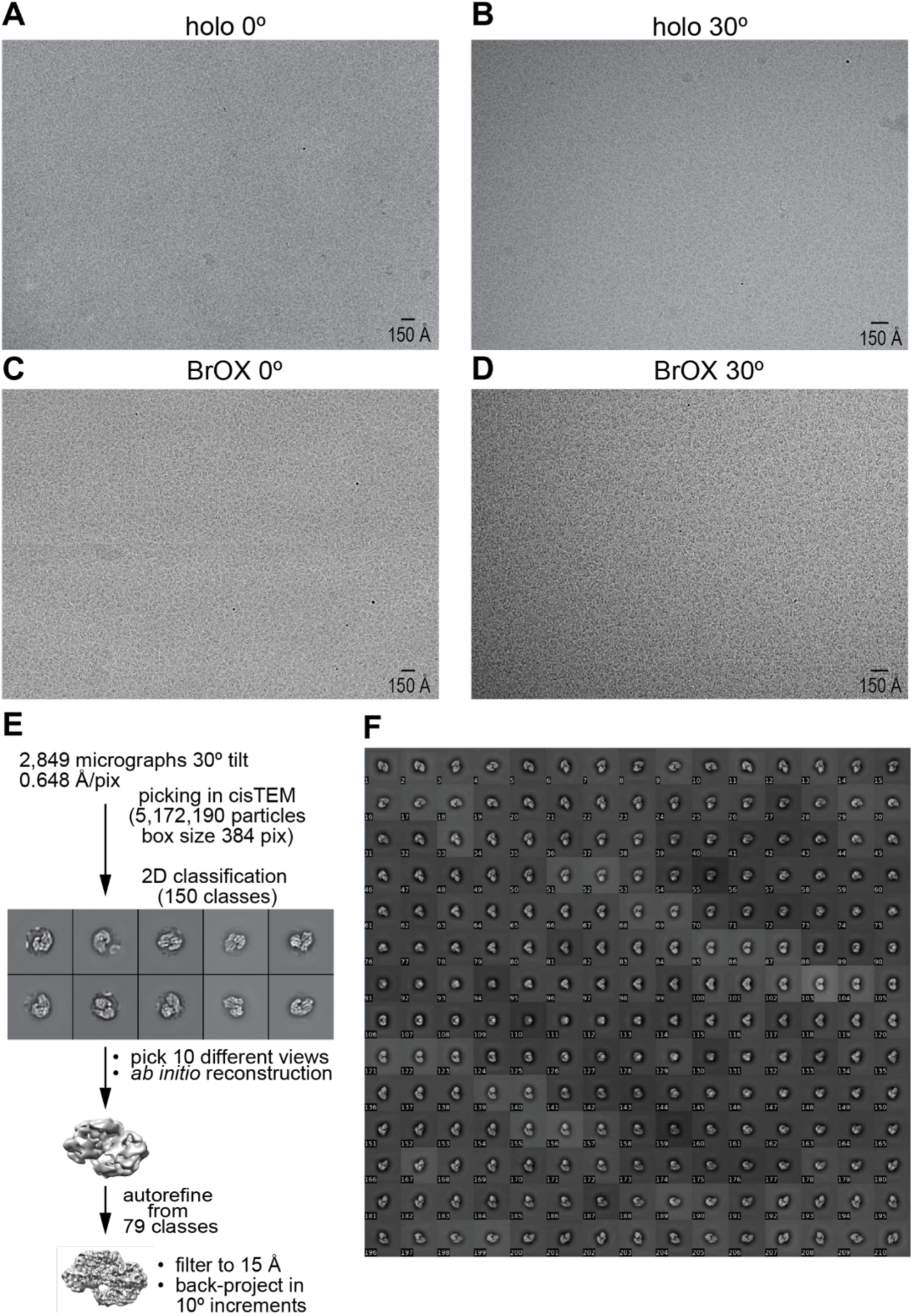
Representative micrographs and 2D template generation. Representative micrograph of A) untilted holo-esVHPO, B) tilted holo-esVHPO, C) untilted BrOx-esVHPO, and D) tilted BrOx-esVHPO. E) To generate 2D templates for templated 2D classification, particles were initially picked from the tilted BrOX-esVHPO dataset and subjected to 2D classification into 150 classes. Ten classes representing different orientations were picked to generate an ab initio reconstruction in cisTEM. Particles extracted from all good classes were then used in autorefinement. To generate the 2D templates, the reconstruction was filtered to 15 Å and back-projected in 10° increments, yielding 210 classes when accounting for C2 symmetry. F) 2D templates used for templated 2D classification.

**Extended Data Figure 3.**
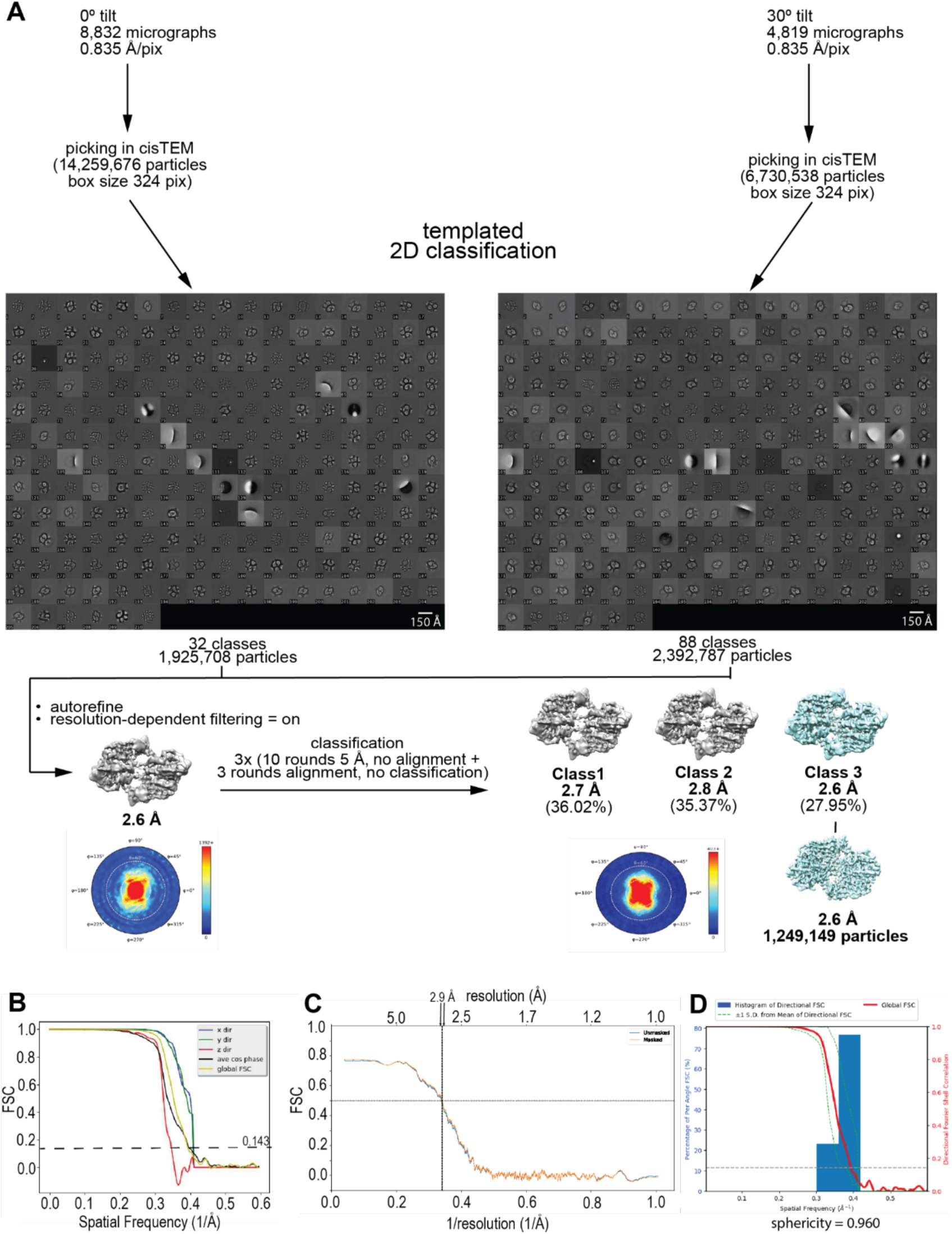
Data processing workflow and validation of holo-esVHPO dataset. **A)** Tilted and untilted datasets were initially processed separately using templated 2D classification. After 2D classification, the best classes were extracted and particles were subjected to autorefinement with resolution-dependent filtering enabled. Subsequently, particles were classified into three classes by repeating 10 rounds of classification without alignment followed by three rounds of alignment without classification three times. The best class showing a resolved ICR was extracted for postprocessing. **B)** 3D FSC curves and global FSC calculated using the 3DFSC server. **C)** Model-to-map FSC calculated in phenix.real_space_refine. **D)** Histogram of directional FSC and calculated sphericity.

**Extended Data Figure 4.**
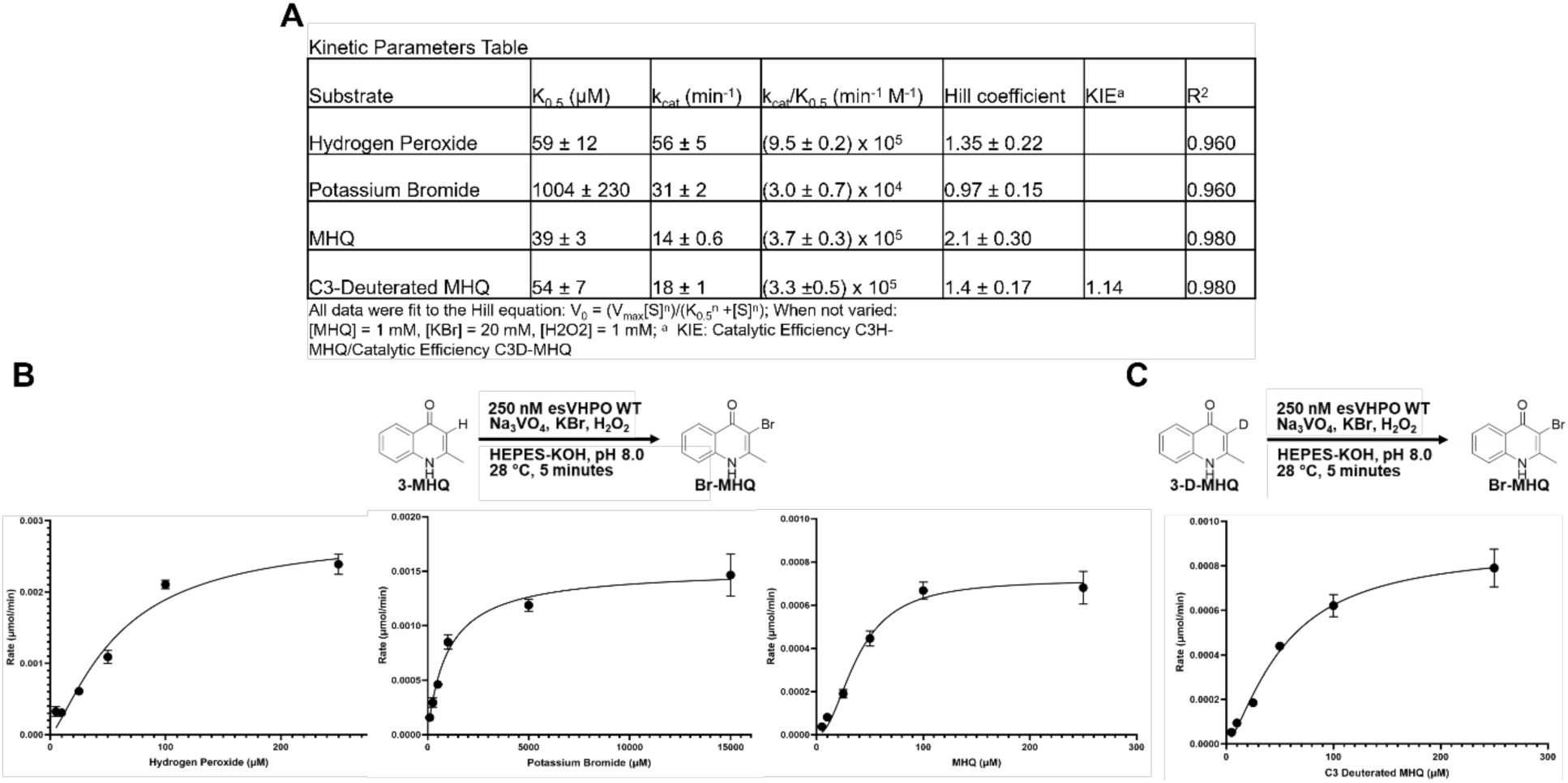
esVHPO wild-type steady state kinetic parameters. A) Steady-state parameters for Hill equation fit to each dataset. Fitting was performed in GraphPad Prism. B) General reaction scheme for each kinetic dataset involving MHQ. Plots of each triplicate dataset are shown with error bars to represent each point and the line of best fit. Error bars represent the mean ± s.d. (n = 3). C) As in B, however varied MHQ was replaced with a C-3 deuterated equivalent (3-D-MHQ). Error bars represent the mean ± s.d. (n = 3).

**Extended Data Figure 5.**
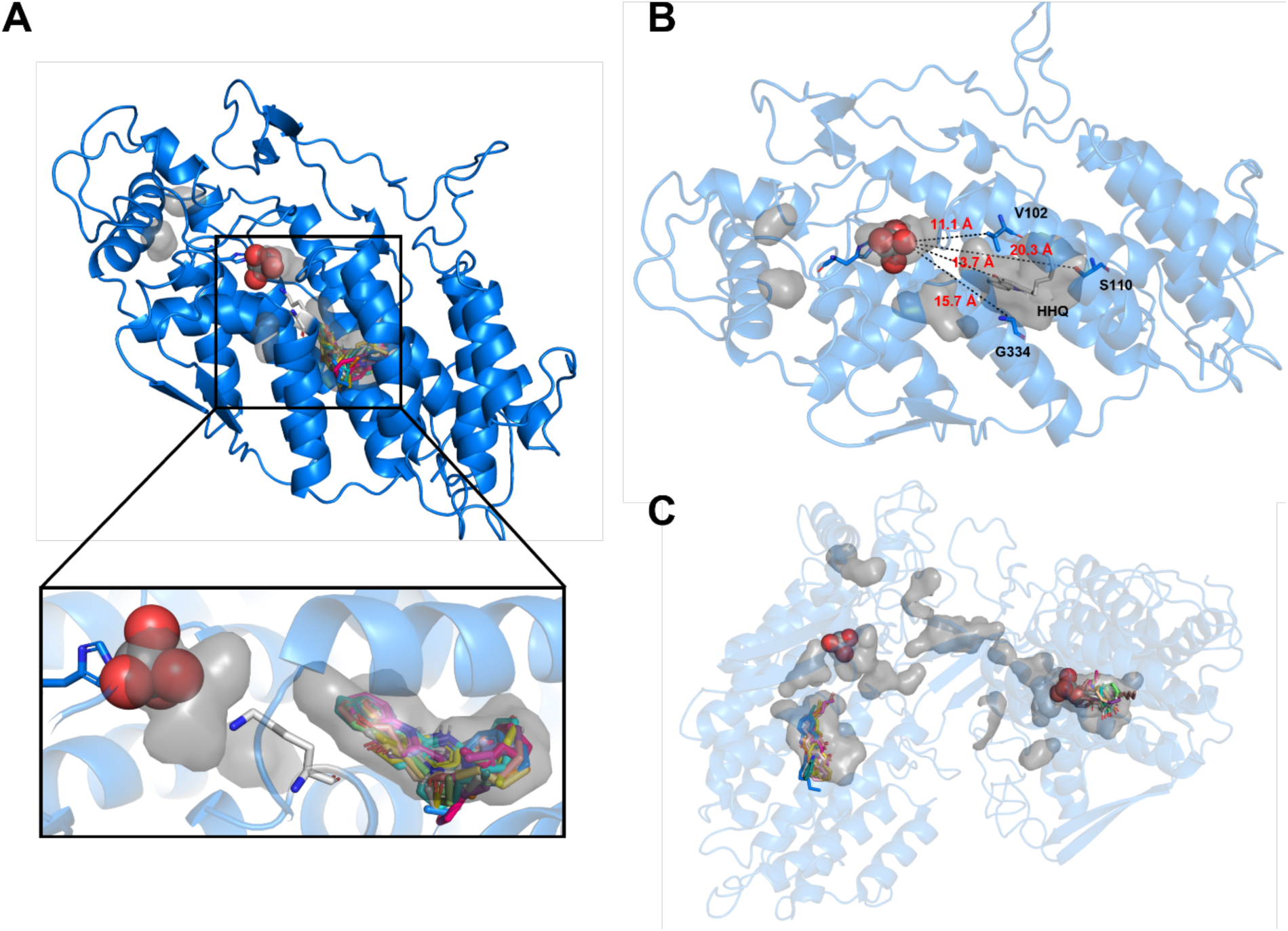
Holo-esVHPO substrate docking. Holo-esVHPO was prepared and docked with HHQ using AutoDock Tools.^60^ Vanadate was removed prior to running docking simulations and returned to the coordinate file after docking poses were simulated for visual clarity. Holo-esVHPO is represented in blue, vanadate as spheres, and internal cavities in grey. A) The twenty lowest energy poses of HHQ docked in monomeric holo-esVHPO. Docking search space was restricted to encompass the vanadate and substrate sites. B) Distance of HHQ and selected residues in the substrate site from vanadate. C) Holo-esVHPO dimer docking with HHQ was performed using NeuroSnap DynamicBind.^35^

**Extended Data Figure 6.**
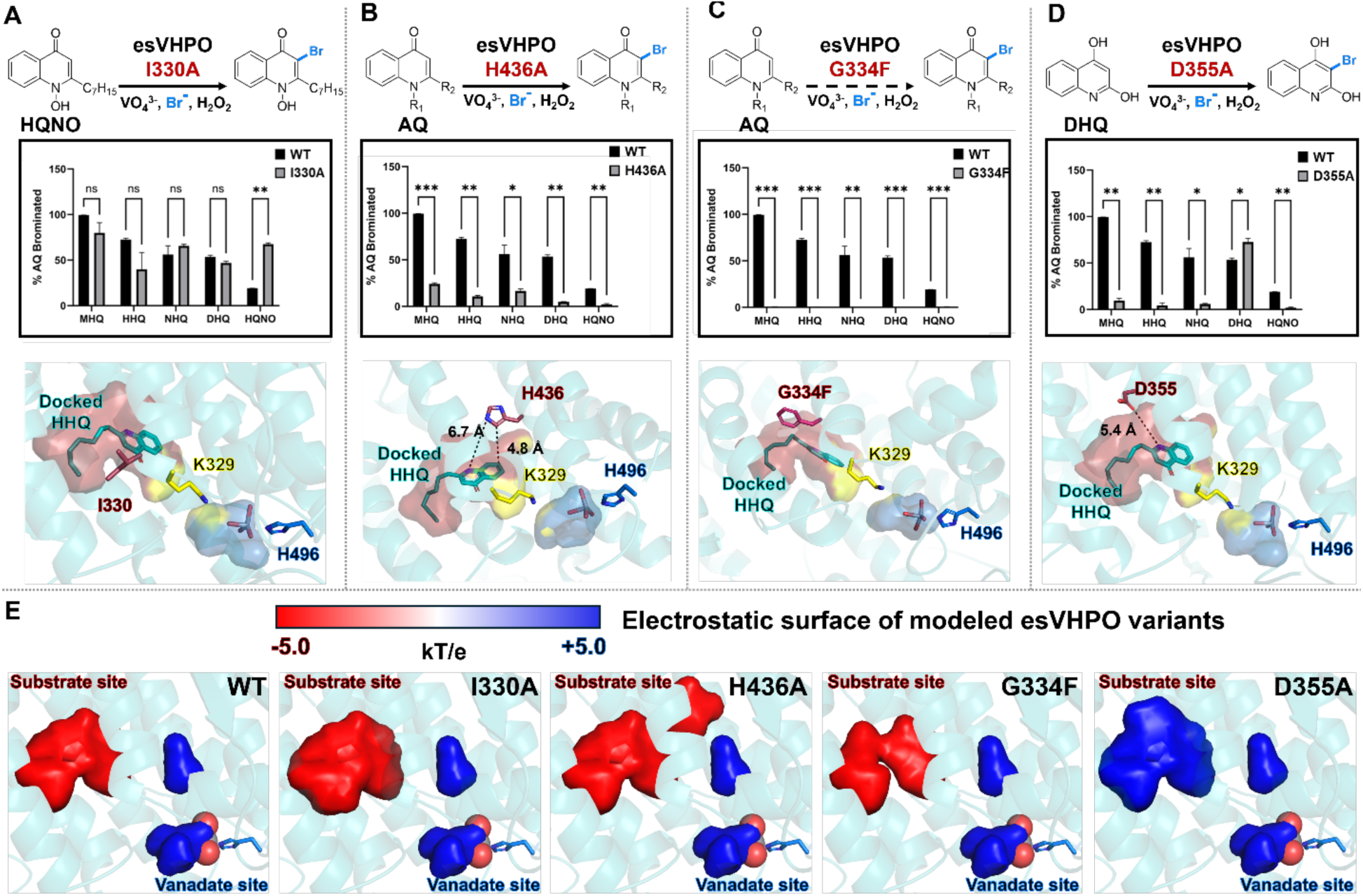
Representative esVHPO variant substrate profiles and modeled effects of mutation on structural features. Reactions catalyzed by selected esVHPO variants. The fold change in bromination compared to wild type is shown. Error bars represent the mean ± s.d. (n = 3). Significance was calculated with the multiple paired t-test. The following p-values were used: >0.033 (ns), <0.033 (*), <0.003 (**), <0.001 (***). HHQ-docked holo-esVHPO variants were modeled in PyMol. Colors are as **Figure 4A**. **E)** Wild-type and the modeled mutated variants were analyzed with APBS electrostatics. Internal cavities are shown as surface representations.

**Extended Data Figure 7.**
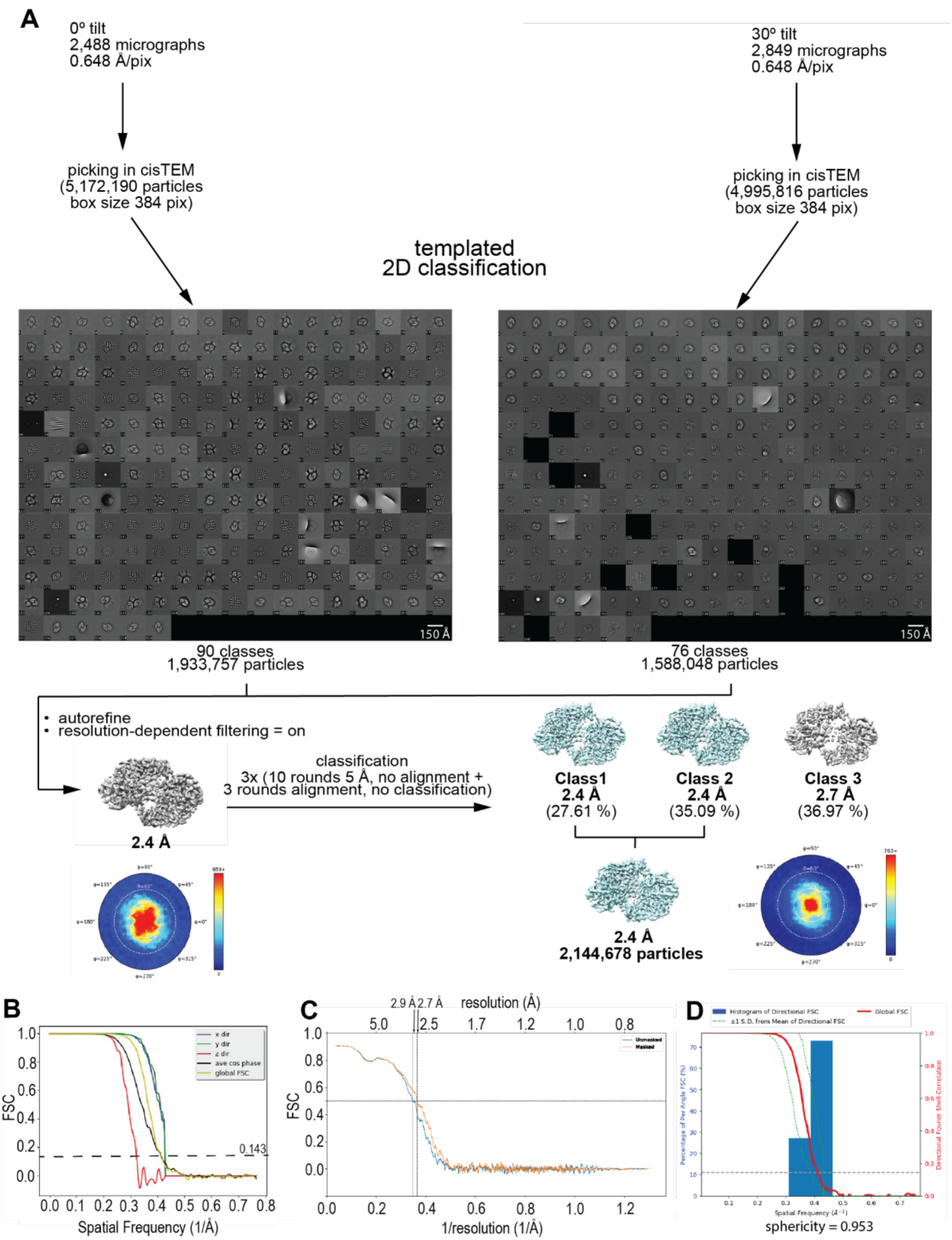
Data processing workflow and validation of BrOx-esVHPO dataset. **A)** Tilted and untilted datasets were initially processed separately using templated 2D classification. After 2D classification, the best classes were extracted and particles were subjected to autorefinement with resolution-dependent filtering enabled. Subsequently, particles were classified into three classes by repeating 10 rounds of classification without alignment followed by three rounds of alignment without classification three times. The best class showing a resolved ICR was extracted for postprocessing. **B)** 3D FSC curves and global FSC calculated using the 3DFSC server. **C)** Model-to-map FSC calculated in phenix.real_space_refine. **D)** Histogram of directional FSC and calculated sphericity.

**Extended Data Figure 8.**
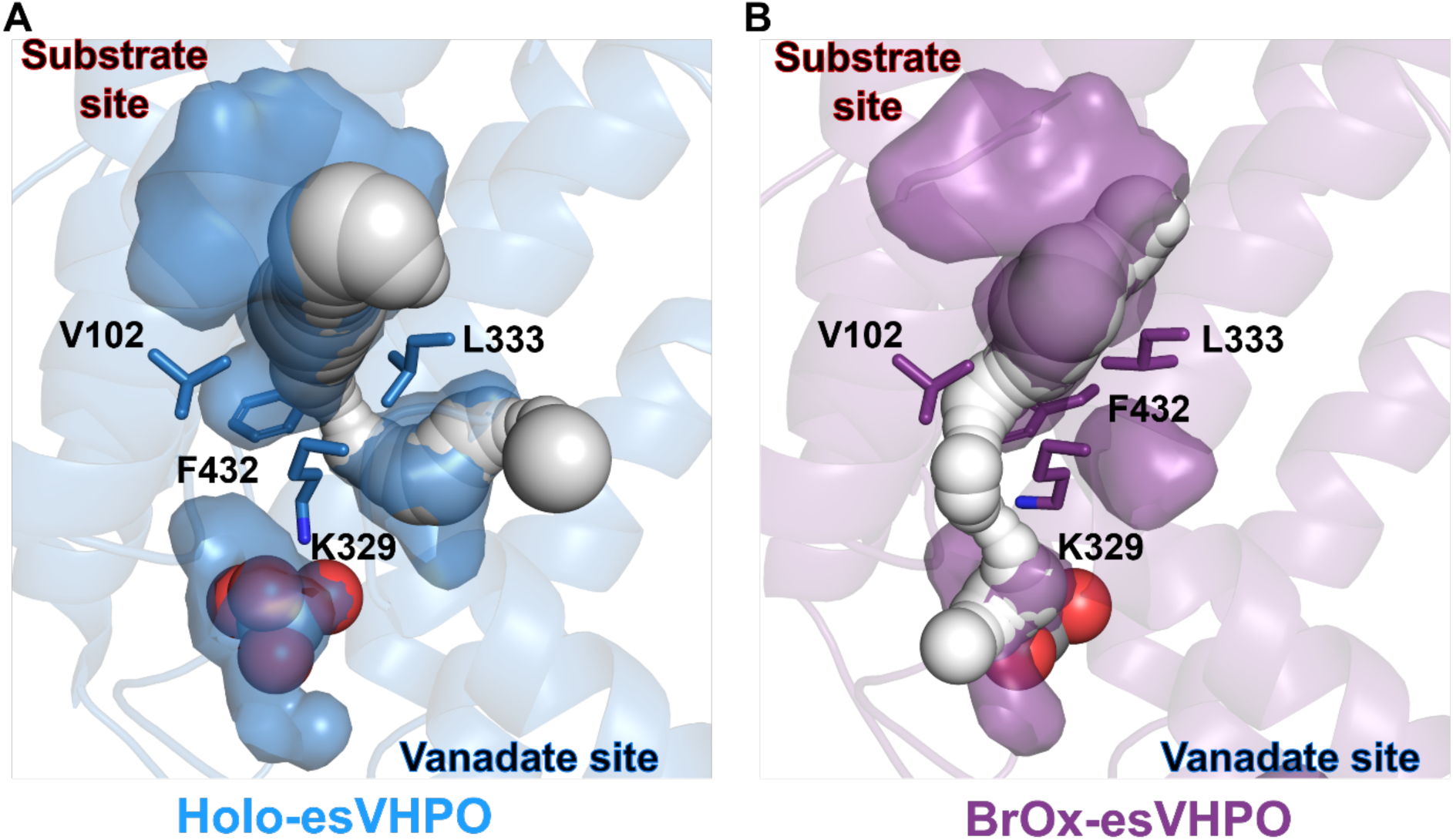
CAVER analysis of esVHPO hydrophobic HOX tunnel. **A)** Holo-esVHPO and **B)** BrOx-esVHPO cryo-EM models were analyzed with the CAVER 3.0 PyMol plugin. Default probe settings were used, vanadate was included in the calculation, and G334 was set as starting point. 3-dimensional structure is represented in ribbons, internal cavities as surface, vanadate as spheres, tunnel forming residues as sticks, and the tunnel as white spheres.

**Extended Data Figure 9.**
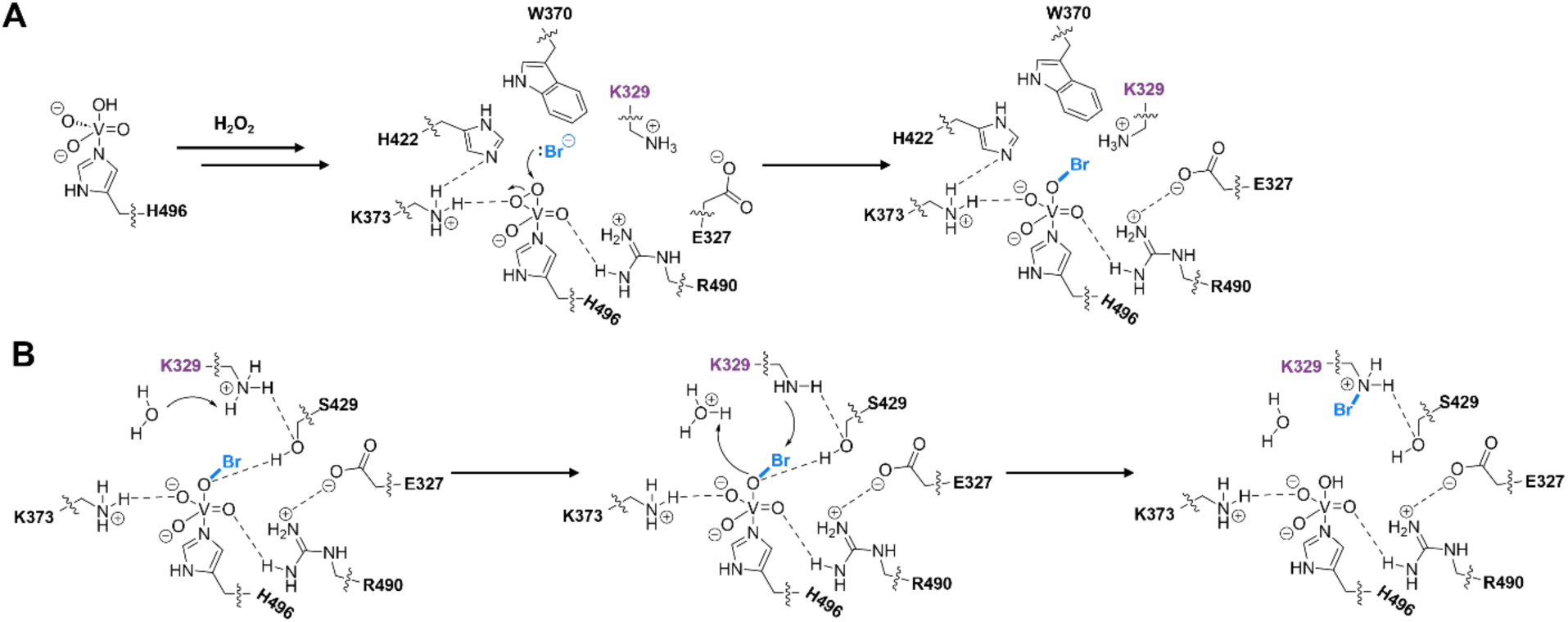
Mechanistic insights in esVHPO halide oxidation and haloamine formation. A) Formation of the vanadate(V)-oxybromide intermediate in esVHPO. Bond distances and electron density imply a strong interaction of K373 with the vanadate oxygen stabilizes the breaking of the peroxo bridge. The salt bridge between R490 and E327 is expected to form during this step. B) Formation of the bromamine intermediate in esVHPO. Bond distances and the requirement of S429 for bromination imply a role in vanadate(V)-oxybromide polarization. A conserved water residue is most likely involved in K329 deprotonation. The positively charged bromamine intermediate is likely stabilized through S429 interactions prior to binding organic substrate.

