## Supplementary material for "Separation of halide oxidation and substrate halogenation chemistries rationalizes site-selective vanadium dependent haloperoxidase catalysis": esVHPO-cryoEM-SI-PDF

**This PDF file includes:**

Figures S1 to S12
Tables S1 to S3
Synthetic Methods
NMR spectra
References

### Table of Contents

1. **Supplementary Figures** **S3**

   Figure S1. *Graphical representation of selective VHPO halide oxidation behavior* S3

   Figure S2. *Topology diagram of Holo-esVHPO* S4

   Figure S3. *Structural homology of esVHPO to highest similarity VHPO crystal structures in the PDB* S5

   Figure S4. *Comparison of FDH and VHPO tunnel residues* S7

   Figure S5. *Sequence alignment of AQ-VHPOs* S8

   Figure S6. *Sequence alignment of characterized selective VHPOs* S9

   Figure S7 *Substrate bromination activity of the soluble aggregate* S10

   Figure S8. *esVHPO variant oligomerization compared to Wild Type* S11

   Figure S9. *Time course of thymol blue assay esVHPO variants* S13

   Figure S10. *AQ bromination profile of esVHPO variants compared to wild type* S14

   Figure S11. *Electrostatic surfaces of structurally resolved selective and non-selective VHPOs* S16

   Figure S12. *Substrate dependence of K329R for HOBr production* S17

2. **Supplementary Tables** **S18**

   Table S1. *Cryo-EM Data collection, reconstruction, and structure refinement parameters* S18

   Table S2. *Primer table* S19

   Table S3. *VHPO sequences used for phylogenetic tree construction* S21

3. **Synthetic Methods** **S23**

   *2-methyl-3-iodo-quinolin-4(1H)-one* S23

   *2-methyl-3-deutero-quinolin-4(1H)-one* S23

   *N-N-dimethyl-1-amino-5-naphthalenesulfinic acid* S24

4. **NMR Spectra** **S25**

   *2-methyl-3-iodo-quinolin-4(1H)-one* S25

   *2-methyl-3-deutero-quinolin-4(1H)-one* S28

   *N-N-dimethyl-1-amino-5-naphthalenesulfinic acid* S31

5. **References** **S34**

### Supplementary Figures

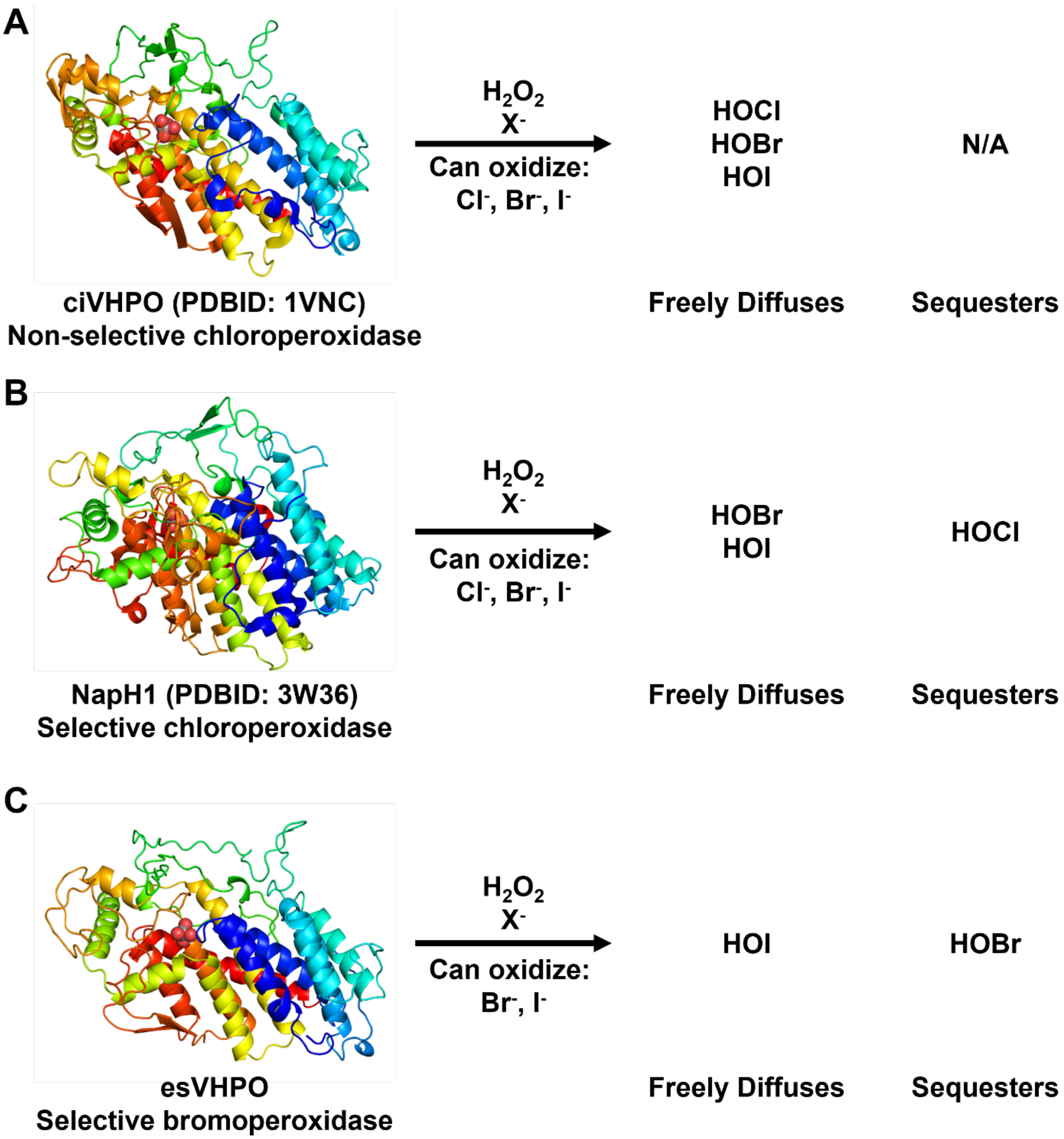

**Figure S1. Graphical representation of selective VHPO halide oxidation behavior.** Control over the oxidized halide species by different types of VHPOs. Freely diffused HOX is detectable by scavenging methods like MCD and TB assays, while retained HOX production can only be detected through reaction with native substrate or derivatization.

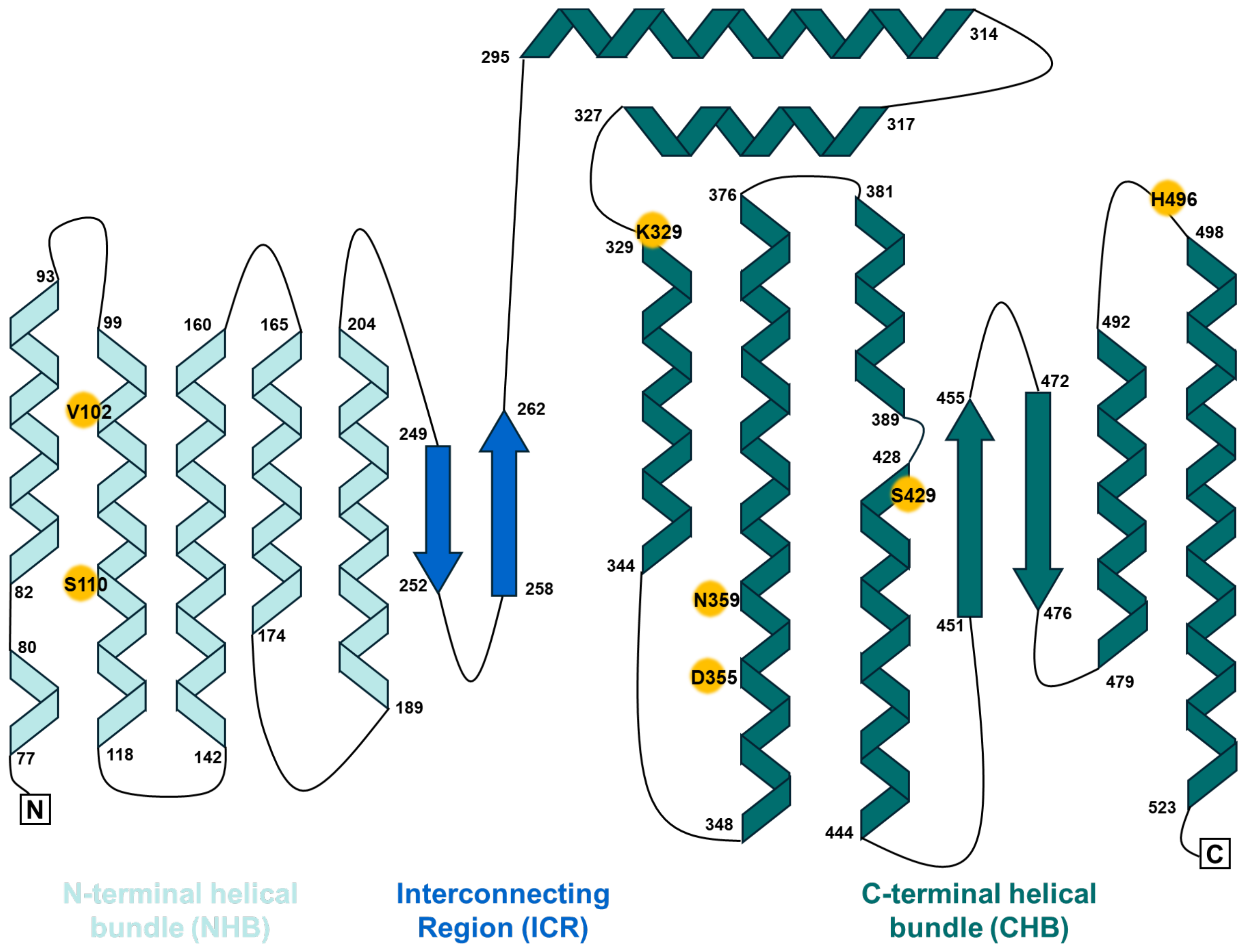

**Figure S2**. **Topology diagram of Holo-esVHPO**. esVHPO is composed of three regions, the N-terminal helical bundle (NHB, red), the Interconnecting Region (ICR, purple), and the C-terminal helical bundle (CHB, blue). The NHB is composed of helix 1 (residues 77-93), helix 2 (residues 99-118), helix 3 (residues 142-160), helix 4 (residues 174-165), and helix 5 (residues 189-204). The ICR is composed of strand 1 (residues 249-252) and strand 2 (residues 258-262). The CHB is composed of helix 6 (residues 295-314), helix 7 (residues 317-327), helix 8 (residues 329-344), helix 9 (residues 348-376), helix10 (residues 381-389), helix 11 (residues 428-444), strand 3 (residues451-455), strand 4 (residues 472-476), helix 12 (479-492), and helix 13 (residues 498-523). Selected residues from the substrate binding site (S110, N359, D355), the halide tunnel (V102, K329), and the vanadate binding site (S429, H496) have been highlighted to localize the pocket regions to the topology diagram. Topology diagram was drawn in Microsoft PowerPoint.

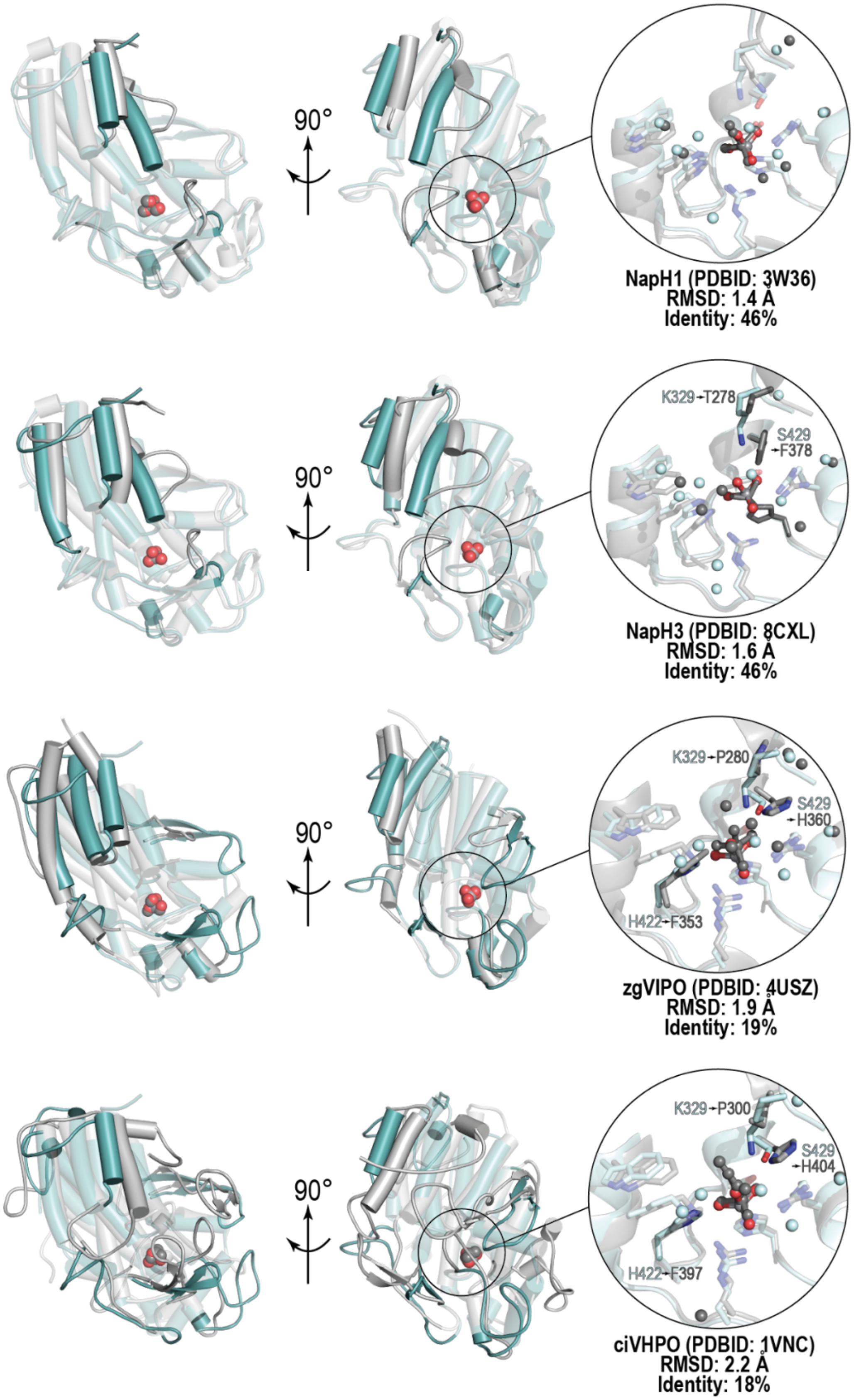
90°
NapH1 (PDBID: 3W36)
RMSD: 1.4 Å
Identity: 46%
90°
K329→T278
S429
→F378
NapH3 (PDBID: 8CXL)
RMSD: 1.6 Å
Identity: 46%
90°
K329→P280
S429
→H360
H422→F353
zgVIPO (PDBID: 4USZ)
RMSD: 1.9 Å
Identity: 19%
90°
K329→P300
S429
→H404
H422→F397
ciVHPO (PDBID: 1VNC)
RMSD: 2.2 Å
Identity: 18%

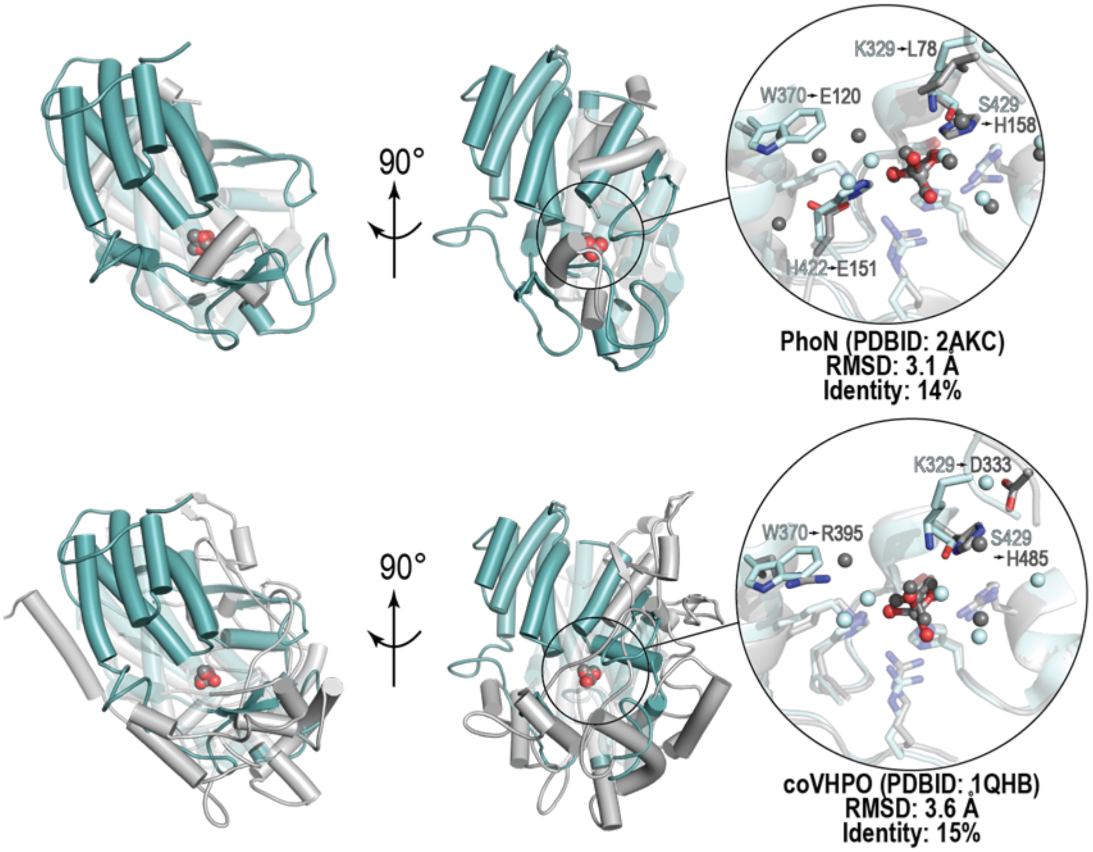

**Figure S3. Structural homology of esVHPO to highest similarity VHPO crystal structures in the PDB**. The Cryo-EM holo-esVHPO model was queried against the entire PDB through the DALI server.[1] The six most similar non-redundant sequences were VCPO NapH1 from *Streptomyces* sp. CNQ-525 (PDBID: 3W36), non-halogenating VHPO NapH3 from *Streptomyces* sp. CNQ-525 (PDBID: 8CXL), zgVIPO1 from *Zobellia galatanivorans* (PDBID: 4USZ), ciVHPO from *Curvularia inaequalis* (PDBID: 1VNC), PAPII-phosphatase, PhoN from *Salmonella enterica* subsp. *enterica serovar Typhimurium* (PDBID: 2AKC), and coVHPO from *Corallina officinalis* (PDBID: 1QHB). RMSD and identity values were calculated by the DALI server. Monomeric units of each protein were aligned with TopMatch.[2] In all cases, esVHPO was the query (teal) and the other VHPO was the target (grey).

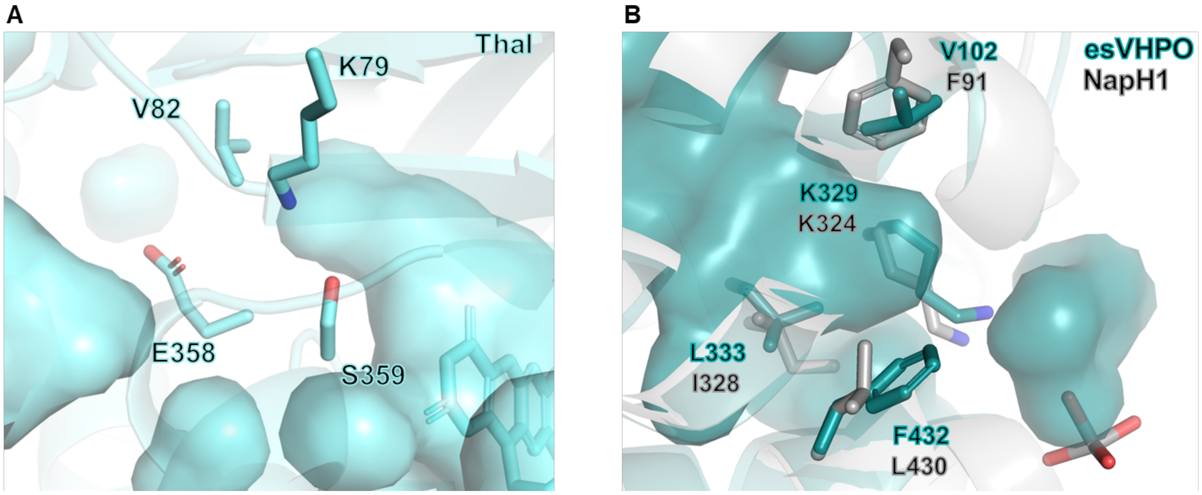

**Figure S4. Comparison of FDH and VHPO tunnel residues**. (A) Hydrophobic tunnel residues that gate the flavin co-factor site and tryptophan binding site in the flavin-dependent halogenase Thal (PDB ID: 7CU2).[3] These residues form the bottleneck that controls the access of HOX to the substrate site. (B) Overlay of esVHPO-holo (PDB ID: 9PTR) and NapH1 (PDB ID: 3W36). Residues that occupy comparable gating positions compared to Thal are highlighted.

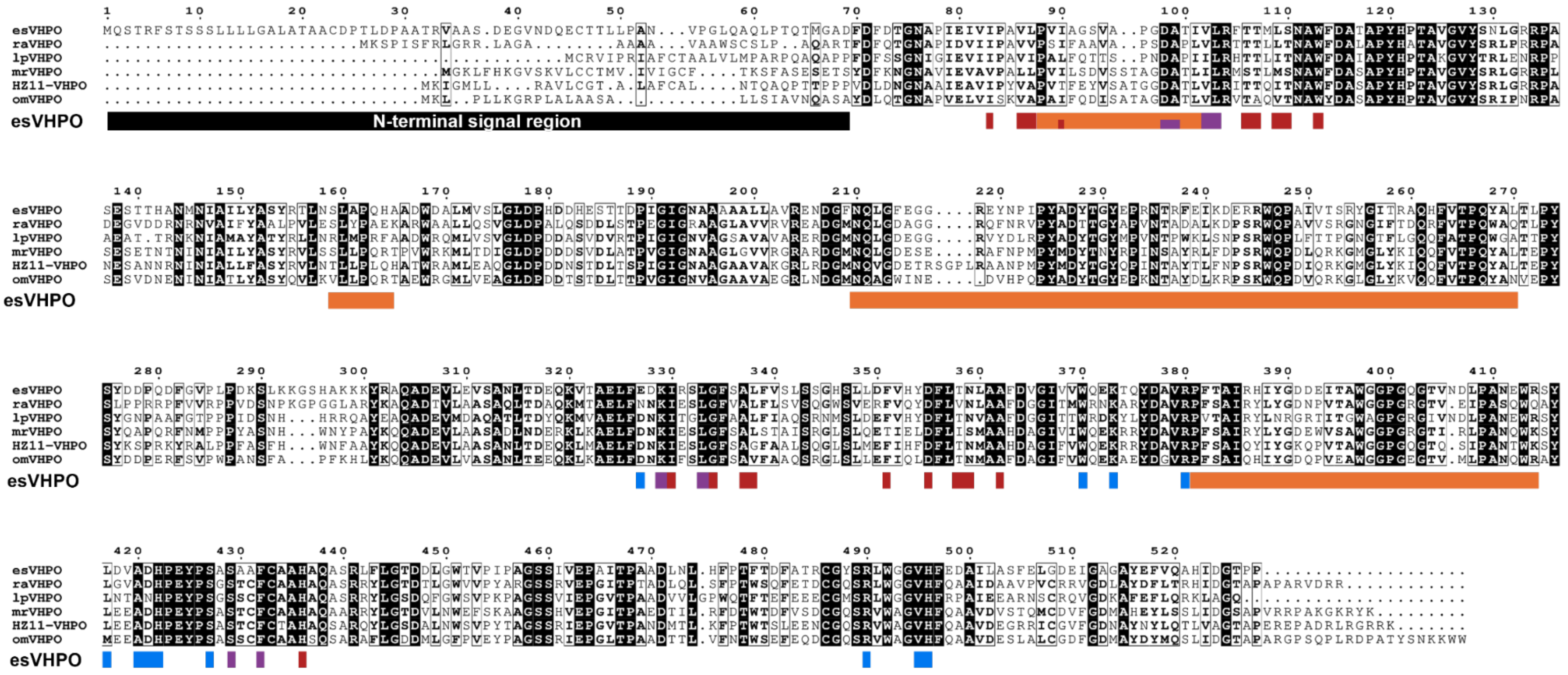

**Figure S5**. **Sequence alignment of AQ-VHPOs.** Sequence alignment was generated in ClustalW[4] and visualized in ESPript (https://endscript.ibcp.fr).[5] Only VHPOs with characterized alkyl quinolone (AQ) C3-bromination activity were selected for the sequence alignment. The sequence region corresponding to the conserved presence of a variable secretion signal peptide is designated with a black rectangle. Structural regions of esVHPO are shown below the sequence alignment as colored rectangles. Red rectangles represent residues that contribute to the substrate binding site, purple rectangles represent residues that contribute to the halide tunnel, blue rectangles represent residues that contribute to the vanadate site, and orange rectangles represent residues that contribute to the dimerization domain. Split rectangles represent residues that contribute to multiple regions.

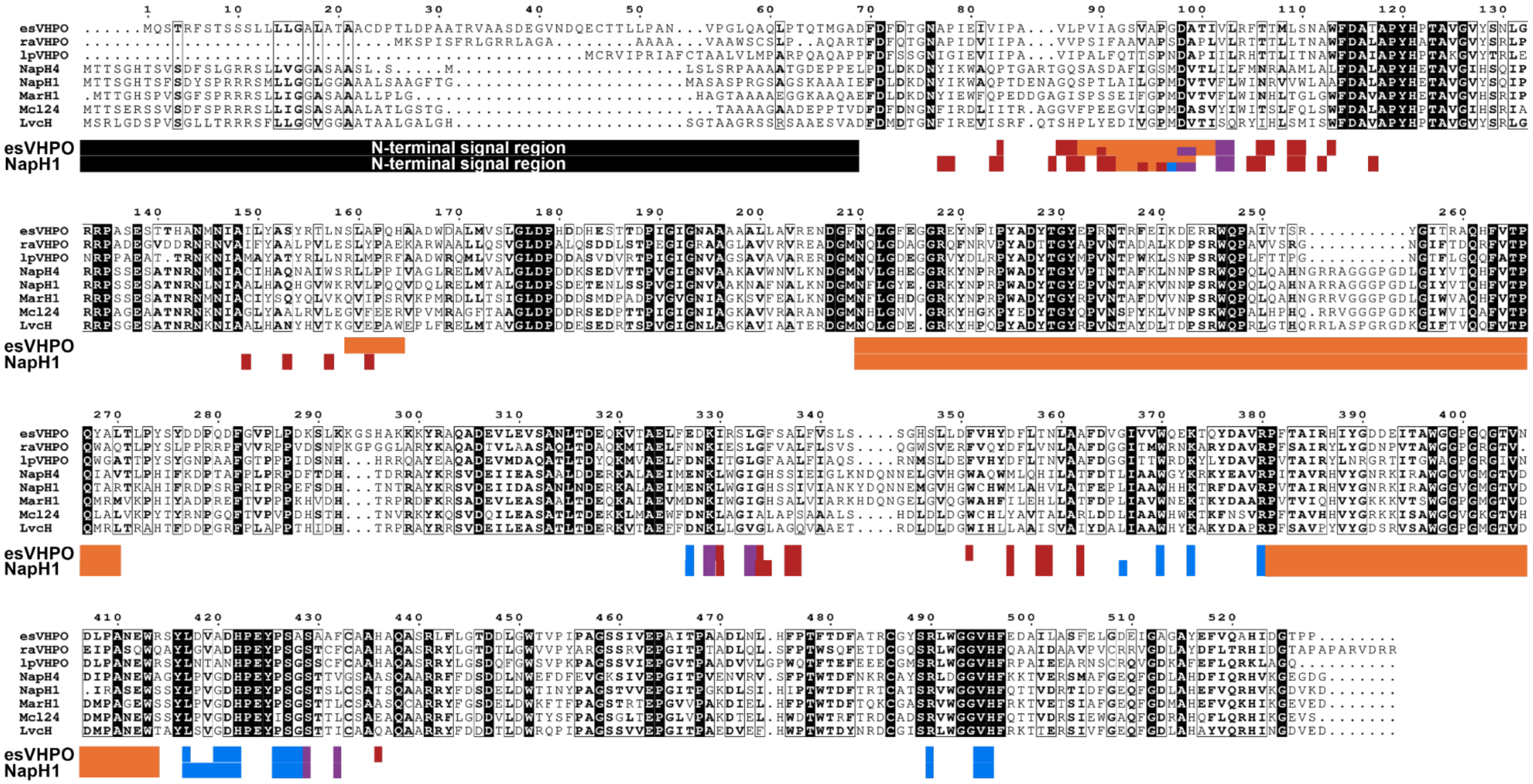

**Figure S6**. **Sequence alignment of characterized selective VHPOs.** Sequence alignment was generated in ClustalW[4] and visualized in ESPript (https://endscript.ibcp.fr).[5] Three representatives of characterized selective AQ bromination VHPOs (esVHPO, raVHPO, and lpVHPO) and five representative characterized selective meroterpenoid chlorination VHPOs (NapH4, NapH1, MarH1, Mcl24, and LvcH) were selected for this alignment. The sequence region corresponding to the conserved presence of a variable secretion signal peptide is designated with a black rectangle. Structural regions of esVHPO and NapH1 are shown below the sequence alignment as colored rectangles. Red rectangles represent residues that contribute to the substrate binding site, purple rectangles represent residues that contribute to the halide tunnel, blue rectangles represent residues that contribute to the vanadate site, and orange rectangles represent residues that contribute to the dimerization domain. Split rectangles represent residues that contribute to multiple regions.

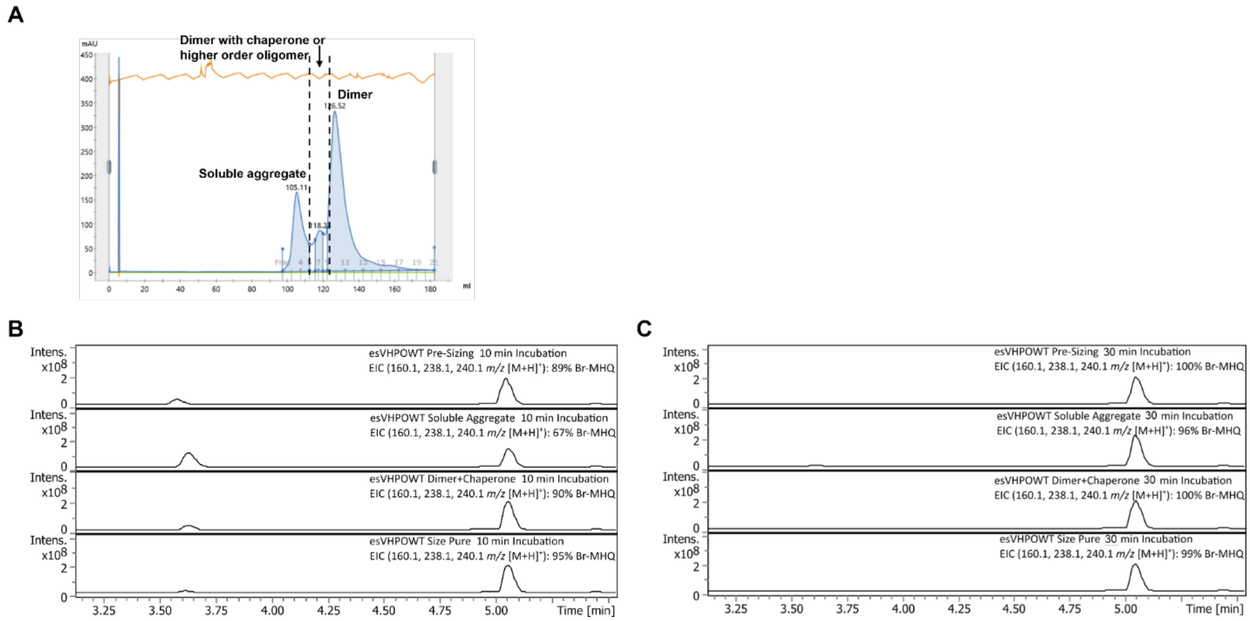

**Figure S7**. **Substrate bromination activity of the soluble aggregate.** (A) Size exclusion chromatogram of esVHPO WT. 10 mg of protein was loaded onto a 120 mL gel filtration column. Protein elution volumes were standardized to BioRad Gel Filtration Standards. Soluble aggregate and dimer were collected separately. The mixture of dimer with chaperone or higher order oligomer could not be distinguished fully and was also collected separately. (B) Bromination of MHQ by esVHPO WT under standard conditions. Reaction was checked after 10 minutes and was ran for non-size purified esVHPO WT and the collected SEC fractions from panel A. MHQ substrate elutes at 3.65 minutes and Br-MHQ product elutes at 5.05 minutes. (C) The same as panel B, however the reaction was run for 30 minutes.

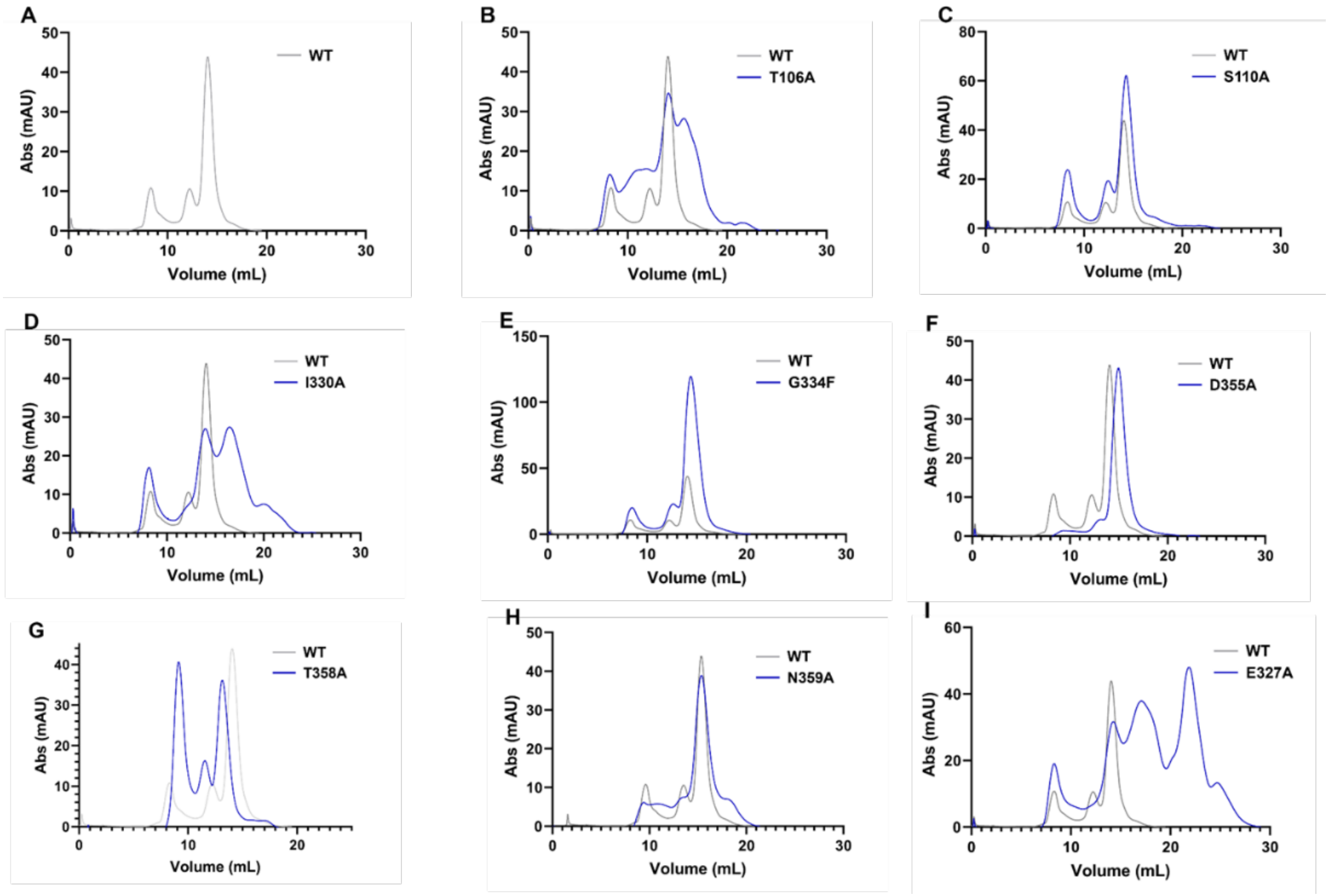
A
B
C
D
E
F
G
H
I
Abs (mAU)
Volume (mL)
WT
T106A
S110A
I330A
G334F
D355A
T358A
N359A
E327A

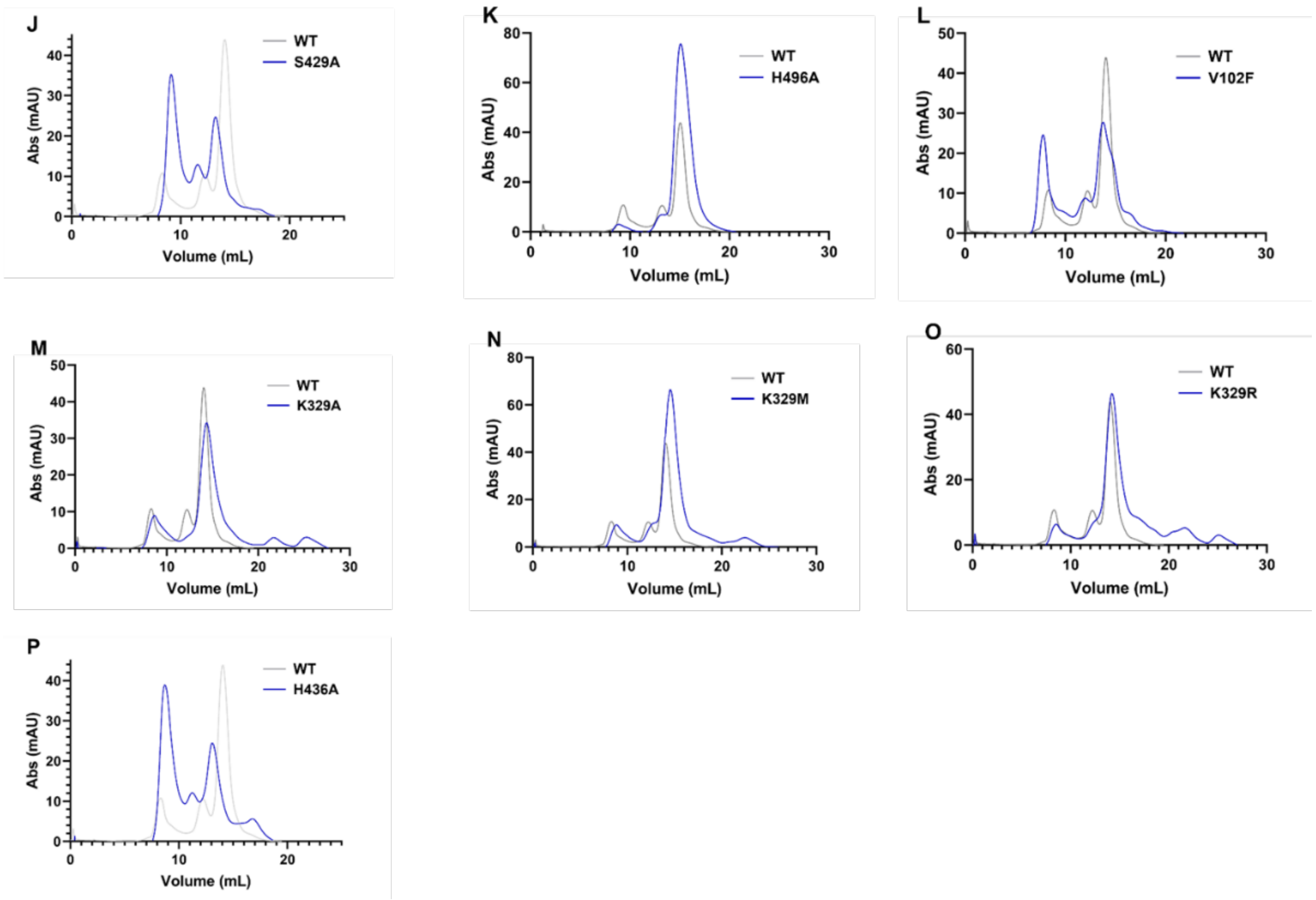

**Figure S8**. **esVHPO variant oligomerization compared to Wild Type.** Size exclusion chromatograms for esVHPO mutant constructs. For each panel, the chromatogram for the labeled mutant construct is shown in dark blue while the WT chromatogram is shown in grey. The same amount of protein was ran on the gel filtration column for all constructs

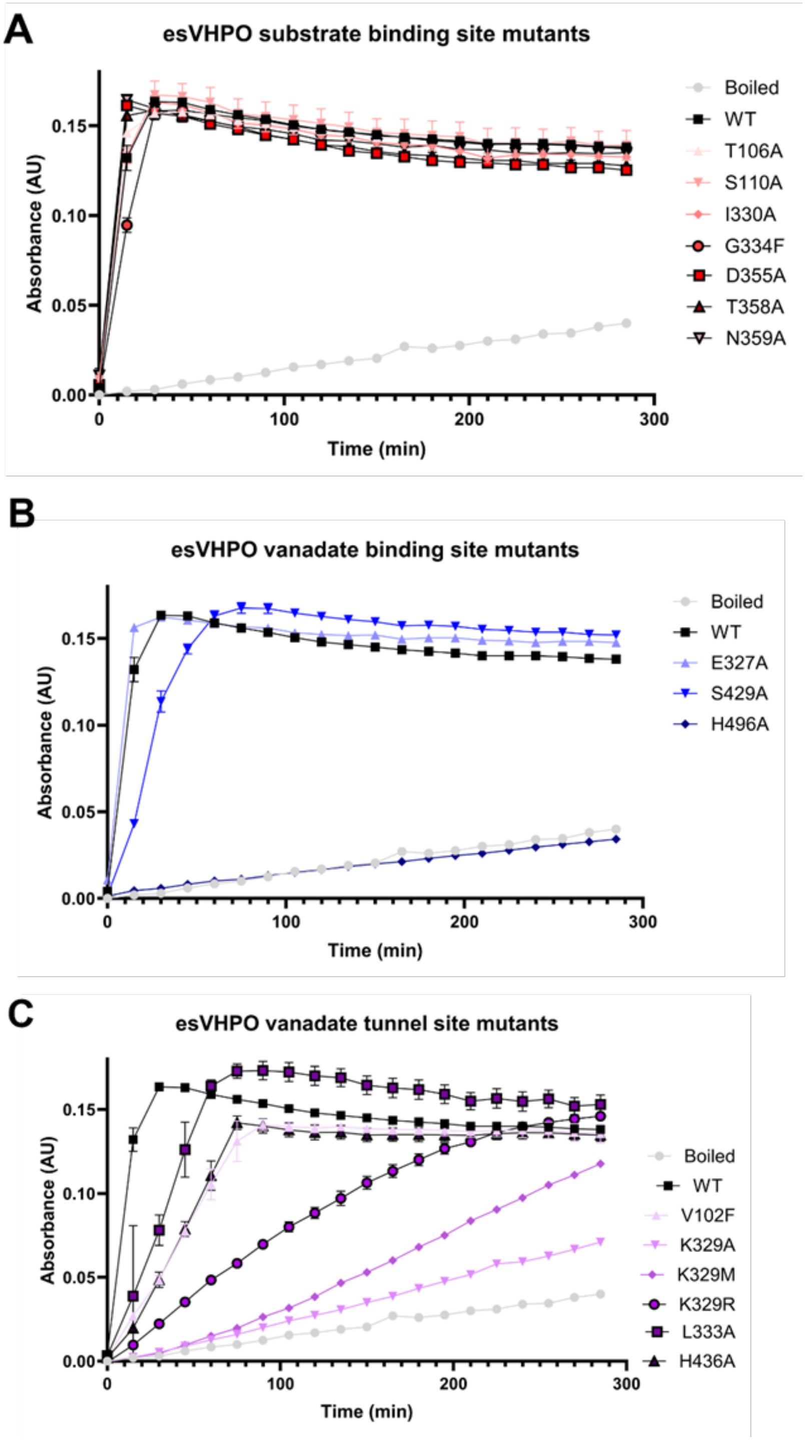

**Figure S9**. **Time course of thymol blue assay esVHPO variants.** Thymol blue time course reactions for esVHPO mutant constructs. The time course curve for boiled esVHPO WT is provided as a negative control and the esVHPO WT curve is included for direct comparison of changes. Boiled-WT and H496A provide background results for the non-enzymatic oxidation of iodide.[6] The three plots are separated by the region of esVHPO they contribute to.

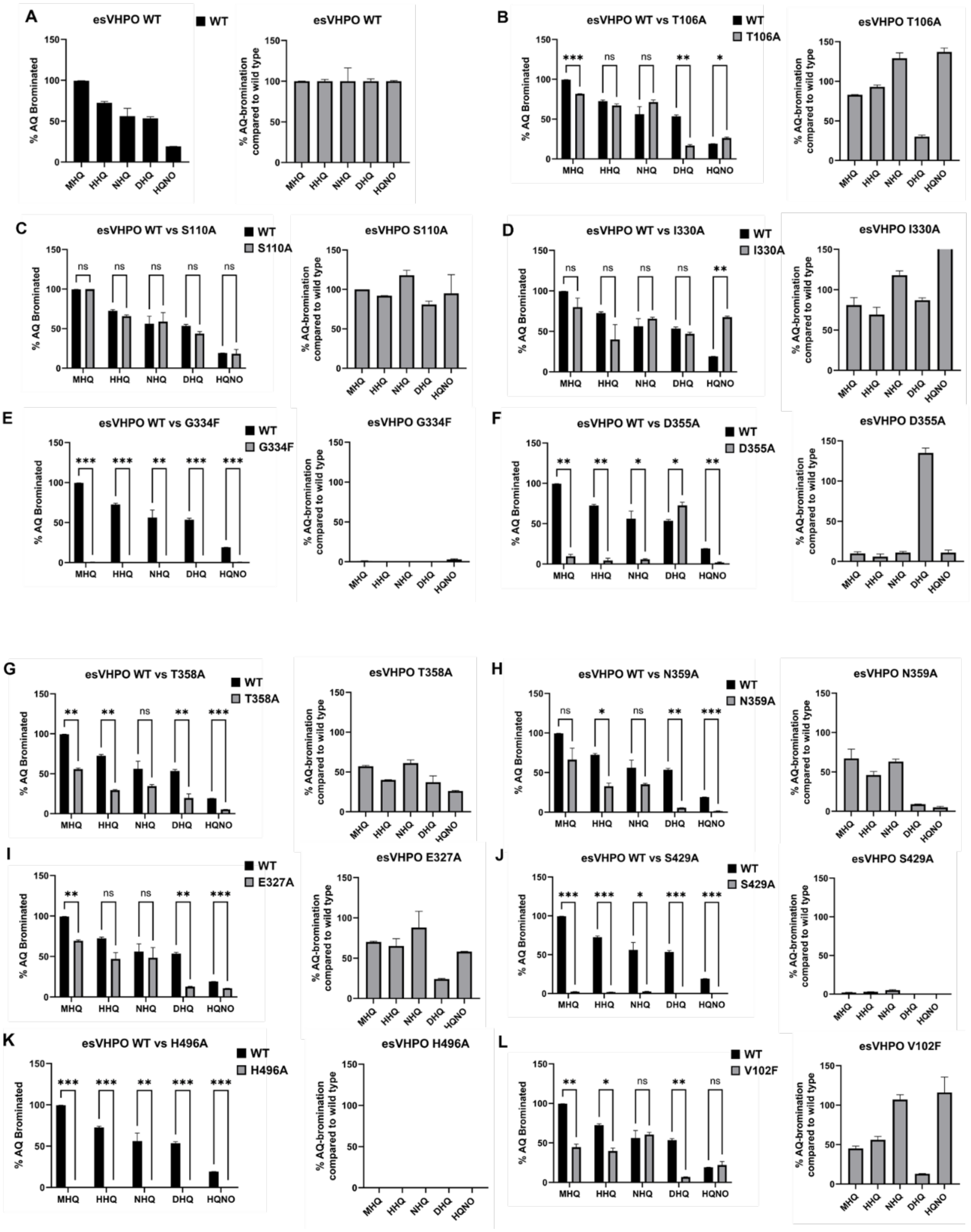
A
esVHPO WT
WT
% AQ Brominated
% AQ-bromination compared to wild type
MHQ
HHQ
NHQ
DHQ
HQNO
B
esVHPO WT vs T106A
T106A
esVHPO T106A
C
esVHPO WT vs S110A
S110A
esVHPO S110A
D
esVHPO WT vs I330A
I330A
esVHPO I330A
E
esVHPO WT vs G334F
G334F
esVHPO G334F
F
esVHPO WT vs D355A
D355A
esVHPO D355A
G
esVHPO WT vs T358A
T358A
esVHPO T358A
H
esVHPO WT vs N359A
N359A
esVHPO N359A
I
E327A
esVHPO E327A
J
esVHPO WT vs S429A
S429A
esVHPO S429A
K
esVHPO WT vs H496A
H496A
esVHPO H496A
L
V102F
esVHPO V102F

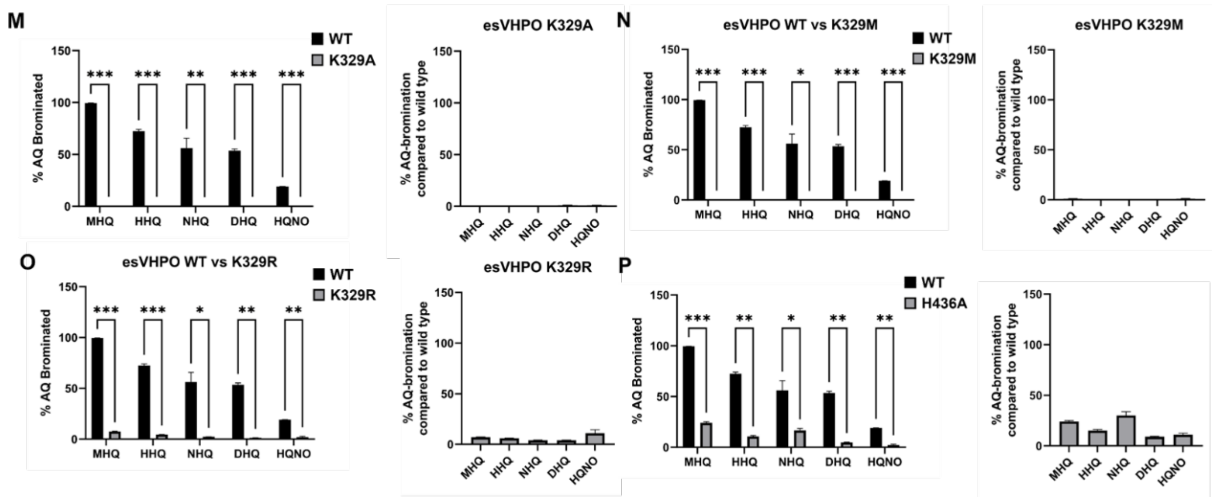

**Figure S10. AQ bromination profile of esVHPO variants compared to wild type**. The AQ bromination activities of esVHPO mutant constructs. For each panel (left), a plot comparing the absolute AQ bromination conversion for each mutant to the WT is shown. Error bars represent the mean ± s.d. (n=3). Significance was calculated with the multiple paired t-test. To judge the significance of the difference between the wild type and mutants the following p-values were used: >0.033 (ns), <0.033 (*), <0.003 (**), <0.001 (***). To the right of each panel is a plot of each mutant's AQ bromination activity normalized to the WT bromination activity of the same substrate.

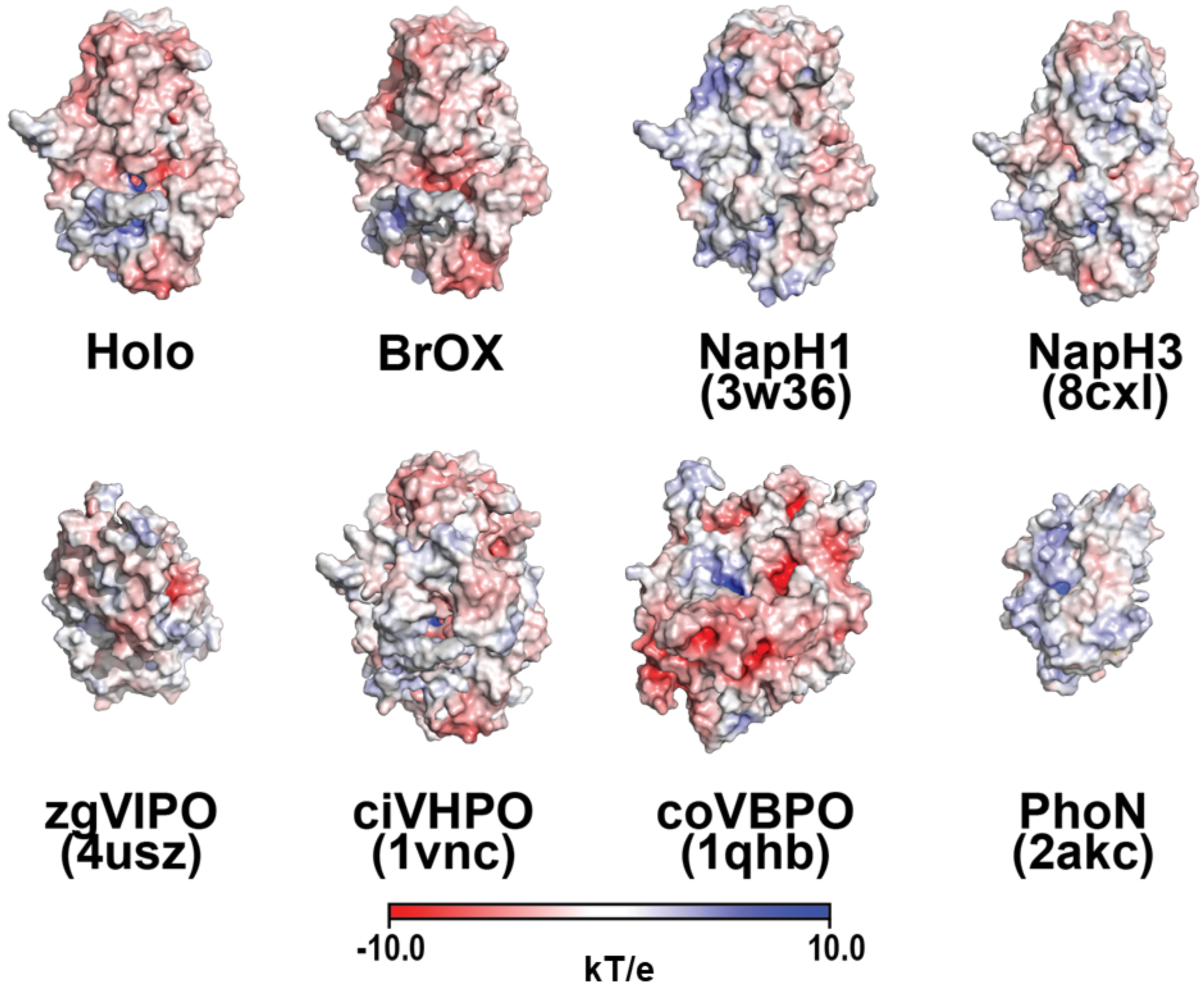

**Figure S11. Electrostatic surfaces of structurally resolved selective and non-selective VHPOs**. Electrostatic surfaces for the monomeric units of the most structurally similar resolved protein structures to esVHPO (see Figure S3). Surface potentials were calculated with the PyMol APBS plug-in. The relative charges are shown as gradient from red (negative) to blue (positive).

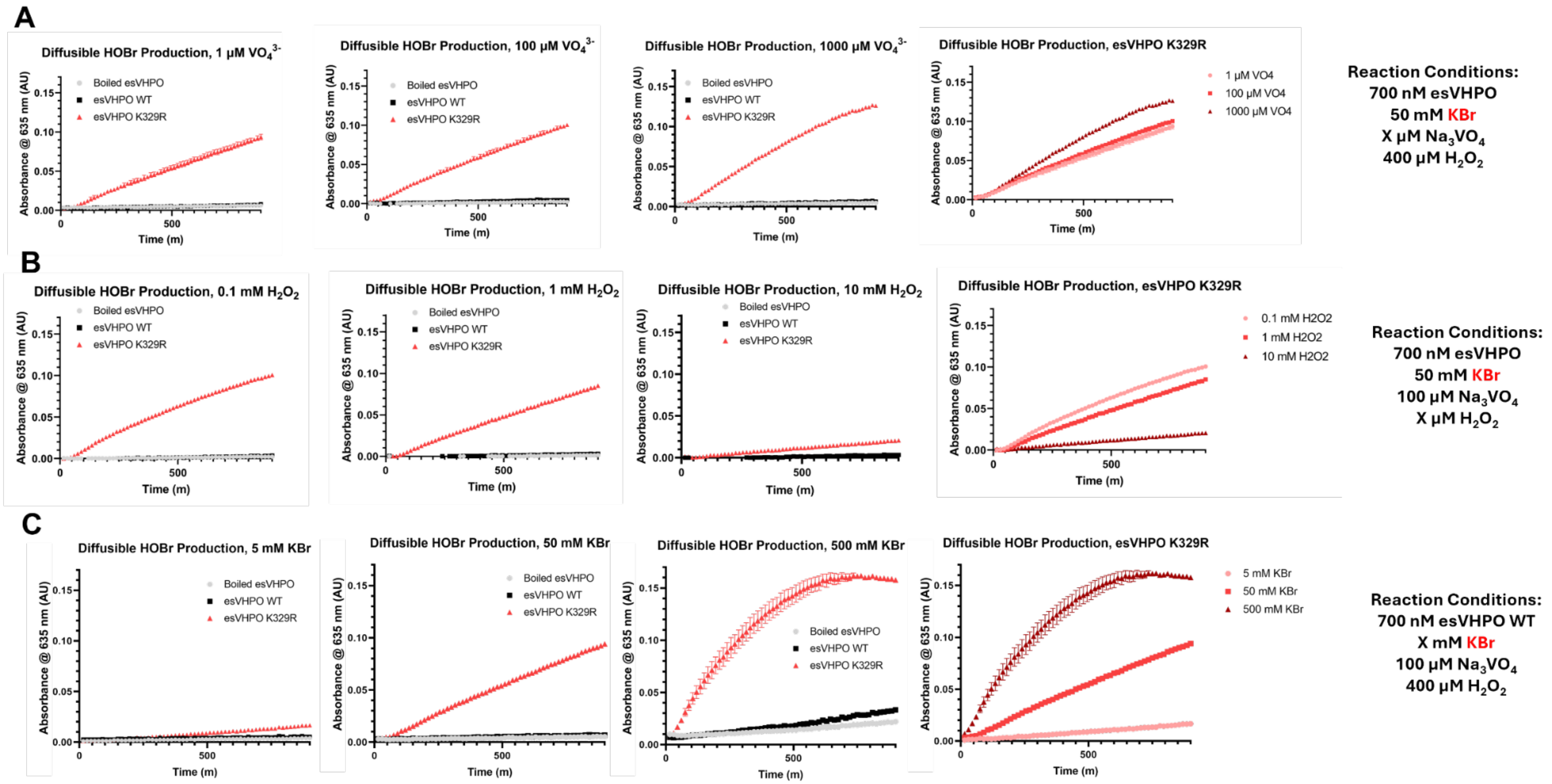

**Figure S12. Substrate dependence of K329R for HOBr production.** TB assay of esVHPO K329R with KBr that shows the production of HOBr unique to this construct. Boiled esVHPO WT (grey) is included as a negative control in all plots. esVHPO WT (black) is included in all plots to show that changing substrate concentrations alone does not generate diffusible HOBr. The fourth column of each panel removes the esVHPO WT curves and summarizes each of the esVHPO K329R conditions. (A) The concentration of added vanadate is altered. (B) The concentration of hydrogen peroxide is altered. (C) The concentration of KBr is altered.

**Table S1. Cryo-EM data collection, refinement and validation statistics**

| | #1 apo (untilted) (EMDB-71844) (PDB 9PTR) | #2 apo (tilted) (EMDB-71844) (PDB 9PTR) | #3 charged (untilted) (EMDB-71845) (PDB 9PTS) | #4 charged (tilted) (EMDB-71845) (PDB 9PTS) |
|---|---|---|---|---|
| **Data collection and processing** | | | | |
| Magnification | 105kx | 105kx | 130kx | 130kx |
| Voltage (kV) | 300 | 300 | 300 | 300 |
| Electron exposure (e–/Å$^2$) | 45.8 | 45.8 | 50 | 50 |
| Defocus range (μm) | -0.8 to -2.8 | -0.8 to -2.8 | -0.8 to -2.8 | -0.8 to -2.8 |
| Pixel size (Å), | 0.835 | 0.835 | 0.6485 | 0.6485 |
| Tilt angle (°) | 0 | 30 | 0 | 30 |
| No. frames | 79 | 79 | 60 | 60 |
| Symmetry imposed | C2 | C2 | C2 | C2 |
| Initial particle images (no.) | 21,142,759 | 6,730,538 | 4,995,816 | 5,172,190 |
| Final particle images (no.) | 1,925,708 | 2,392,787 | 1,933,757 | 1,588,048 |
| Particle images final reconstruction (no.) | 1,249,149 | | 2,144,678 | |
| Map overall resolution (Å) | 2.6 | | 2.4 | |
| FSC threshold | 0.143 | | 0.143 | |
| Map resolution range (Å) | 1.6-14.7 | | 1.7-14.2 | |
| Sphericity | 0.960 | | 0.953 | |
| **Refinement** | | | | |
| Initial model used (PDB code) | Alphafold prediction | | Alphafold prediction | |
| Model resolution (Å) | 2.9 | | 2.8 | |
| FSC threshold | 0.5 | | 0.5 | |
| Model composition | | | | |
| Non-hydrogen atoms | 7186 | | 7161 | |
| Protein residues | 922 | | 918 | |
| Ligands | 2 | | 2 | |
| Water | 24 | | 49 | |
| *B* factors (Å$^2$) | | | | |
| Protein | 118.2 | | 137.4 | |
| Ligand | 166.1 | | 146.7 | |
| R.m.s. deviations | | | | |
| Bond lengths (Å) | 0.03 | | 0.005 | |
| Bond angles (°) | 0.570 | | 0.765 | |
| Validation | | | | |
| MolProbity score | 1.33 | | 1.50 | |
| Clashscore | 1.36 | | 3.03 | |
| Poor rotamers (%) | 2.30 | | 1.77 | |
| Ramachandran plot | | | | |
| Favored (%) | 96.84 | | 96.61 | |
| Allowed (%) | 3.16 | | 3.17 | |
| Disallowed (%) | 0.00 | | 0.22 | |

**Table S2. Primer table**

| Name | Sequence (5'->3') | Description |
|---|---|---|
| esVHPO V102F F | CGCAACGATCTTTCTTCGCTTCACCACGATGCTGAG | Site-directed mutagenesis of E. coli codon optimized N-terminal 6x-His Tagged esVHPO WT in pET28a(+), V102F forward primer |
| esVHPO V102F R | TCGCCCGGTGCCACGCTG | Site-directed mutagenesis of E. coli codon optimized N-terminal 6x-His Tagged esVHPO WT in pET28a(+), V102F reverse primer |
| esVHPO T106A F | ACTTCGCTTCGCAACGATGCTGAGC | Site-directed mutagenesis of E. coli codon optimized N-terminal 6x-His Tagged esVHPO WT in pET28a(+), T106A forward primer |
| esVHPO T106A R | ACGATCGTTGCGTCGCCCGG | Site-directed mutagenesis of E. coli codon optimized N-terminal 6x-His Tagged esVHPO WT in pET28a(+), T106A reverse primer |
| esVHPO S110A F | CACGATGCTGGCAAACGCGTGGTTTG | Site-directed mutagenesis of E. coli codon optimized N-terminal 6x-His Tagged esVHPO WT in pET28a(+), S110A forward primer |
| esVHPO S110A R | GTGAAGCGAAGTACGATCGTTGCGTCGC | Site-directed mutagenesis of E. coli codon optimized N-terminal 6x-His Tagged esVHPO WT in pET28a(+), S110A reverse primer |
| esVHPO E327A F | TGAGCTGTTTGCAGACAAGATCCGC | Site-directed mutagenesis of E. coli codon optimized N-terminal 6x-His Tagged esVHPO WT in pET28a(+), E327A forward primer |
| esVHPO E327A R | GCAGTCACCTTTTGCTCGTCCGT | Site-directed mutagenesis of E. coli codon optimized N-terminal 6x-His Tagged esVHPO WT in pET28a(+), E327A reverse primer |
| esVHPO K329A F | GTTTGAGGACGCGATCCGCAGTC | Site-directed mutagenesis of E. coli codon optimized N-terminal 6x-His Tagged esVHPO WT in pET28a(+), K329A forward primer |
| esVHPO K329A R | AGCTCAGCAGTCACCTTTTGCTCGT | Site-directed mutagenesis of E. coli codon optimized N-terminal 6x-His Tagged esVHPO WT in pET28a(+), K329A reverse primer |
| esVHPO K329M F | GTTTGAGGACATGATCCGCAGTC | Site-directed mutagenesis of E. coli codon optimized N-terminal 6x-His Tagged esVHPO WT in pET28a(+), K329M forward primer |
| esVHPO K329M R | AGCTCAGCAGTCACCTTTTGCTCGT | Site-directed mutagenesis of E. coli codon optimized N-terminal 6x-His Tagged esVHPO WT in pET28a(+), K329M reverse primer |
| esVHPO K329R F | GTTTGAGGACAGGATCCGCAGTC | Site-directed mutagenesis of E. coli codon optimized N-terminal 6x-His Tagged esVHPO WT in pET28a(+), K329R forward primer |
| esVHPO K329R R | AGCTCAGCAGTCACCTTTTGCTCGT | Site-directed mutagenesis of E. coli codon optimized N-terminal 6x-His Tagged esVHPO WT in pET28a(+), K329R reverse primer |
| esVHPO I330A F | TGAGGACAAGGCACGCAGTCTG | Site-directed mutagenesis of E. coli codon optimized N-terminal 6x-His Tagged esVHPO WT in pET28a(+), I330A forward primer |
| esVHPO I330A R | AACAGCTCAGCAGTCACCTTTTGCTC | Site-directed mutagenesis of E. coli codon optimized N-terminal 6x-His Tagged esVHPO WT in pET28a(+), I330A reverse primer |

| | | |
|---|---|---|
| esVHPO L333A F | GATCCGCAGTGCAGGTTTCTCAGC | Site-directed mutagenesis of E. coli codon optimized N-terminal 6x-His Tagged esVHPO WT in pET28a(+), L333A forward primer |
| esVHPO L333A R | TTGTCCTCAAACAGCTCAGCAGTCACC | Site-directed mutagenesis of E. coli codon optimized N-terminal 6x-His Tagged esVHPO WT in pET28a(+), L333A reverse primer |
| esVHPO G334F F | CCGCAGTCTGTTTTTCTCAGCATTATT | Site-directed mutagenesis of E. coli codon optimized N-terminal 6x-His Tagged esVHPO WT in pET28a(+), G334F forward primer |
| esVHPO G334F R | ATCTTGTCCTCAAACAGCTCAGC | Site-directed mutagenesis of E. coli codon optimized N-terminal 6x-His Tagged esVHPO WT in pET28a(+), G334F reverse primer |
| esVHPO D355A F | CGTACACTACGCATTCTTGACCAAC | Site-directed mutagenesis of E. coli codon optimized N-terminal 6x-His Tagged esVHPO WT in pET28a(+), D355A forward primer |
| esVHPO D355A R | AAGTCCAGTAAGCTGTGACCG | Site-directed mutagenesis of E. coli codon optimized N-terminal 6x-His Tagged esVHPO WT in pET28a(+), D355A reverse primer |
| esVHPO T358A F | CGACTTCTTGGCAAACTTGGCCG | Site-directed mutagenesis of E. coli codon optimized N-terminal 6x-His Tagged esVHPO WT in pET28a(+), T358A forward primer |
| esVHPO T358A R | TAGTGTACGAAGTCCAGTAAGCTGTGACC | Site-directed mutagenesis of E. coli codon optimized N-terminal 6x-His Tagged esVHPO WT in pET28a(+), T358A reverse primer |
| esVHPO N359A F | CTTCTTGACCGCATTGGCCGCGTTC | Site-directed mutagenesis of E. coli codon optimized N-terminal 6x-His Tagged esVHPO WT in pET28a(+), N359A forward primer |
| esVHPO N359A R | TCGTAGTGTACGAAGTCCAGTAAGCTGTGACCG | Site-directed mutagenesis of E. coli codon optimized N-terminal 6x-His Tagged esVHPO WT in pET28a(+), N359A reverse primer |
| esVHPO S429A F | CCCGTCGGCAGCAGCAGCGTTTT | Site-directed mutagenesis of E. coli codon optimized N-terminal 6x-His Tagged esVHPO WT in pET28a(+), S429A forward primer |
| esVHPO S429A R | TACTCTGGGTGGTCAGCTACGTCGAGGTATGAGCG | Site-directed mutagenesis of E. coli codon optimized N-terminal 6x-His Tagged esVHPO WT in pET28a(+), S429A reverse primer |
| esVHPO H436A F | TTGCGCAGCGGCAGCCCAGGCAAGTC | Site-directed mutagenesis of E. coli codon optimized N-terminal 6x-His Tagged esVHPO WT in pET28a(+), H436A forward primer |
| esVHPO H436A R | AACGCTGCCGATGCCGACGGGTACTCTGGGT | Site-directed mutagenesis of E. coli codon optimized N-terminal 6x-His Tagged esVHPO WT in pET28a(+), H436A reverse primer |
| esVHPO H496A F | GGGCGGTGTAGCCTTCGAAGACG | Site-directed mutagenesis of E. coli codon optimized N-terminal 6x-His Tagged esVHPO WT in pET28a(+), H496A forward primer |
| esVHPO H496A R | CTCGCTGCGGATACAGCCGTCTTTG | Site-directed mutagenesis of E. coli codon optimized N-terminal 6x-His Tagged esVHPO WT in pET28a(+), H496A reverse primer |

**Table S3. VHPO sequences used for phylogenetic tree construction**

| Accession | Source Species | Name |
|---|---|---|
| **Selective Meroterpenoid VHPOs** | | |
| 3W36 | *Streptomyces* sp. CNQ-525 | NapH1 |
| ABS50492.1 | *Streptomyces* sp. CNQ-525 | NapH4 |
| AGH68909.1 | *Streptomyces* sp. CNH-189 | Mcl24 |
| AGH68925.1 | *Streptomyces* sp. CNH-189 | Mcl40 |
| WP_047018063 | *Streptomyces* sp. CNQ-509 | MarH1 |
| JGI Gene ID: 2802568834 | *Streptomyces* sp. CNZ-289 | LvcH |
| **Selective AQ-VHPOs** | | |
| WP_156035230.1 | *Microbulbifer* sp. Hz11 | HZ11-VHPO |
| WP_076514445.1 | *Oleibacter marinus* | omVHPO |
| WP_052553269.1 | *Enhygromyxa salina* | esVHPO |
| WP_157898942.1 | *Luteitalea pratensis* | lpVHPO |
| WP_128198684.1 | *Rubrivivax albus* | raVHPO |
| WP_132875434.1 | *Tamaricihabitans halophyticus* | thVHPO |
| **Characterized non-selective Bacteria 2 VHPOs** | | |
| 4USZ | *Zobellia galactanivorans* | *Zg*VIPO1 |
| CAZ96406.1 | *Zobellia galactanivorans* | *Zg*VIPO2 |
| WP_011620573.1 | *Synechococcu*s sp. CC9311 | *Ss*VBPO |
| WP_047157019.1 | *Trichodesmium erythraeum* | *Te*VBPO |
| **Uncharacterized non-selective Bacteria 2 VHPOs** | | |
| WP_214610180.1 | *Zobellia barbeyronii* | N/A |
| WP_121850220.1 | *Euzebyella marina* | N/A |
| WP_138657405.1 | *Maribacter algarum* | N/A |
| WP_102183604.1 | *Fischerella thermalis* | N/A |
| KAA2253937.1 | *Solihabitans fulvus* | N/A |
| WP_109251987.1 | *Marinicauda salina* | N/A |
| ERT09389.1 | *Lyngbya aestuarii* BL J | N/A |
| WP_140911141.1 | *Methylomonas koyamae* | N/A |
| WP_087144333.1 | *Crenothrix polyspora* | N/A |
| WP_090316358.1 | *Nitrosomonas oligotropha* | N/A |
| SMD20679.1 | *Kibdelosporangium aridum* | N/A |
| HXA52335.1 | *Solibacter usitatus* | N/A |
| **Characterized non-selective Algal VHPOs** | | |
| 5LPC | *Acaryochloris marina* | *Am*VHPO |
| 5AA6 | *Ascophyllum nodosum* | *An*VHPO |
| 1QHB | *Corallina officinalis* | *Co*VBPO |
| **Uncharacterized non-selective Algal VHPOs** | | |
| AJ491786.1 | *Laminaria digitata* | N/A |
| AAC35279 | *Fucus distichus* | N/A |
| PXF42802.1 | *Gracilariopsis chorda* | N/A |
| XP_005714237.1 | *Chondrus crispus* | N/A |

| | | |
|---|---|---|
| **Characterized non-selective Fungal 1 VHPOs** | | |
| 1VN1 | *Curvularia inaequalis* | *Ci*VCPO |
| HwvCPO | *Hortaea werneckii* UBOCC-A-208029 | *Hwv*CPO |
| **Uncharacterized non-selective Fungal 1 VHPOs** | | |
| CAA72622.1 | *Alternaria didymospora* | N/A |
| XP_003708974.1 | *Pyricularia oryzae* | N/A |
| **Uncharacterized Fungal 2 VHPOs** | | |
| CAA7260301.1 | *Cyclocybe aegerita* | N/A |
| RPA75822.1 | *Ascobolus immersus* | N/A |
| XP_024737652.1 | *Hyaloscypha bicolor* | N/A |
| XP_028466698.1 | *Sodiomyces alkalinus* | N/A |
| **Uncharacterized Rotifera VHPOs** | | |
| CAF1181624.1 | *Didymodactylos carnosus* | N/A |
| UJR11683.1 | *Adineta vaga* | N/A |
| CAF3411370.1 | *Rotaria socialis* | N/A |
| CAF1548122.1 | *Rotaria magnacalcarata* | N/A |
| **Type-II Phosphatidic Acid Phosphatases** | | |
| P26976.1 | *Salmonella enterica* | PhoN |
| P0A924 | *Escherichia coli* | N/A |

### Synthetic Methods

2-methyl-3-iodo-quinolin-4(1H)-one

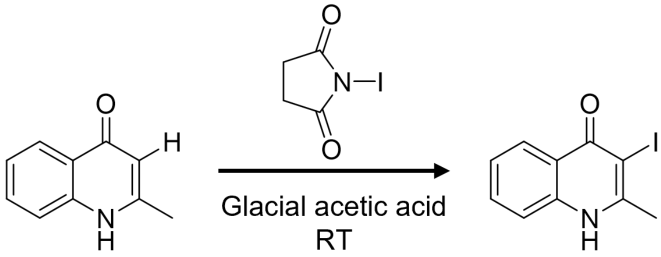

Method was adapted from a previously established protocol for the iodination of 2-heptyl-3-iodo-quinolin-4(1H)-one.N-iodosuccinimide.[7] (0.444 g, 1.98 mmol) was added to a stirred solution of MHQ (0.30 g, 1.89 mmol) in 10 mL glacial acetic acid and stirred at room temperature for four hours. DCM was added to quench the reaction and resulted in the formation of precipitate in the aqueous layer. The aqueous layer was vacuum filtered, rinsed until it was a neutral pH, dried, and collected as an off-white powder (0.249 g, 0.87 mmol, 46% yield).

$^1$H NMR (500 MHz, DMSO-d$_6$) δ 11.97 (s, NH), δ 7.88 (d, *J* = 7.9 Hz, 1H), δ 7.48 (t, *J* = 8.3 Hz, 1H), δ 7.37 (d, *J* = 8.2 Hz, 1H), δ 7.16 (t, *J* = 7.5 Hz, 1H), δ 2.44 (s, 3H). HRMS (MALDI-qQTOF; positive mode; 50:50 CHCA:DHB matrix) Calculated exact mass for $C_{10}H_9INO$ 285.9724 [$(M+H)^+$]. Observed 285.9722 [$(M+H)^+$]

2-methyl-3-deutero-quinolin-4(1H)-one

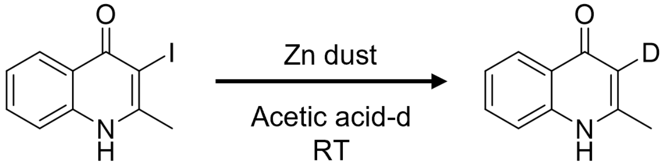

Method was adapted from a previously established protocol for the deuteration of 2-heptyl-3-deutero-quinolin-4(1H)-one.[7] 2-methyl-3-iodo-quinolin-4(1H)-one (0.188 g, 0.66 mmol) and zinc dust (0.05 g, 3.4 mmol) were added to a flame dried round-bottom flask. The flask was evacuated and refilled with argon three times in a dry Schlenk tube. Monodeuteroacetic acid (15 ml), which had previously been degassed by three freeze-pump-thaw cycles and stored under argon, was added to the mixture and stirred at room temperature under argon overnight. Potassium carbonate (9 g) was dissolved in $D_2O$ (15 mL), added to the reaction, and stirred under argon for 10 minutes to quench the reaction. DCM (50 mL) was added to dissolve the resulting precipitate and the aqueous layer was filtered to remove remaining unwanted solids. The filtered aqueous layer was extracted again with DCM (2 x 50 mL). All organic layers were combined, dried over $MgSO_4$, and dried *in vacuo* to give a white powder (0.049 g, 0.31 mmol, 47%). 90% deuteration was indicated by the integration of the H-3 singlet in $^1$H NMR [δ 6.34 (s, 0.09 H)].

$^1$H NMR (500 MHz, $CDCl_3$) δ 8.38 (d, *J* = 8.0 Hz, 1H), δ 7.74 (d, *J* = 8.4 Hz, 1H), δ 7.63 (d, *J* = 7.8 Hz, 1H), δ 7.41 (t, *J* = 7.7 Hz, 1H), δ 2.42 (s, 3H). HRMS (MALDI-qQTOF; positive mode; 50:50 CHCA:DHB matrix) Calculated exact mass for $C_{10}H_9DNO$ 161.0820 [$(M+H)^+$]. Observed 161.0813 [(M+H)

N-N-dimethyl-1-amino-5-naphthalenesulfinic acid (dansyl sulfinic acid)

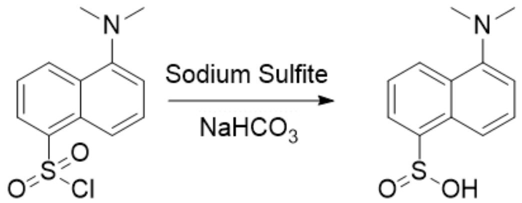

Dansyl sulfinic acid ($DANSO_2H$) was synthesized following a previously published protocol.[8] Dansyl chloride (1.35 g, 5.0 mmol) was mixed with sodium sulfite (2.52 g, 20 mmol), sodium bicarbonate (1.68 g, 20 mmol), and 10 mL water. The reaction was stirred at 70 °C for 2.5 hours. Concentrated sulfuric acid was added to the reaction until precipitate stopped forming. The solid was collected with vacuum filtration and washed with DI water until the pH was neutral. The resulting solid was further dried *in vacuo*. The dried precipitate was then resuspended in 2.8 M sodium hydroxide. Sulfuric acid was added to the solution until no more precipitate formed (roughly pH 4). The precipitate was vacuum filtered and washed with DI water until the pH was neutral. Resulting solid was further dried *in vacuo*, resulting in a white powder (0.34 g, 1.44 mmol, 29% yield). By NMR, the product was about 85% pure and by mass spec the major contaminant was dansyl sulfonic acid; notably all dansyl chloride was fully reacted.

$^1$H NMR (500 MHz, DMSO-$d_6$) δ 8.89 (d, $J$ = 8.6 Hz, 1H), 8.22 (d, $J$ = 8.6 Hz, 1H), 8.06 (d, $J$ = 7.0 Hz, 1H), 7.74-7.57 (m, 3H), 3.17 (s, 6H). HRMS (MALDI) exact mass calculated for $C_{12}H_{13}NO_2S$ 235.0667 [$(M+H)^+$], found 235.0663 [$(M+H)^+$].

**NMR Spectra**

2-methyl-3-iodo-4(*1H*)-quinolinone (I-MHQ) spectra

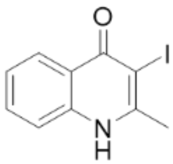

**I-MHQ**
500 MHz $^1$H
in $d_6$-DMSO

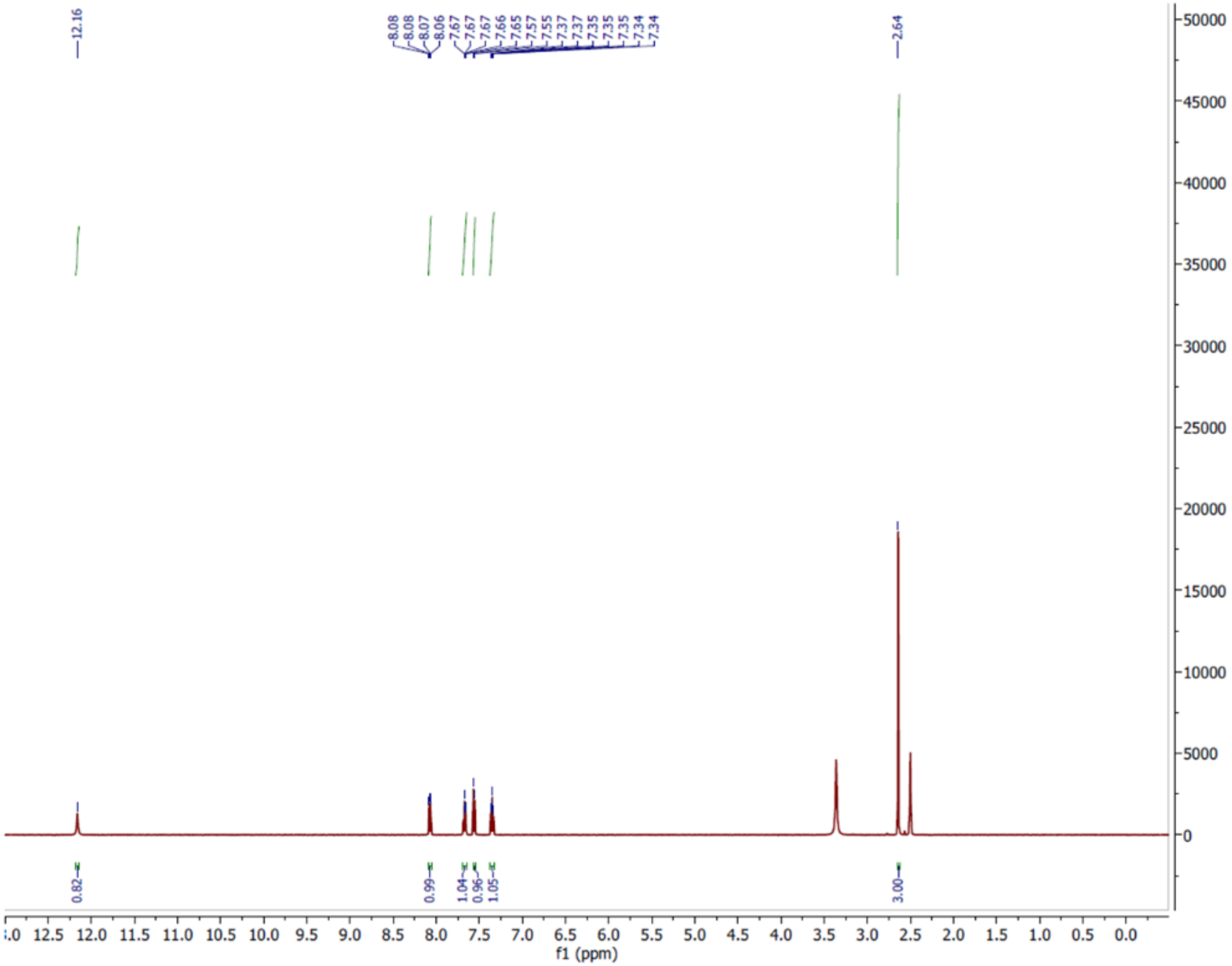

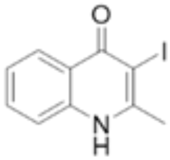

**I-MHQ**
500 MHz COSY
in $d_6$-DMSO

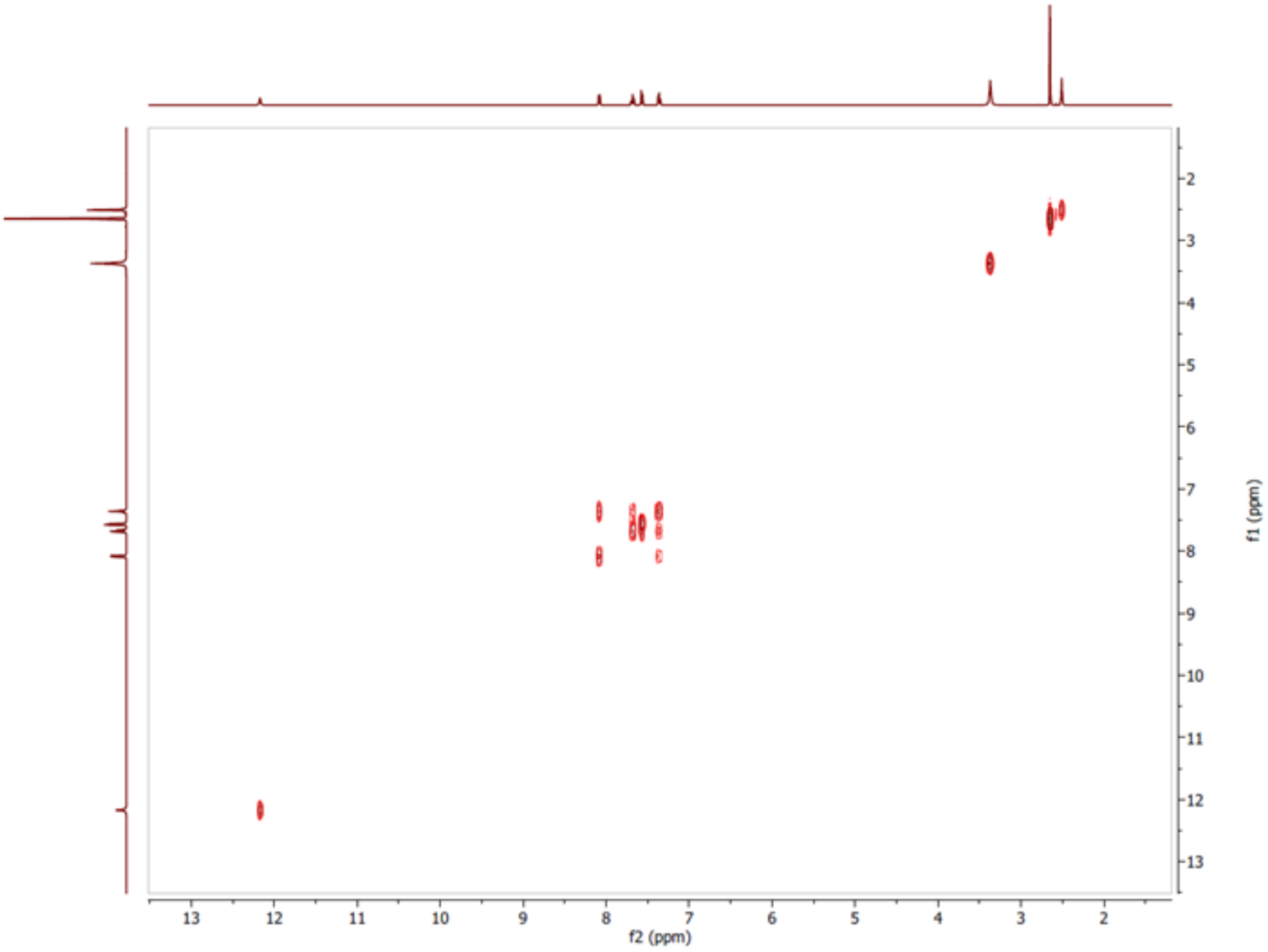

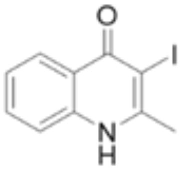

**I-MHQ**
500 MHz HSQC
in $d_6$-DMSO

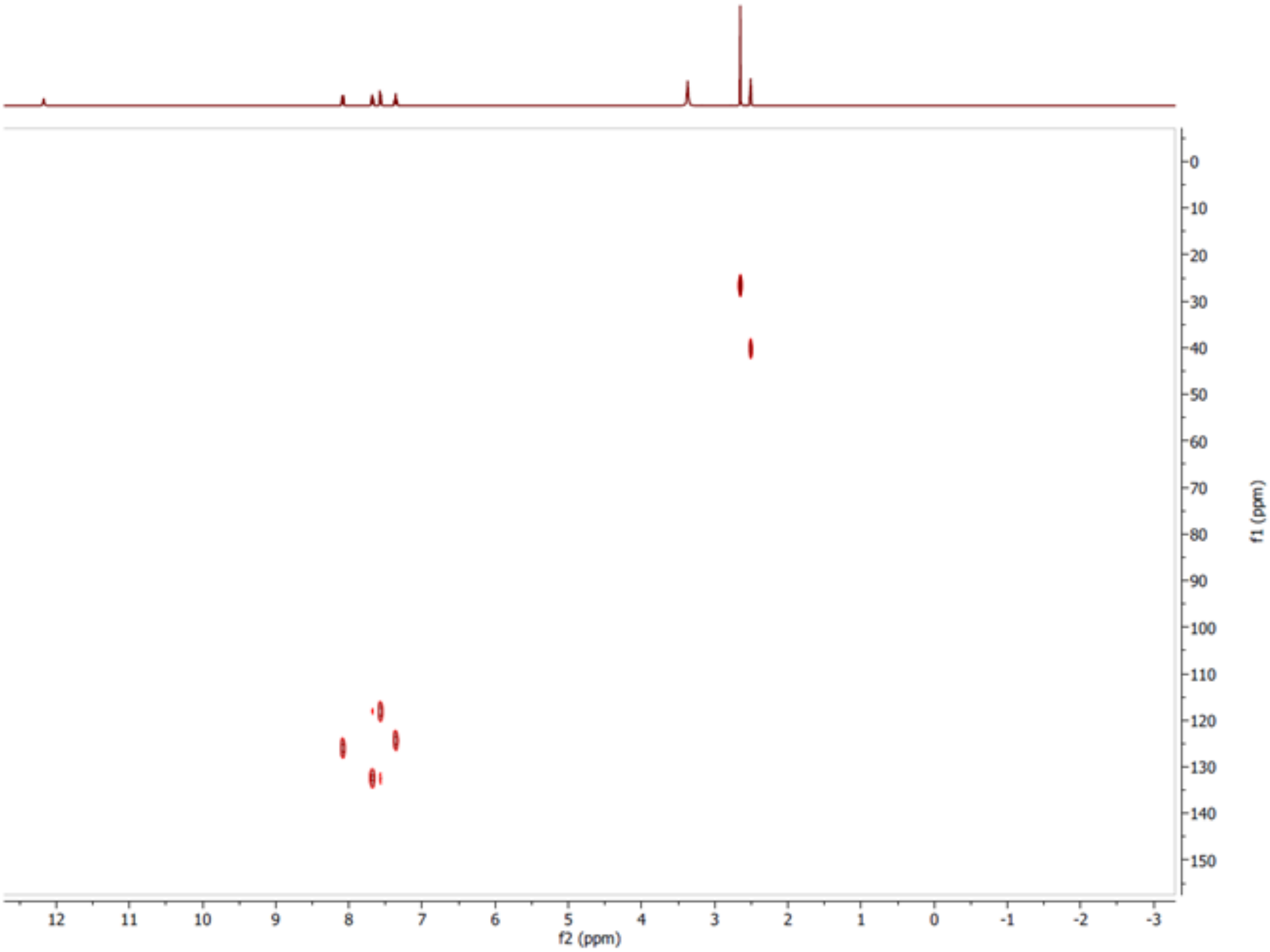

2-methyl-3-deutero-4(*1H*)-quinolinone (D-MHQ) spectra

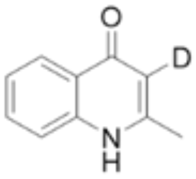

**D-MHQ**

500 MHz $^{1}H$
in $CD_3OD$

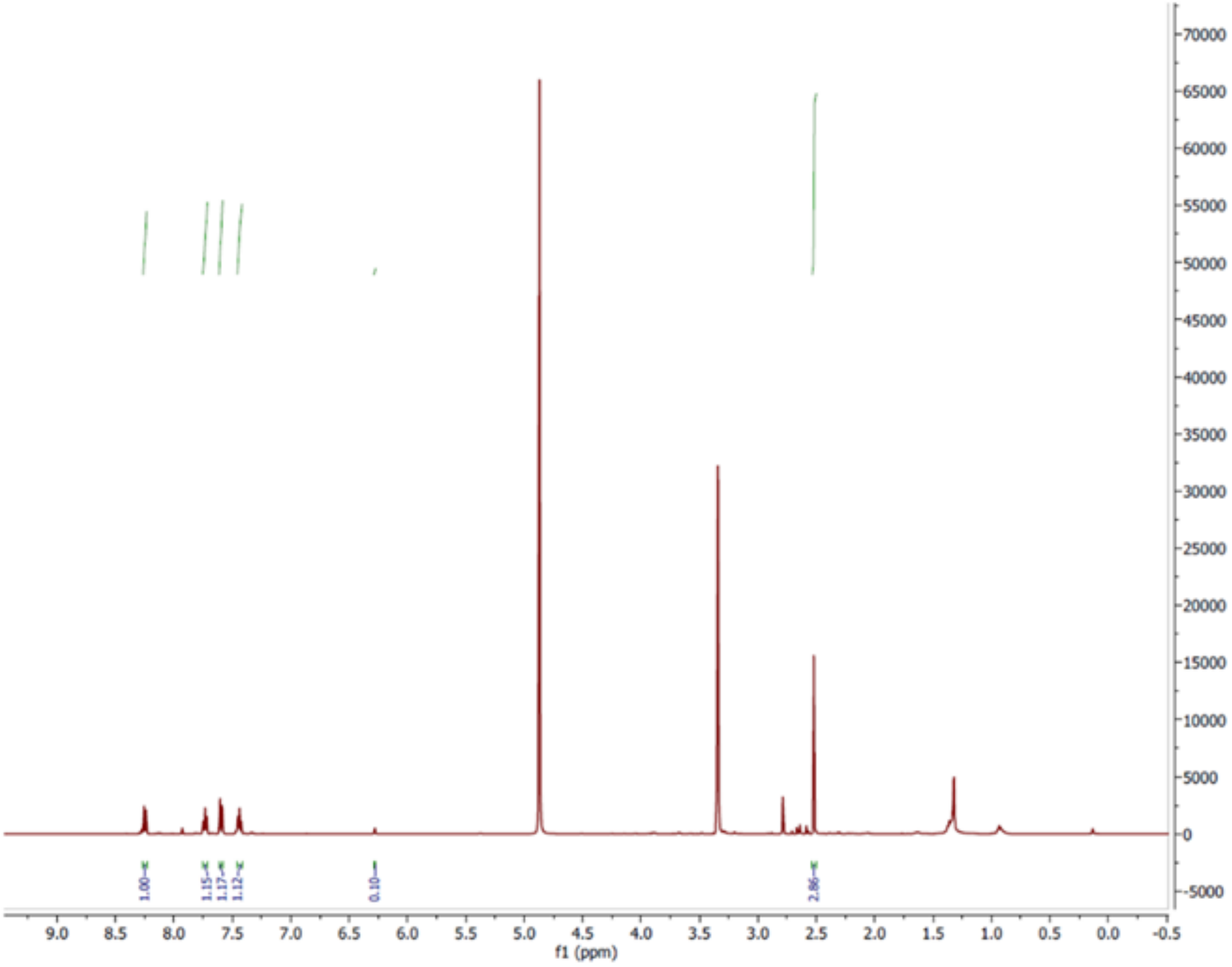

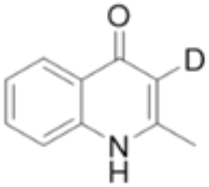

**D-MHQ**

125 MHz $^{13}C$
in $CD_3OD$

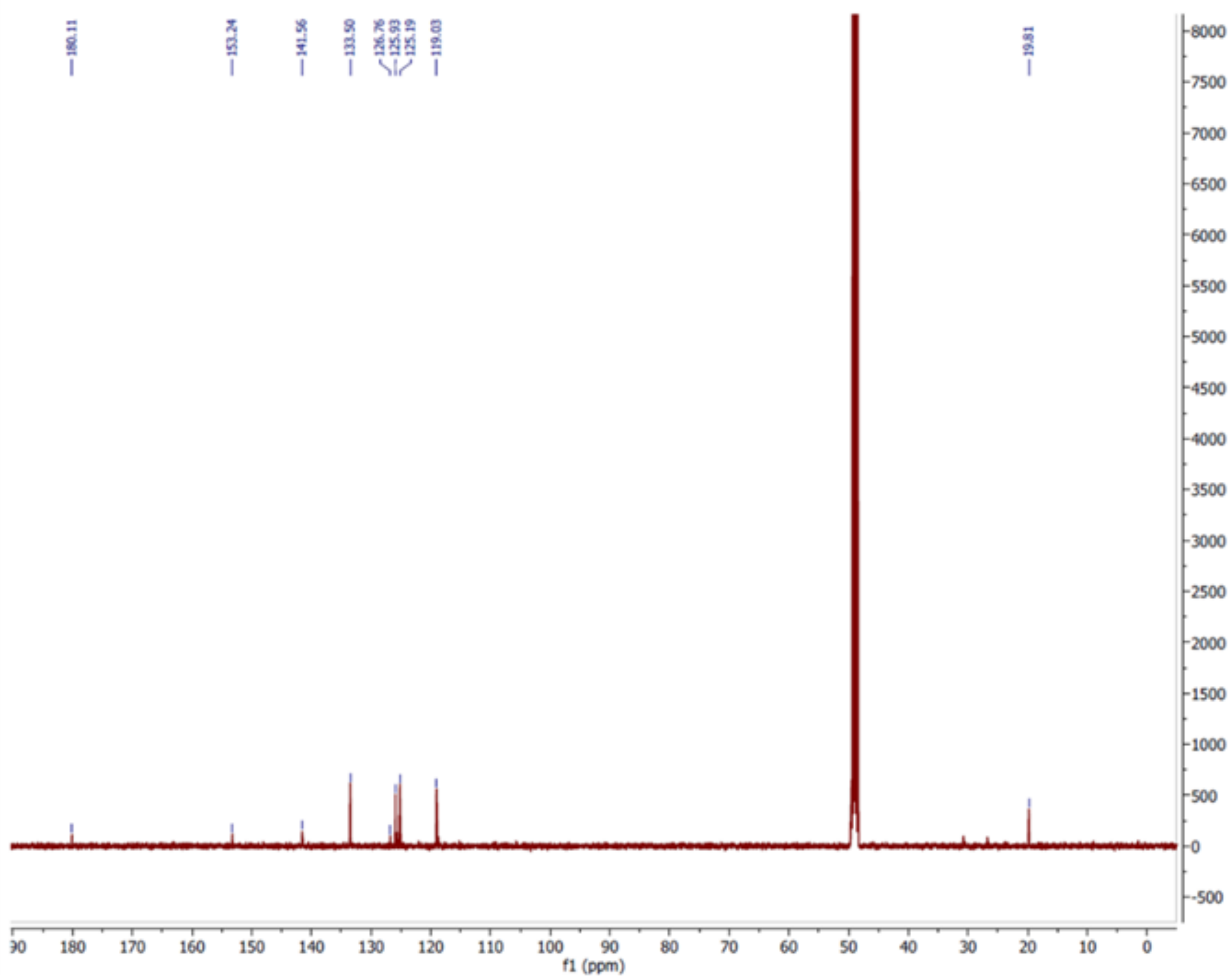

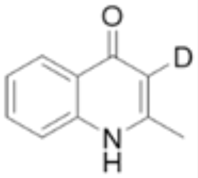

**D-MHQ**

500 MHz HSQC
in $CD_3OD$

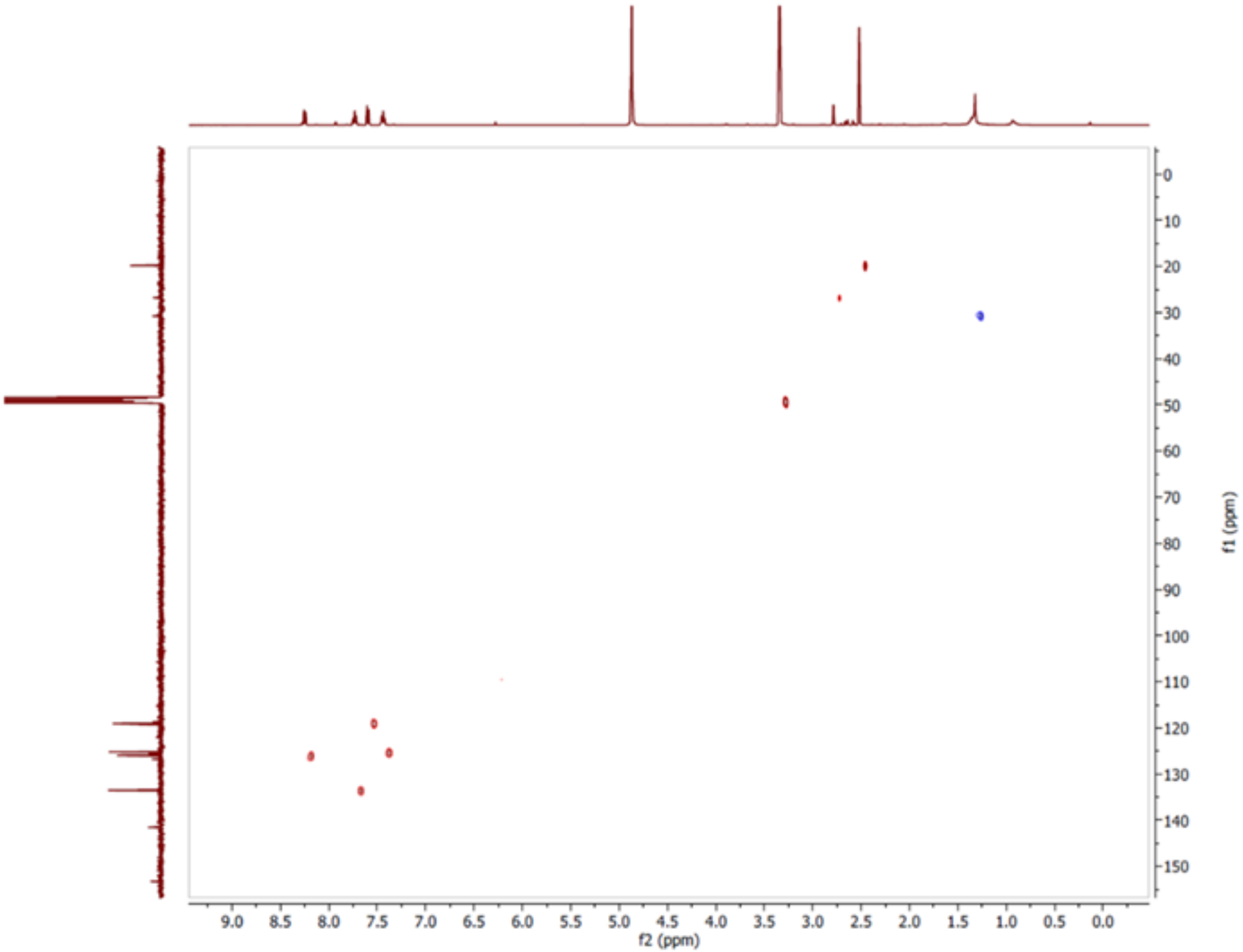

Dansyl sulfinic acid spectra

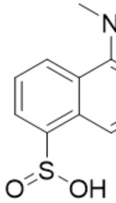

**DANSO$_2$H**
500 MHz $^1$HNMR
in $d_6$-DMSO

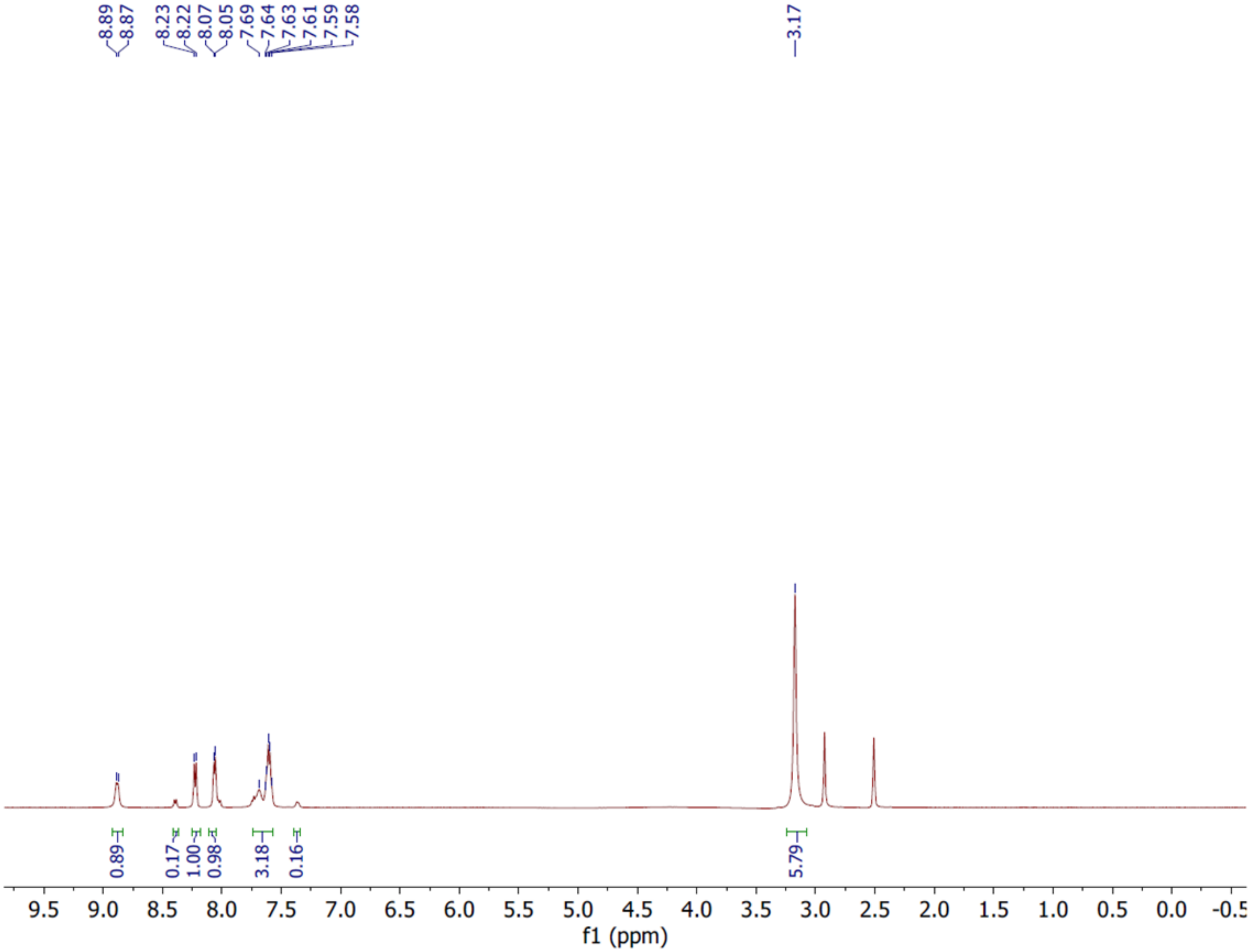

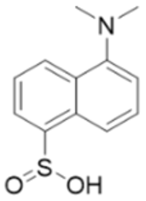

**DANSO$_2$H**
500 MHz COSY
in $d_6$-DMSO

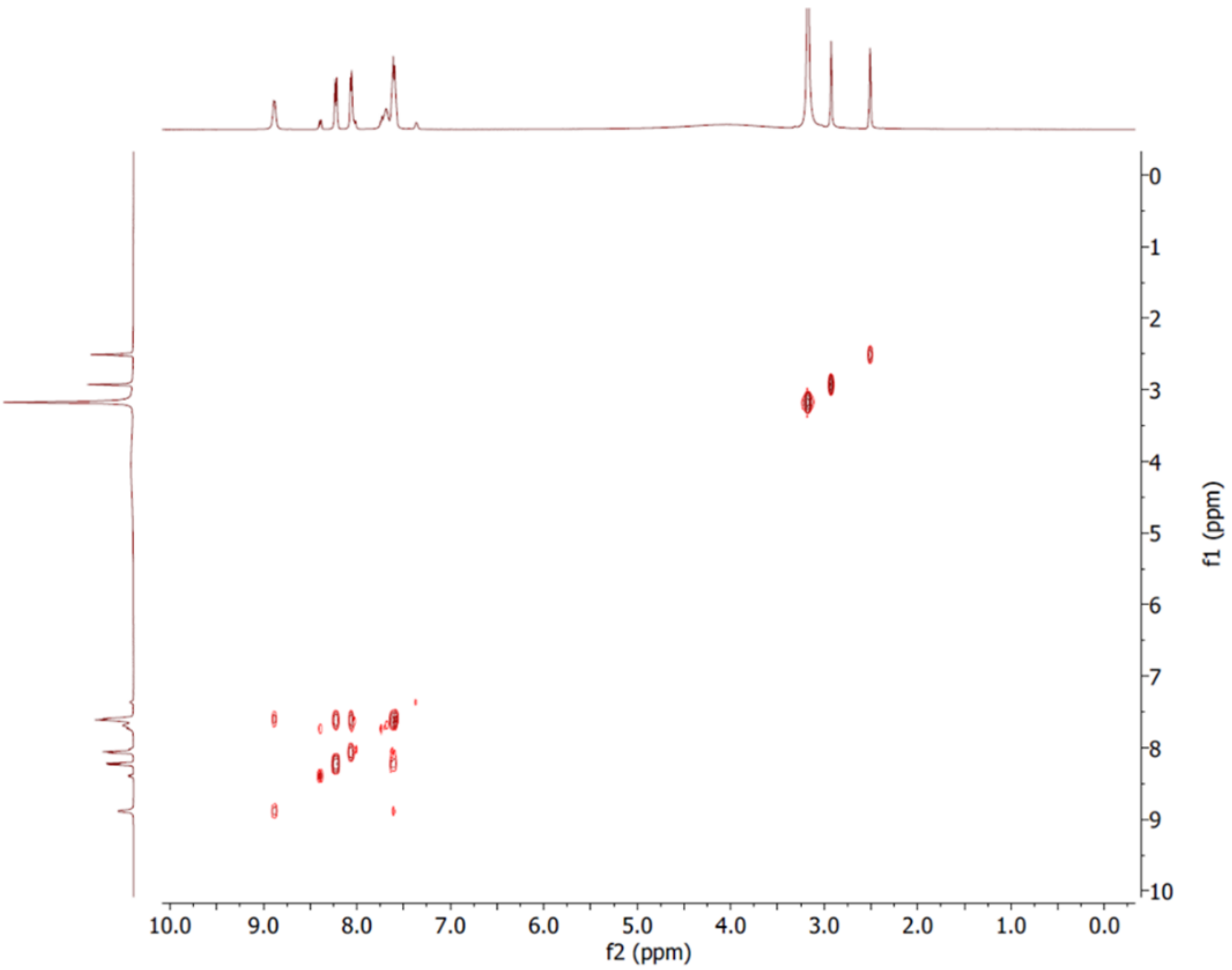

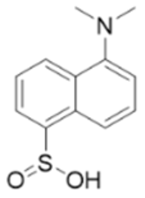

**DANSO$_2$H**
500 MHz HSQC
in $d_6$-DMSO

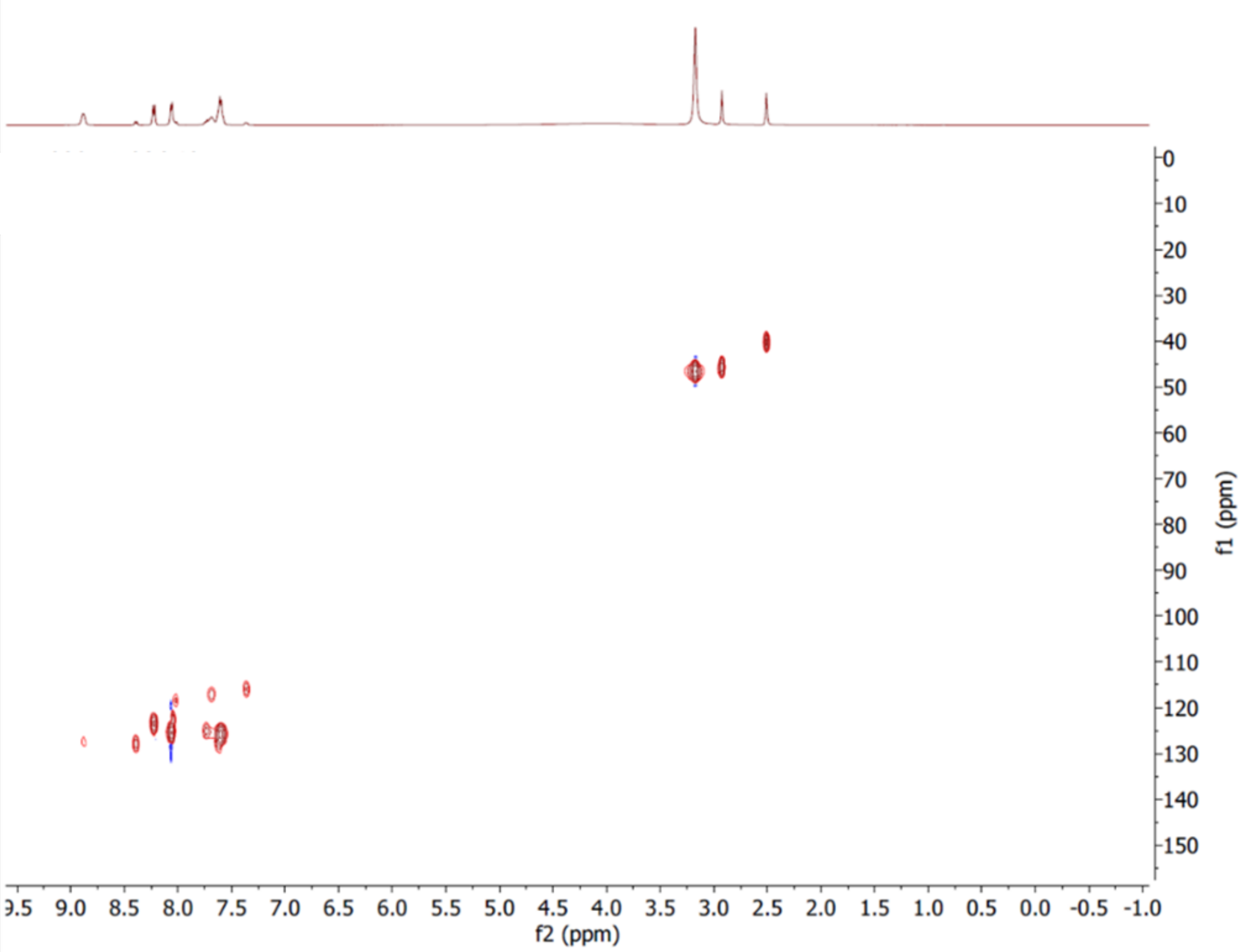
